# A mechanical pacemaker sets rhythmic nephron formation in the kidney

**DOI:** 10.1101/2023.11.21.568157

**Authors:** Samuel H. Grindel, Sachin N. Davis, Igor Guranovic, John M. Viola, Aria Huang, Kenji Yeoh, Michael J. Yelpo, Rachel S. Kim, Grace Y. Liu, Jiageng Liu, Kate Bennett, Grace Qian, Catherine M. Porter, Arushi E. Sahay, Lele Song, Jonathan Levinsohn, Andrew P. McMahon, Nils O. Lindström, Liling Wan, Kotaro Sasaki, Katalin Susztak, Alex J. Hughes

## Abstract

The developing mammalian kidney exponentially duplicates nephron-forming stem cell niches at the tips of the urinary collecting duct tree to achieve massively parallel function. Nephron formation rate has a clinically meaningful effect on person-to-person variability in nephron endowment^1–5^, while exerting *in vitro* control could enable sustained waves of nephrogenesis in organoid-derived synthetic kidney tissues^6,7^. However, how the kidney arrives at an appropriate number and ratio of nephrons to collecting ducts is unclear. Here we show that nephron formation is rhythmic and synchronized with branching of the ureteric bud tree (the future urinary collecting ducts). We correlate human and mouse spatial transcriptomics data with the branching ‘life-cycle’ to uncover rhythmically alternating signatures of nephron progenitor differentiation and renewal. The nephron progenitor rhythm parallels rhythmic nuclear elongation and other hallmarks of mechanical tension in surrounding stromal cells that we attribute to branching-induced deformation. Stroma-specific knockdown of actomyosin activity leads to a striking loss of synchronization between nephron formation and ureteric bud branching without blocking either. These results suggest that the stroma acts as a mechanically entrained pacemaker for nephron formation. Together, our findings uncover a feedback mechanism for clock-like coordination of organ composition during exponential growth.

## Introduction

Periodic patterning in space achieves parallelization of tissue function, such as in hair follicles^8^, intestinal villi^9,10^, and digits^11^. Periodic patterning in time allows tissues to accommodate moving reference frames caused by growth. One example is the clock and wavefront model^12^ thought to govern formation of somites, the future vertebrae. Here a triggering clock interacts with a traveling wave that shifts the zone of competence for cells to condense into somites along an elongating body axis. The kidney is a radical example in which nephrons are continuously patterned over embryonic day E12.5 to postnatal day 4 in the mouse, during which the organ grows in volume by >1,000-fold^13^. With embryonic organ formation lying beyond the reach of intravital imaging in mammals, very little is known about how repetitive patterning (much less organ-wide tissue composition) is maintained during such extreme organ growth.

Collectives of nephron progenitor cells cap tips of the branching ureteric bud (UB) tree during kidney development, forming traveling stem cell niches just below the kidney surface in an inductive front known as the nephrogenic zone (**Fig. 1A**). These caps contribute cells that drop behind the front and condense into pre-tubular aggregates (PTAs) — the earliest nephron stage. Only a fraction of a cap’s progenitors commit to each successive nephron, enabling niches to maintain persistent progenitor pools during development^13^. Nephron progenitor (‘cap mesenchyme’) populations are thought to balance renewal vs. differentiation by interpreting autonomous cues and those from adjacent UB, early nephron, and stromal cell populations^14–18^. FGF/EGF/MAPK^18–23^, Wnt/β-catenin^17,24–26^, Hippo/Yap^27–33^, BMP/TGFβ^34–36^, and Notch^37–39^ pathways are involved here, among others. However, this picture of a consistent balance conflicts with the observation that cells commit in avalanche-like streams from the cap mesenchyme into discrete PTAs rather than continuously^40^. This suggests that either 1) cell differentiation rate is constant in time; cells accumulate in the niche until a suitable site^41^ or other unknown trigger^42^ for condensation arises, or 2) cell differentiation rate varies rhythmically, pushing a fraction of progenitors into an irreversibly differentiated state at the same time. Such a rhythm could be autonomous or arise from feedback associated with the dominant rhythmic feature of kidney development - the UB branching life-cycle.

**Fig. 1:**
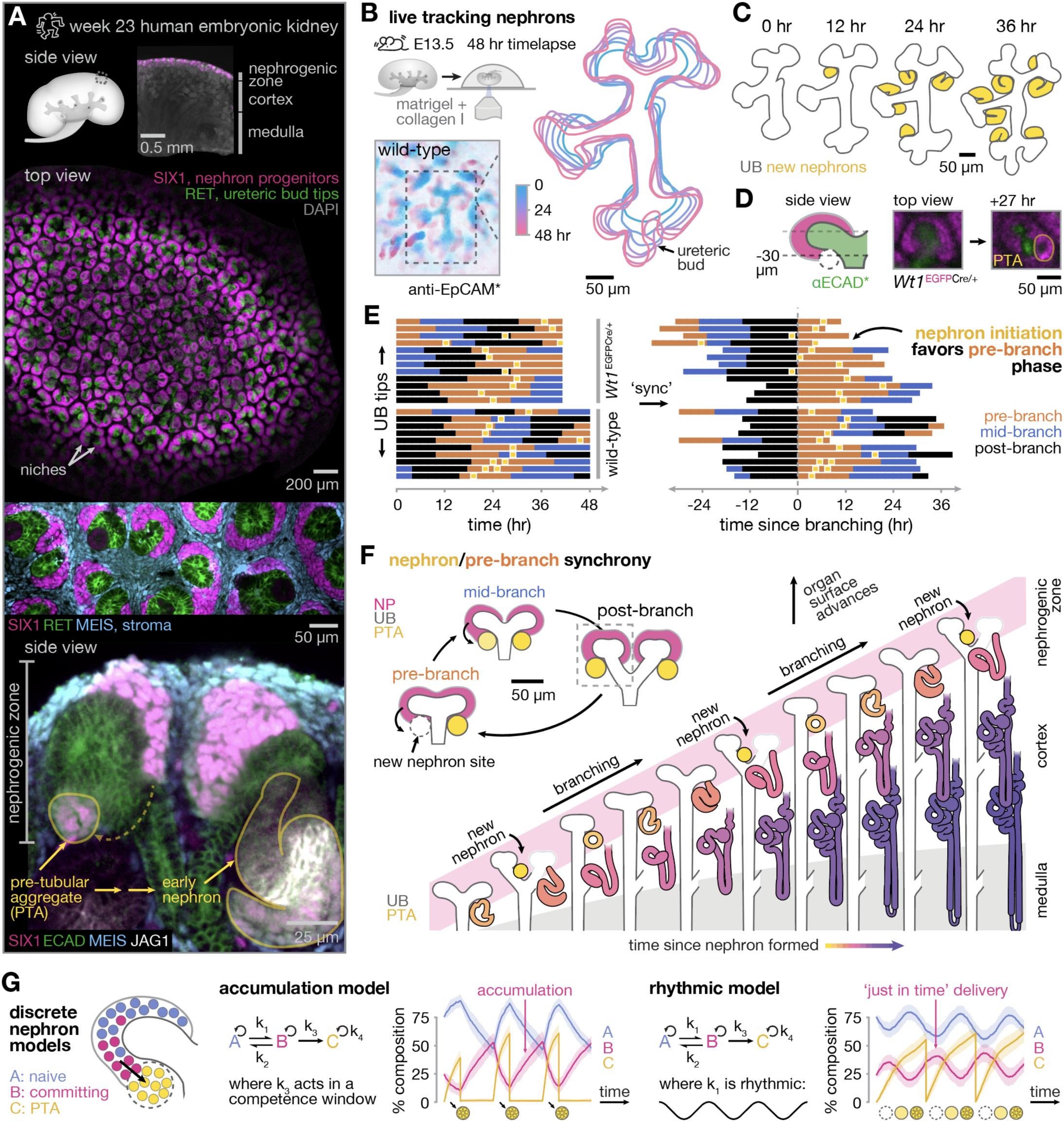
Condensation of new mouse nephrons is synchronized with branching of the ureteric bud epithelial tree. (**A**) Human week 23 kidney immunofluorescence images showing cell compartments constituting the nephrogenic niche. Each ECAD+ RET+ ureteric bud tip s surrounded by SIX1+ (human) / SIX2+ (mouse) nephron progenitors (the cap mesenchyme) and bordered by MEIS1/2/3+ PDGFRA+ stroma. (**B**) *Top-left*, Schematic of live E13.5 mouse embryonic kidney 3D explant culture in matrigel/collagen 1 hydrogel. *Bottom-right*, time-coded immunofluorescence and segmented projections of the ureteric bud. (**C**) Manual segmentation of nascent nephrons based on emergent EpCAM staining. (**D**) *Top*, Sketch and confocal immunofluorescence images of a representative niche in Wt1^EGFPCre/+^ kidney over 27 hr explant culture, showing nascent nephron condensation as a WT1+ cell cluster beneath ureteric bud tip. (**E**) Ticker-tape plots of nephron formation events relative to ureteric bud tip branching stage (see Fig. 2) against real time (*left*), and after aligning each tip to a common time since branching initiation; *n* = 3 kidneys each for wild-type and Wt1^EGFPCre/+^. (**F**) *Left*, Model of rhythmic contribution of committing nephron progenitors (NP) to condensation sites newly formed by ureteric bud (UB) branching morphogenesis. *Right*, Model of hythmic nephron formation in tissue context, consistent with mouse kidneys up to E19.5 (Short *et al.*, 2014) and ‘period one’ human kidneys up to week 15 (Osathanondh & Potter, 1963). (**G**) 3-state stochastic simulations of nephron progenitor sub-population composition in the niche, where naive and differentiating / committing cells are in equilibrium. ‘Accumulation model’: Committing cells accumulate in the niche until allowed to segregate into pretubular aggregates (PTAs) within periodic competence windows. ‘Rhythmic model’: Committing cells contribute continuously to PTAs, but are formed from naive cells with a rhythmic differentiation rate k_1_. In both models, all cell states proliferate at rate k_4_ and PTA cells leave the niche at the start of each branching cycle.

Here we organize spatial transcriptomic, protein marker, and biophysical data across ∼10^3^ human and mouse niches by the branching stages of their associated ureteric bud tips. We find rhythmic transcriptional changes in both nephron progenitors and the immediately adjacent stroma. Nephron progenitor commitment and condensation synchronize with the early phase of each UB branching life-cycle, interspersed with periods in which nephron progenitors are biased toward renewal. Seeking an underlying clock, we show that the nephron progenitor rhythm is coordinated with stromal cell entry, differentiation, and exit from the niche along the corticomedullary (shallow-to-deep) axis. Live explant cultures show that proper synchronization of nephrogenesis with ureteric bud branching is dependent on stromal tension and differentiation. These data establish a rhythmic principle of nephrogenic niche regulation. Committing cells are delivered ‘just in time’ to nephron condensation sites newly formed by branching, ensuring a consistent balance between tissue compartments during rapid organ growth.

## Results

### Mouse nephrons form rhythmically at the initiation of ureteric bud tip branching through fractional commitment of progenitors

We hypothesized that an unknown pace-maker synchronizes nephron formation with ureteric bud branching because of two observations. First, nephron formation initiates in the curved ‘armpits’/‘angles’ formed as UB tips begin to branch (whichever is not already occupied)^13,41–47^. Second, the rates of nephron formation and UB branching in the mouse embryonic kidney are equivalent from E15-E19.5 (ref. ^13^). However, synchrony between the two has not been formally quantified. We tackled this using a newly-developed explant culture system that preserves much of the native 3D structure of the mouse kidney^48^ (**Fig. 1B**). Taking the reference frames of individual UB tips, we annotated the timing of transitions between branching stages (pre-, mid-, and post-branching) and the earliest points at which nephron condensation events were perceptible local to tips.

Branch staging was based on semi-quantitative morphology features (described later in **Fig. 2**). Nephron condensation was scored in wild-type kidneys using a live EpCAM antibody label for the UB that also stains newly forming distal tubules/connecting segments (**Fig. 1C**, **Movie S1**). New nephrons were similarly scored in *Wt1*^EGFPCre/+^ embryos in which all condensing nephron progenitors are labeled^49–51^ (**Fig. 1D**, **Movie S2**). Where at least one nephron formed, 21 of 25 events occurred in the pre-branching stage as the ureteric bud tip elongates from a symmetrical ‘ampulla’ to a pill-shaped ‘T-tip’ bearing new armpit sites (wild-type: 10/12, 83%; *Wt1*^EGFPCre/+^: 11/13, 85%), **Fig. 1E**. These data verify that nephron formation is rhythmic and synchronized with the ureteric bud tip branching life-cycle (**Fig. 1F**, **Movie S3A**). We do not use ‘rhythmic’ to imply that branching period for any given tip is consistent over time, that branching is synchronized among different tips, or that a nephron always forms when a site is available.

**Fig. 2:**
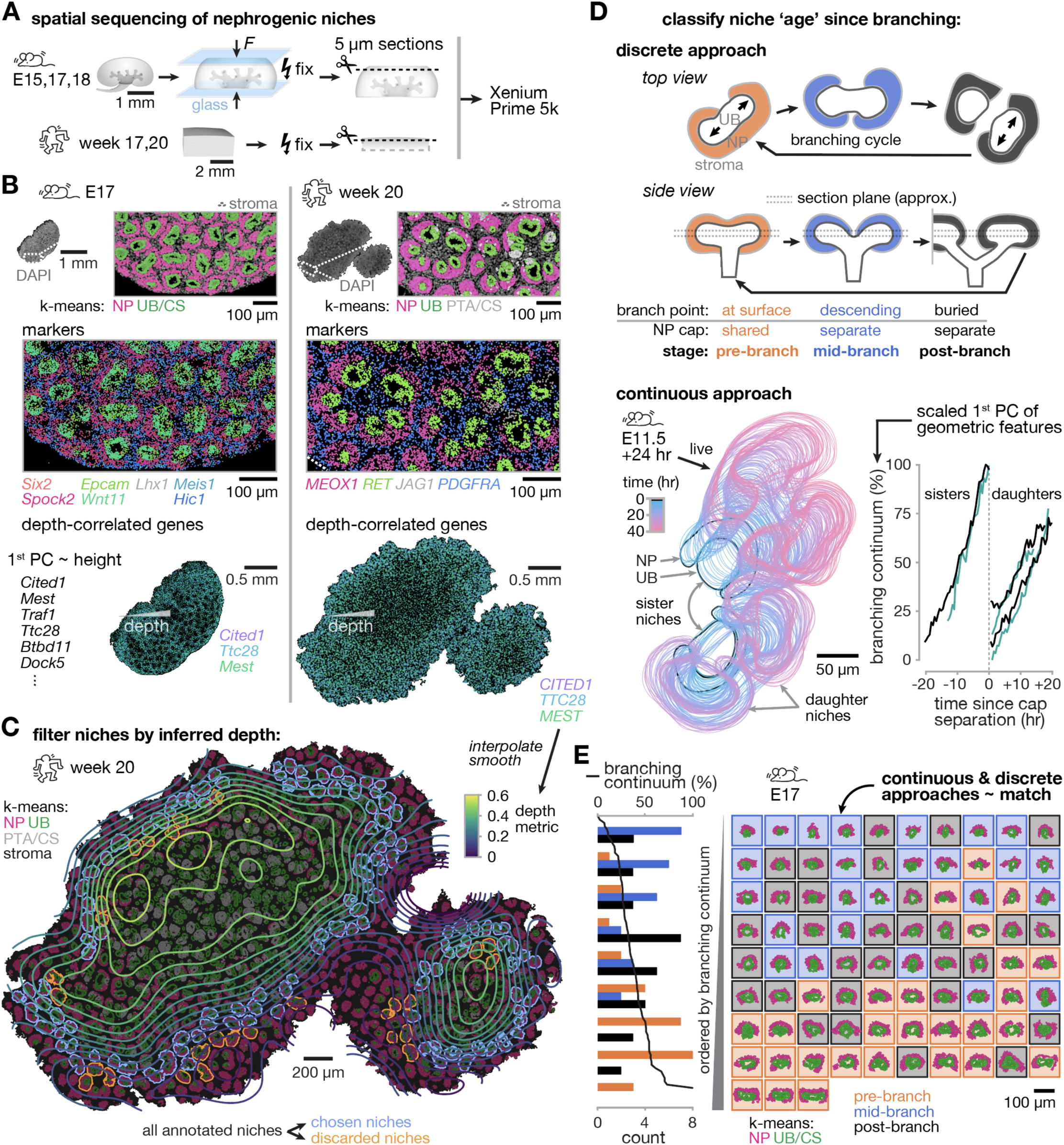
Registering spatial transcriptomes of nephron progenitor niches to the branching life-cycle. (**A**) Schematic of kidney sectioning for spatial sequencing. (**B**) *Top*, Example mouse and human kidney slices showing overlays of DAPI fluorescence and cells detected by Xenium spatial sequencing and colored by k-means cluster. *Middle*, Marker transcripts for cap mesenchyme (nephron progenitors, NP, ∼magenta), ureteric bud (UB, ∼green), early nephrons (gray), and stroma (∼blue) detected by Xenium. PTA/CS: pretubular aggregate/connecting segment. *Bottom*, Example depth-correlated transcripts in the first principle component of those variable among niches. (**C**) Contours of the depth metric overlaid on k-means Xenium view, highlighting ‘chosen’ niches sharing a similar anatomical position in depth. (**D**) *Top*, Schematic of qualitative life-cycle metrics (branch point position in *z* and cap mesenchyme connectivity) used to score each niche as pre-, mid-, or post-branching. *Bottom*, Outlines of ureteric bud and cap mesenchyme compartments in an E12.5 mouse kidney explant over a 40 hr confocal fluorescence time-lapse and plot of a continuous branching state score (‘branching continuum’) relative to real time of cap mesenchyme niche splitting during branching. (**E**) Histogram of discrete branching stage counts per row of a matrix of niches ordered left-to-right and top-to-bottom by increasing branching continuum. Niche outlines are colored by discrete branching stage. All nephron progenitor and ureteric bud cells associated with each niche are shown, colored by Xenium k-means cluster.

### Placing niche transcriptomes into the reference frame of the ureteric bud branching life-cycle to enrich for correlations with nephron progenitor state

At least two models for rhythmic, discrete nephron formation are consistent with current understanding (**Fig. 1G**). To establish predictions, we used stochastic simulation of dynamics among three nephron progenitor states^42^ — A) naive cells in equilibrium with B) differentiating/committing cells; and C) cells irreversibly sequestered within PTAs. Committing nephron progenitors could be mustered by accumulation in the niche if PTA formation only occurs during a rhythmic competence window^41^. Alternatively, committing cells could be delivered ‘just in time’ to PTAs through rhythmic differentiation from the naive state. Both models predict that a cyclical transcriptional state of nephron progenitors would manifest over the branching life-cycle, albeit with distinct predictions for the phase relationship between the two.

Measuring this required us to 1) sequence nephron progenitors from separate niches, and 2) assign niches to branching stages before comparing gene expression between them. We performed Xenium Prime 5k, which returns spatial positions of single transcripts at sub-cellular resolution for ∼5,100 genes of interest in 5 µm-thick tissue sections. We ran duplicate Xenium assays on human week 17 and 20 and partially flattened mouse E15, 17, and 18 embryonic kidneys (**Fig. 2A**, **Table S1**; **Fig. S1** presents HCR RNA-smFISH marker validation). We began with a pseudobulk analysis of gene expression in nephron progenitors (**Fig. S2**) and their nearest-neighbor stromal and ureteric bud cells. For this reason, we use a broad definition of ‘stroma’ in this paper that includes all cells in the ‘ribbons’ separating nephron progenitor niches. Starting with nephron progenitors, we found that the first principle component of variable genes among niches was dominated by those having a gradient in expression with tissue depth (**Fig. 2B**). This reflects a known naive-to-committing gradient in nephron progenitor identity along the corticomedullary axis^34,45^. We filtered niches using a ‘depth metric’ created from several height-correlated genes (*CITED1*, *TTC28*, *MEST*) to retain only those in a roughly comparable anatomical zone (**Fig. 2C, Fig. S3A for mouse**). We next assigned discrete ureteric bud branching stages to niches using branch point position and cap mesenchyme connectivity as metrics (**Fig. 2D**). Seeking a continuous estimate, we created a ‘branching continuum’ score; the scaled first principle component of a set of geometric features of each niche. The branching continuum scaled linearly with real time for a mouse kidney explant culture timelapse and ordered niches similarly to their discrete branching stages (**Fig. 2D,E**, **Movie S4; Fig. S4 for human**). These data reflect successful categorization of niches along a UB branching life-cycle continuum and exclusion of depth-related effects.

### Nephron progenitor and stromal transcriptomes reveal rhythmic renewal and differentiation waves synchronized with the ureteric bud branching life cycle

We next sought to detect transcriptional differences within niche cell compartments between pre-, mid-, and post-branching niches. Unsupervised hierarchical clustering sorted niches roughly by these categories (**Fig. 3A**, **Fig. S3B**, **Figs. S5-S13**). Known markers^17,25,26,35,40,42,43,52–57^ in blocks of clustered variable genes exposed a rhythmic transition in nephron progenitors between a differentiation/proliferation state earlier and a naive state later in the UB branching life-cycle. We validated that several markers of these states are rhythmic by whole-mount immunofluorescence in E17 mouse kidneys (**Fig. S14, Note S1**). Comparing E17 slices at different depths indicated that the nephron progenitor rhythm takes on a naive vs. proliferation signature shallower in the niche and a naive vs. proliferation + differentiation signature deeper in the niche (compare **Fig. S9B** and **Fig. S10B**). This adds a temporal rhythm to published accounts of a correlation between depth, differentiation, and proliferation^13,43^. A similar rhythm in stroma appeared between a cortical nephrogenic stroma (‘shallow’) state and a medullary nephrogenic stroma (‘deep’) state. These categories mirror a known differentiation gradient set up by a retrograde ‘conveyor belt’ of cells into deeper tissue layers relative to branching niches^53,54,58–60^, though a connection to the branching life-cycle was not previously known. While a rhythm emerged in the ureteric bud, it primarily reflected the rhythmic inclusion of differentiating nephron progenitors as they fuse (anastomose) with the bud at newly formed connecting segments^43,61^. These data are evidence that the transcriptional state of niches is not monolithic over the ureteric bud branching cycle. Rather, mouse and human nephron and stromal progenitors change rhythmically in pathways relevant to renewal vs. differentiation decision-making.

**Fig. 3:**
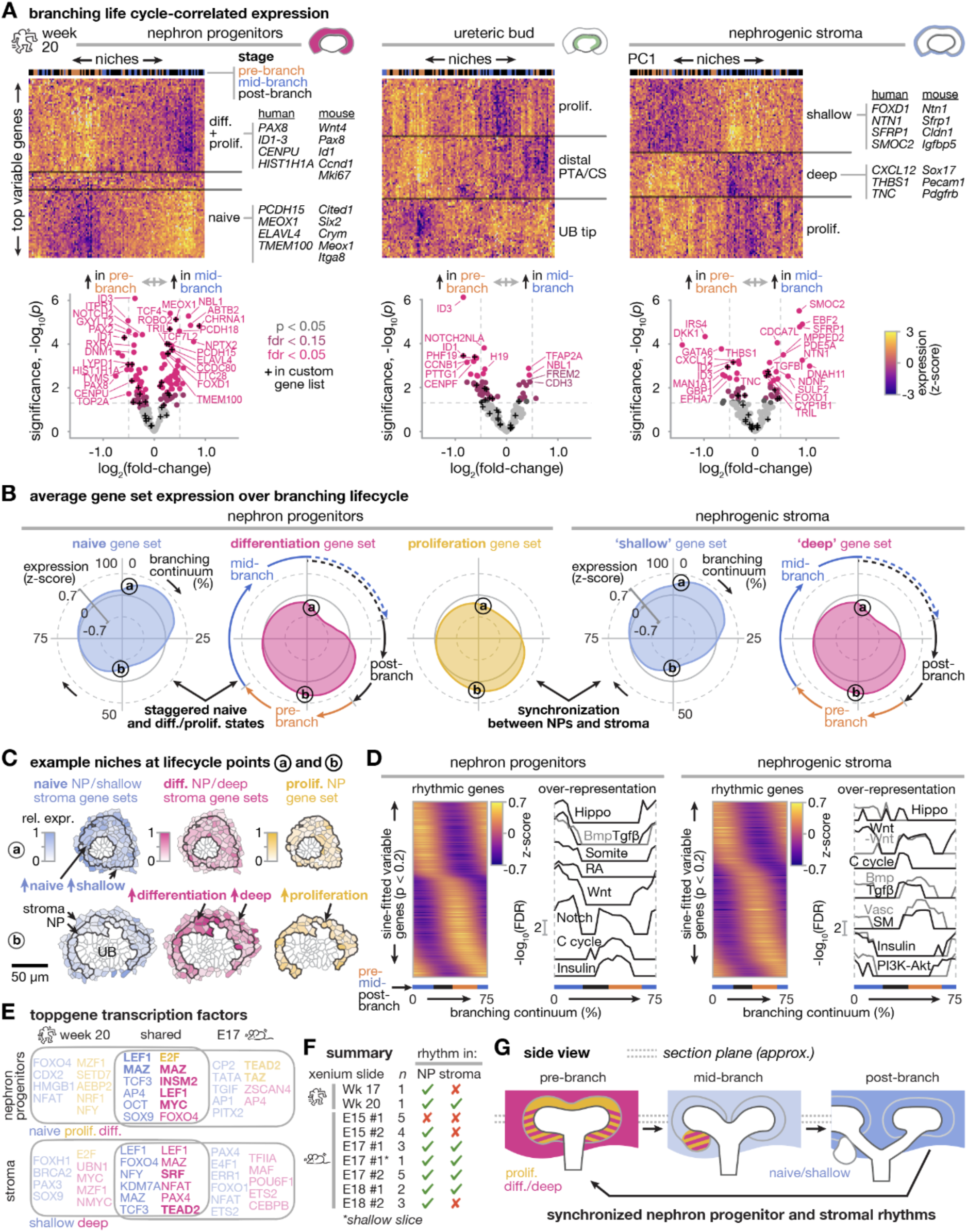
Nephron progenitor and stromal compartments of the human nephron-forming niche have differentiation rhythms that synchronize with each other and the ureteric bud branching life-cycle. (**A**) Unsupervised hierarchical clustering heatmaps (top 100 variable genes, *top*) and differential expression plots (top 200 variable genes, *bottom*) for pseudobulked transcript expression across niches at different branching lifecycle stages (*n* = 120 niches in a human week 20 tissue slice, see **Fig. S5** for full analysis). Blocks of genes are qualitatively annotated based on preponderance of genes previously associated with the indicated cell states. As part of Xenium analysis, we pre-selected 91 custom genes (of the 5092 total) known to be relevant to nephrogenic niche development, which are indicated with crosses. (**B**) Radial plots of mean z-scores for gene sets differentially expressed in (A) and qualitatively associated with the indicated cell states for nephron progenitor and stromal cell compartments. Radial distance from the center is expression level, angular distance clockwise from the 12 o’clock position indicates branching continuum. (a) and (b) labels correspond to rows in (C). (**C**) Heatmaps of gene set expression for representative niches at single-cell resolution in renewal (a) and differentiation (b) phases. (**D**) Heatmap of expression vs. branching continuum for rhythmic genes ordered by peak position fitting a sine wave model at p < 0.2 and line plots of GO/KEGG/reactome analyses performed using a sliding window of width 25 branching continuum units over the same genes (Hippo, Hippo signaling pathway/multiple species (stroma); Bmp, BMP signaling pathway; Tgfβ, signaling by TGFB family members; Somite, somite development; RA, response to retinoic acid; Wnt, canonical Wnt signaling pathway; -Wnt, negative regulation of Wnt signaling pathway; Notch, notch signaling pathway; C cycle, cell cycle; Insulin, insulin signaling pathway / receptor signaling pathway/cascade (stroma); Vasc, kidney vascular development; SM, smooth muscle cell differentiation; PI3K-Akt, PI3K signaling / PI3K-Akt signaling pathway). (**E**) Notable ToppGene transcription factor binding sites unique and shared among week 20 human and E17 mouse kidney tissue, colored by qualitative cell state. (**F**) Qualitative summary of transcriptional rhythm detection among all experiments (see Figs. S5-S13 for full analyses). (**G**) Schematic model of rhythmic transcriptional changes in nephron progenitor and stromal compartments over branching lifecycle.

Representing scores created from differentially expressed genes in radial plots show that the differentiation state of nephron progenitors is synchronized with the deep differentiating state of stromal cells, peaking during new branch initiation (**Fig. 3B,C; Fig. S3C,D**). 29% and 24% of detectable genes in the nephron progenitor and stromal compartments, respectively, fit a sine-wave expression rhythm when human week 20 niches were ordered along the branching continuum. This revealed wave-like up-and down-regulation of genes at phase-shifts spanning the branching life-cycle (**Fig. 3D, Fig. S3E**).

The genes with greatest fit to the rhythmic sine-wave model were enriched for renewal/differentiation markers in nephron progenitors (top two: STXBP5, NBL1 / PAX2, NOTCH2 respectively), and shallow/deep markers in stroma (top two: CDCA7L, FOXD1 / DKK1, ID3 respectively). A sliding-window GO/KEGG/reactome gene set over-representation analysis revealed transcriptional rhythms in several pathways relevant to nephron progenitor renewal and differentiation, notably Wnt^17,24–26^, Hippo^27–33^, retinoic acid (RA)^62–64^, BMP/TGFβ^34–36^, Notch^37–39^, and insulin^65–67^ signaling (**Fig. 3D, Fig. S3E**). The stromal pathways showed some overlap with these, plus renal vasculature and smooth muscle sets specific to their differentiation^54,55^. Examining rhythmic transcripts indicated that Wnt/β-catenin and retinoic acid signaling activities are likely higher in nephron progenitors in pre-branching (differentiation-biased) niches, while YAP activity is likely higher in those in mid-/post-branching (renewal-biased) niches (**Note S2**). We validated this for Wnt/β-catenin and YAP by whole-mount immunofluorescence in E17 mouse kidneys (**Fig. S14**, **Note S1**). Supporting a causal role for the relative timing of these pathways, we found that appropriately timed YAP and RA activation is sufficient to bias the response of human iPSC-derived nephron progenitor organoids to Wnt/β-catenin signaling in favor of renewal and differentiation, respectively (**Note S3, Fig. S15-S17**). ToppGene transcription factor binding site over-representation analysis on rhythmic nephron progenitor genes highlighted E2F^68^, MAZ^69^, LEF1^70^ and MYC^71^ in the proliferation/differentiation phase, all with known involvement in this process (**Fig. 3E**). Notable stromal transcription factors included TEAD (Hippo/Yap) and SRF, which have cooperative and independent roles in cell differentiation along the cortical to medullary stromal conveyor belt^54^, and similar roles in the lung mesenchyme^72^. Hierarchical clustering and differential expression analyses detected both nephron progenitor and stromal rhythms in 4 of 5 experiments after E15 in mouse, and in the human week 20 but not week 17 experiments (**Fig. 3F**, **Figs. S5-S13)**. These data reveal temporal synchronization between ureteric bud branching and differentiation in adjacent streams of nephron and stromal progenitors (**Fig. 3G**, **Movie S3B**).

### Nephron progenitor and stromal rhythms derive from shifts in sub-population composition in the niche

We next sought to place niche transcriptional rhythms in a single-cell context to test for underlying sub-population flux. Sub-clusters were identified after integrating Xenium data with scRNAseq data from E17 mouse nephrogenic zone cells and published week 17 human kidney data^40^ (**Fig. 4A,B**, **Fig. S18A,B**). The data indicate that nephron progenitor and stromal rhythms occur through changes in cell sub-cluster occupancy (**Fig. 4C,D, Fig. S18C,D, Fig. S19A,B, Fig. S20A,B**). Primed/committing nephron progenitors are at their highest proportion in the pre-branching phase, supporting ‘just in time’ delivery to newly forming PTAs in that phase via rhythmic differentiation. This argues against constant-rate differentiation and cell accumulation over the branching lifecycle (**Fig. 1G**). Milder composition shifts were observed among ureteric tip/trunk clusters (**Fig. S19C,D; Fig. S20C,D**). Dimensionality reduction with CONCORD^73^ showed that differentiating nephron progenitors are close in UMAP space to cells ‘orbiting’ in the cell cycle in mouse (**Fig. 4E, Fig. S18E**). This suggests a coordination with the UB branching life-cycle, both of which are ∼24 hr after E15 (ref. ^13,74^). In human, the data corroborate existence of UB-adjacent and -distant streams of nephron progenitors that join the forming nephron at staggered positions and times^40,52^, while revealing a proliferation signature in the UB-adjacent stream (**Note S4**, **Fig. S21**). These data corroborate cell state transitions dependent on ureteric bud branching life-cycle, with coordination between cell cycle and nephron progenitor differentiation.

**Fig. 4:**
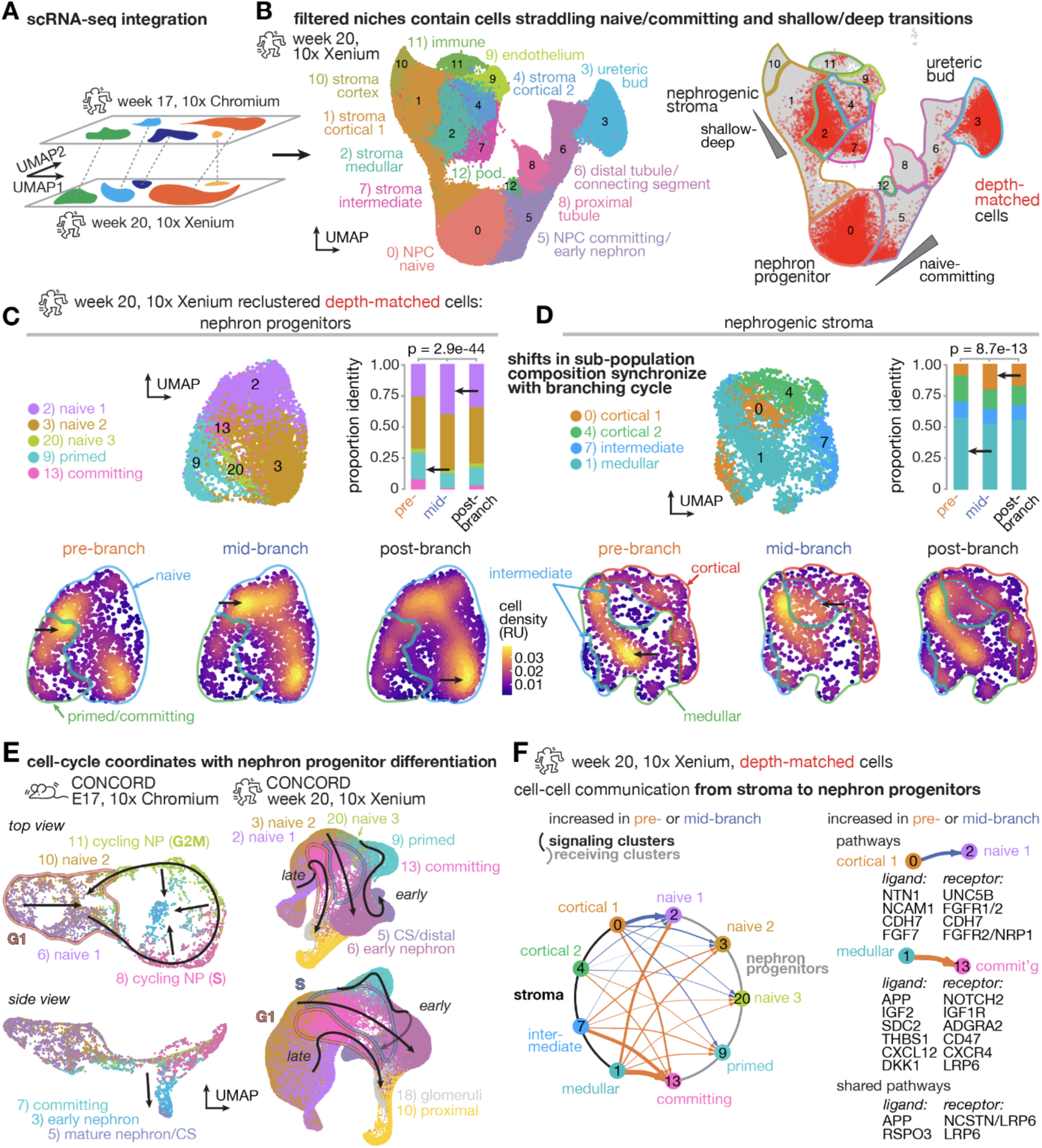
Nephron progenitor and stromal rhythms arise from ‘just in time’ changes in niche sub-population composition. (**A**) Schematic of single-cell integration of the human week 20 Xenium data with published human week 17 nephrogenic zone cell scRNA-seq through canonical correlation analysis. (**B**) UMAP representation of the latent integrated CCA space with Xenium data labeled by cluster (*left*) and by membership in niches filtered for comparable anatomical location using depth metric (*right*). (**C,D**) *Top left*, UMAPs of isolated depth-filtered nephron and stromal progenitors re-clustered from assignments in (A). *Top right*, Bar charts showing proportion of progenitor identity by niche branching stage. Significant deviation from expected frequencies of cell type identity were detected by Chi-squared testing. *Bottom*, UMAPs colored by relative local cell density. Black arrows indicate notable differences in the data. Note that ‘cortical’, ‘medullar’, etc. labels relate to stroma in the nephrogenic zone, not kidney-wide. (**E**) UMAP representation of CONCORD latent space from nephron lineage identities in mouse E17 nephrogenic zone cell Chromium scRNA-seq and human week 20 Xenium spatial scRNA-seq datasets, respectively, colored by cluster assignment. Arrows are qualitative trajectories. (**F**) Cell-cell communication inference through CellChat. *Left*, Circle plot showing differential cell-cell communication interaction weight between clusters originating from pre- or mid-branching niches. Arrow thickness indicates size of differential interaction weight and color indicates branching stage with highest weight. *Right*, Notable ligand/receptor pairs for selected cluster interactions.

Cell-cell communication analysis by CellChat^75^ predicted rhythmic stroma-to-nephron progenitor signaling between cortical (shallow)-to-naive and medullar (deep)-to-committing cell states in pathways consistent with gene set over-representation analyses (**Fig. 4F, Fig. S18F**). However, UB-to-nephron progenitor communication analysis did not indicate a rhythmic component (**Fig. S19E, Fig. S20E**). These data focused our attention on stromal feedback as a possible mechanism for nephrogenesis pacemaking.

### The stromal rhythm reflects a biophysical transition relevant to nephrogenesis

The stromal ribbons separating nephron progenitor niches create a molecular microenvironment that participates in regulating nephron progenitor renewal vs. differentiation^16,28,76–81^. However, the potential mechanical role of the stroma is less explored. Relative to ureteric bud and nephron progenitor cells, we found that dissociated nephrogenic zone stromal cells had higher traction force and phospho-myosin light chain (pMLC) levels^82^, a marker of cell tension^83–85^. *In vivo*, we measured lower nuclear area and higher nuclear aspect ratio in ribbons relative to stromal cells lying above the niche (ref. ^86^ and **Fig. 5A,B**; **Fig. S22** for mouse), another indicator of tension^86,87^. Along with maintenance of normal niche size, this nuclear elongation requires actomyosin activity^86^, indicating that ribbons form a tensile network. Stromal nuclear aspect ratio also correlated with UB branching stage — highest in ribbons lining pre-branching niches (**Fig. 5B**). This matches a higher ribbon tension in pre-branching niches that we inferred by tissue rebound upon laser ablation^86^. The data together indicate that stromal ribbon cells are under rhythmic tension synchronized with UB branching (**Movie S3C**). This may be caused by branching-driven contributions to local niche pressure^86^. These features evoke a rolling mill analogy, where viscoelastic dough (stroma) drops between rollers (nephrogenic niches), being compressed and elongated into sheets (stromal ribbons).

**Fig. 5:**
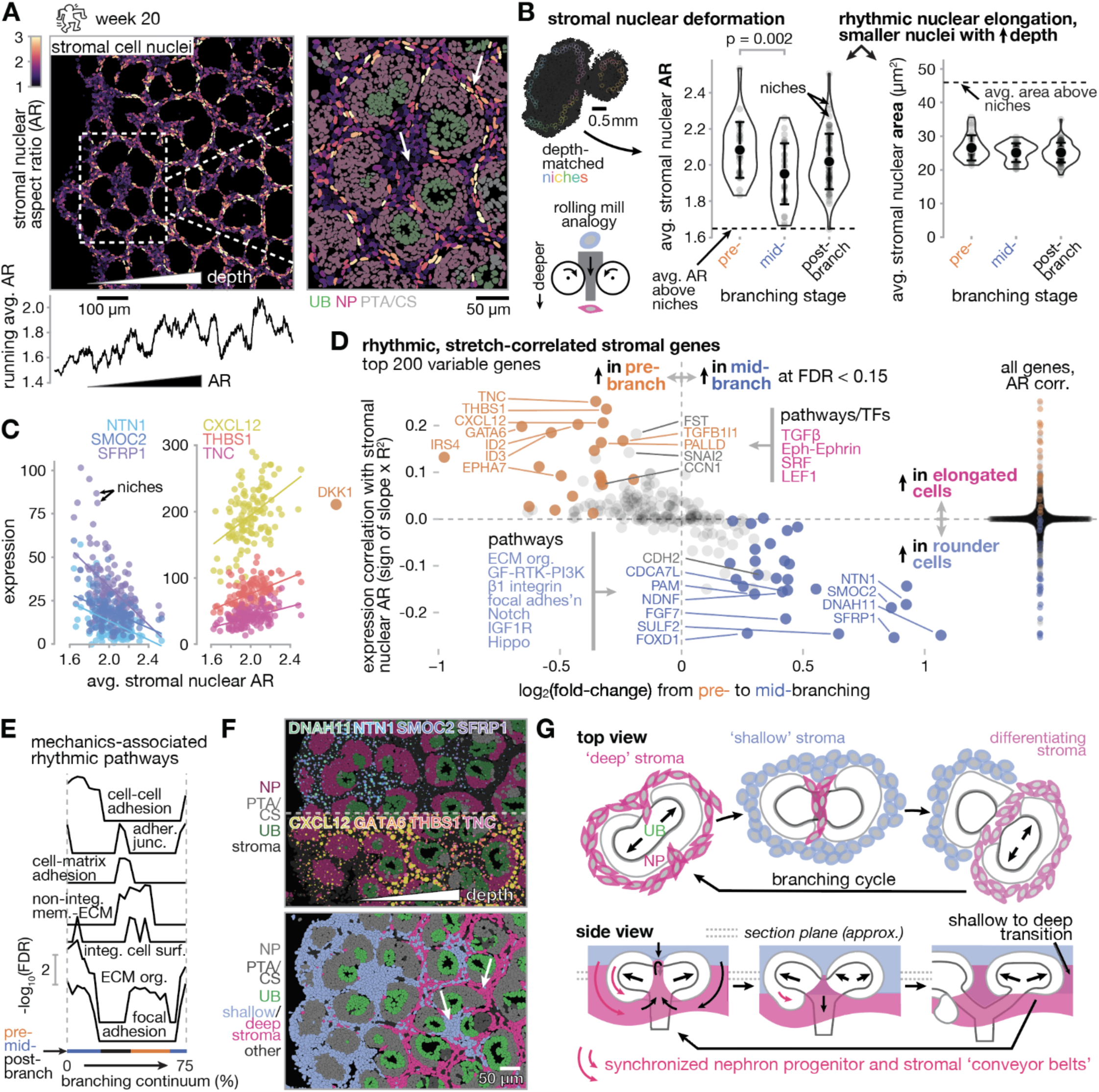
The stromal rhythm correlates with a biophysical transition during differentiation and descent of coordinated ‘conveyor belts’ of stroma and nephron progenitors. (**A**) *Top*, heatmap of stromal cell nuclear aspect ratio (AR) from week 20 human kidney Xenium slice, inset shows detail relative to other niche compartments colored by k-means cluster. White arrows highlight spatial heterogeneity in stromal nuclear AR among niches and within families of niches. *Bottom*, plot of running average stromal nuclear AR vs. horizontal distance from tissue periphery inward (i.e. with increasing tissue depth). (**B**) *Left*, Xenium view of stromal cells lining niches in a similar anatomical depth zone. *Right*, Plots of average stromal nuclear aspect ratio and area vs. niche branching stage (mean ± SD; *n* = 120 niches). (**C**) Plots of example nuclear AR-correlated transcripts. (**D**) Plot of transcript correlation with nuclear AR vs. degree of rhythmic variation over the branching lifecycle. Transcripts in the top left quadrant are enriched in pre-branching niches and stromal cells with higher aspect ratio. Notable ToppGene pathways and transcription factor binding sites associated with AR-correlated transcripts are listed. (**E**) Line plots of GO/KEGG/reactome analyses performed using a sliding window over rhythmic genes fitting a sine wave model at p < 0.2 and ordered by peak position along the branching continuum axis (cell-cell adhesion, cell-cell adhesion via plasma-membrane adhesion molecules; adher. junc., adherens junction; non.-integ. mem.-ECM, non-integrin membrane-ECM interactions; integ. cell surf., integrin cell surface interactions; ECM org., extracellular matrix organization). (**F**) *Top*, Xenium view of notable hits from (**D**) overlaid on niche compartments colored by k-means cluster; point number and size indicate relative transcript density. *Bottom*, Stromal cells in the same field, colored by shallow/deep state scored by gene set expression. (**G**) Schematic model of rhythmic changes in stromal cell biophysical and differentiation state local to the niche over branching lifecycle.

We used nuclear stretch as a proxy for ribbon tension to study its intersection with the stromal transcriptional rhythm. Indeed, rhythmic genes were strongly enriched in nuclear aspect ratio-correlated genes, particularly those characteristic of shallow and deep stromal states (**Fig. 5C,D**). This revealed that stroma with elongated nuclei in the deeper differentiated state surrounds niches early in branching. Conversely, stroma with rounder nuclei in the shallow, less differentiated state surrounds niches later in branching. ToppGene pathway and transcription factor binding site over-representation analysis on nuclear stretch-correlated genes returned similar results to those for rhythmic genes (notably WNT, Hippo/YAP, SRF), but with an additional emphasis on cell biophysical features such as focal adhesion, β1 integrin, and ECM organization gene sets (**Fig. 5D**; cf. **Fig. 3D,E**). Revisiting the sliding window analysis of rhythmic genes also revealed mechanics-associated pathways (**Fig. 5E**). Similar observations hold for the E17 mouse kidney (**Fig. S22**). These correlations account for heterogeneity in the niche stromal microenvironment at the individual branch level in mouse (**Fig. S22F**) and additionally at the branch family ‘rosette’^88^ level in human (**Fig. 5F**). Specifically, shallow-state stroma infiltrates into nephrogenic zone ribbons during mid-branching (**Fig. 5G, Movie S3B, Movie S4**). Once the next branching cycle begins, these cells elongate as part of a biophysical transition during their differentiation to deep-state stroma. This may be a response to local ribbon mechanics. Together the data support a strong intersection between stromal ribbon tension, nuclear stretch, and differentiation.

Several rhythmic genes in stroma that correlated with nuclear aspect ratio have known non-autonomous effects on nephrogenesis as secreted/ECM proteins, including FGF7 (ref. ^89,90^), NTN1 (refs. ^91,92^), SFRP1 (ref. ^93^), and CXCL12 (ref. ^76^). For example, recombinant SFRP1 reduced nephrogenesis and ureteric bud branching in rat embryonic kidney explants, indicating a potential role in preserving nephron progenitor naivety or in preventing ectopic nephron formation^93^. We clarified that SFRP1 preserves nephron progenitor naivety in mouse embryonic kidney explants (**Note S5**, **Fig. S23**). SFRP1 serves as one example of rhythmic, nuclear shape-correlated factors in nephrogenic zone stroma that take part in nephron progenitor regulation. This prompted us to study the connection between stromal mechanics and nephrogenesis in further detail.

### The stroma is a mechanically entrained pacemaker for nephrogenesis

One hypothesis for coordination between the nephron progenitor and stromal conveyor belts^58^ is that stromal differentiation responds to branching mechanics, in turn modifying the nephron progenitor microenvironment to alternatingly promote and inhibit nephrogenesis. YAP is an important mechanotransducer^94,95^ and contributes to stromal cell differentiation with SRF^54,72,96,97^. Drake *et al.* found that YAP is required for differentiation of deep pericyte, fibroblast, and myofibroblast subpopulations from shallow FOXD1+ stroma^54^. Supraphysiological YAP activity by stromal LATS1/2 knockout caused precocious and ectopic differentiation of myofibroblasts and loss of nephrogenesis. A similar role for YAP in shallow-to-deep differentiation was found in the lung mesenchyme^72^. Drake *et al.* found relatively low YAP nuclear localization in the nephrogenic zone stroma relative to deeper stroma. However, stromal Yap/Taz knockout reduced shallow (*Ntn1*, *Ogn*, *Sfrp1*, *Foxd1*, *Smoc2*), deep (*Igfbp3, Pdgfrb, Zeb2*) and myofibroblast (*Myl9*, *Tagln*/SM22a, *Acta2*/aSMA) markers indiscriminantly. This indicates that stromal YAP activity has effects on cell identity that span the nephrogenic zone.

Prompted by rhythmic changes in stromal cell tension^86^, shape, mechanics-associated pathways and a WNT-Hippo/YAP-SRF signature, we quantified tissue depth and branching life-cycle effects on stromal pMLC levels and YAP nuclear:cytoplasmic ratio in mouse kidneys (**Fig. 6A,B**). Both indicate the mechanical state of cells^83–85,98,99^. We found that pMLC and nuclear YAP levels peaked in stromal ribbons at around the midplane of the nephrogenic zone. Both also correlated with the UB branching life-cycle, peaking at the post- to pre-branching transition at which ribbon tension builds^86^ and stromal cells elongate and differentiate. These data support a model in which ribbon tension acts through YAP-SRF to induce stromal differentiation during progenitor descent through ribbons (**Fig. 6B**).

**Fig. 6:**
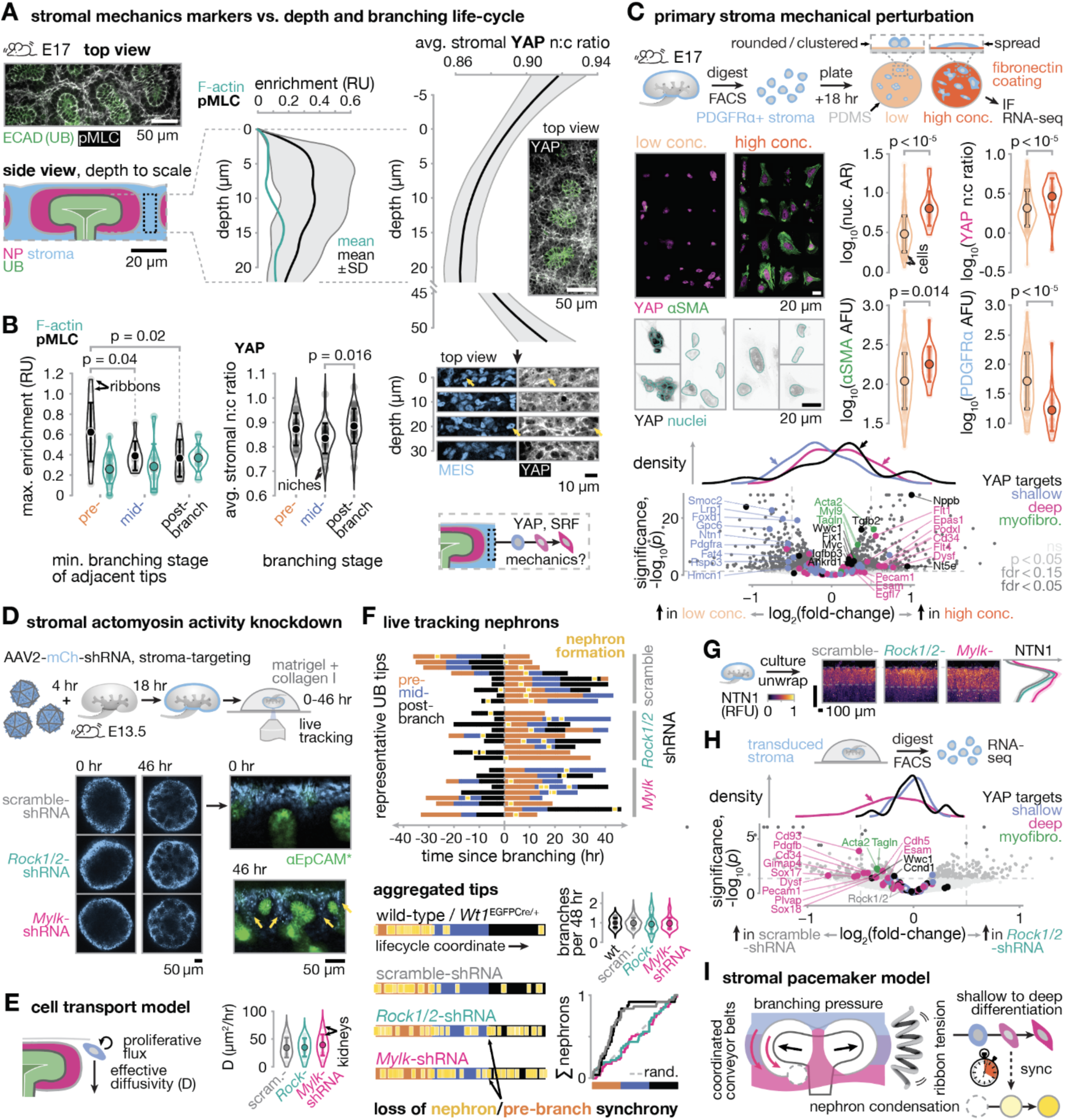
Stromal tension is sufficient for its differentiation and necessary for nephrogenesis pacemaking. (**A**) *Left*, Whole-mount immunofluorescence micrograph of phospho-myosin light chain (pMLC) enrichment in E17 kidney stromal ribbons at the niche midplane. *Right*, pMLC ‘enrichment’ relative to plane-wise average (*n* = 36 ribbons). F-actin serves as a non-rhythmic, uniformly-distributed control for comparison in (A) and (B), *n* = 36 ribbons. Average YAP nuclear:cytoplasmic (n:c) ratio (*n* > 30 niches) in stromal ribbons vs. tissue depth. Representative micrographs are shown for the indicated depths; yellow arrows, representative nuclei. (**B**) *Left*, Plots of UB branching life-cycle dependence of maximum pMLC enrichment and average niche midplane YAP n:c ratio (*n* = 85 niches across 3 kidneys) in stromal ribbons. *Right*, Schematic hypothesis for mechanically-triggered differentiation of stroma from shallow to deep states. (**C**) *Left*, Representative confocal immunofluorescence average projections of YAP localization and aSMA differentiation marker staining with channel intensity normalized to DAPI. Cells have been re-arranged into a grid. *Below*, Representative average projections. *Right*, Plots of nuclear aspect ratio, YAP nuclear:cytoplasmic ratio, and aSMA differentiation and PDGFRA naive marker fluorescence per cell. Data are representative of results from two independent litters. *Bottom*, Bulk RNA-seq differential expression analysis among adhesion conditions (*n* = 3 litters). Gene sets for stromal deep vs. shallow state (from PC1 of Xenium data), myofibroblast markers, and YAP target genes are overlaid, with density distributions along the fold-change axis. (**D**) *Top*, Schematic of AAV2 transduction of mouse embryonic kidney cortical nephrogenic stroma prior to 3D explant culture. *Bottom*, Live confocal fluorescence sections and ‘unwrapped’ insets of stromal cell infiltration (yellow arrows). (**E**) *Left*, Schematic of cell transport model capturing cortical stromal cell proliferative flux and infiltration rate (effective diffusivity) as fitting parameters. *Right*, Plot of effective diffusivity vs. shRNA treatment. (**F**) *Top*, Ticker-tape plots of nephron formation events relative to ureteric bud tip branching stage after aligning each tip to a common time since branching initiation; representative data for *n* = 29, 33, 44 tips across 9, 10, 9 kidneys each for scramble-shRNA, *Rock1/2*-shRNA, and *Mylk*-shRNA conditions respectively. *Bottom left*, Aggregated ticker-tapes in which nephron formation events are plotted at the fraction of the corresponding branching stage that they formed in. Stages are idealized as equal thirds of the branching life-cycle. *Bottom right*, plots of branching events per 48 hr estimated as # stage transitions / 3, and cumulative distributions of nephrons formed relative to the idealized ticker-tape timelines. (**G**) Average projection immunofluorescence micrographs and intensity profiles for cultured kidneys sectioned and stained for shallow stromal marker NTN1. (**H**) Bulk RNA-seq differential expression analysis among shRNA treatment conditions (*n* = 3 litters). (**I**) Schematic model for stromal mechanosensing and nephrogenesis pacemaking. Plots with error bars or envelopes are all mean ± SD.

To test this, we reasoned that mimicking branching-derived tension in primary stromal cells should phenocopy stromal differentiation. We cultured PDGFRA+ primary nephrogenic zone stromal cells on silicone substrates with low- vs. high-adhesion fibronectin coatings (**Fig. 6C**). In this setting, tension due to adhesion is transmitted to the nucleus via the cytoskeleton and LINC complex, triggering YAP nuclear translocation^99^. Relative to those on low-adhesion substrates, cells on high-adhesion substrates increased in nuclear aspect ratio, YAP nuclear:cytoplasmic ratio, and myofibroblast marker alpha–smooth muscle actin (ɑSMA) staining; while decreasing in staining for the naive marker PDGFRA. Bulk RNA-seq showed that these cells increased their expression of YAP target genes^100,101^ (*Ankrd1, Nppb, Wwc1*), myofibroblast/SRF markers^54^ (ɑSMA/*Acta2*, transgelin*/Tagln*, *Myl9*), and deep genes relative to shallow genes (taken from the first principal component of Xenium data). Several of the latter genes are associated with the vasculature^55^, including *Flt1* (VEGFR1) and *Pecam1*. ToppGene analysis of upregulated genes confirmed over-representation of smooth muscle and SRF signatures. These data indicate that cell tension induced by adhesion *in vitro* is sufficient to phenocopy changes in YAP activation and aspects of the differentiation trajectory of stromal cells as they transition from a shallow to deep state *in vivo*.

We reasoned that disrupting tension-driven stromal differentiation would reveal if it is necessary for nephrogenesis pacemaking. We found that adeno-associated virus type 2 (AAV2) selectively transduces shallow stromal cells superficial to ureteric bud midplanes (i.e. cortical nephrogenic zone stroma, **Fig. 6D**, **Fig. S24A-C**)^102^. Since non-muscle myosin II contractility is required for YAP mechanosensing of cell tension^94,99,103^, we used AAV2 to deliver shRNAs against two myosin II activators — ROCK1/2 and myosin light-chain kinase (*Mylk*/MLCK). We confirmed >2-fold knockdown of each by western blot in NIH/3T3 mouse embryonic fibroblasts by 48-72 hr (**Fig. S24D**). E13.5-14.5 mouse embryonic kidneys in 3D culture showed comparable cortical to medullary flow of transduced stroma regardless of the shRNA delivered (**Fig. 6D**, **Movie S5**). Fitting fluorescence data to a transport model showed that the timescale of stromal infiltration ∼ *x*^2^/2*D* = 34 hr is comparable to the branching timescale *in vivo* and in 3D culture^13,48,74^ (**Fig. 6E**). UB branching rate was also similar among the conditions (**Fig. 6F**). However, nephron condensation events showed a striking loss in synchrony with the UB branching life-cycle upon stromal *Rock1/2*-shRNA or *Mylk*-shRNA treatment but not control scramble-shRNA treatment (**Fig. 6F**). The same result held for *Rock1/2*-shRNA vs. scramble-shRNA in *Wt1*^EGFPCre/+^ kidneys, which otherwise had comparable PTA formation dynamics with normal proximal cell recruitment (**Fig. S24E**). Though they formed normally, nascent connecting segments in *Rock1/2/Mylk*-shRNA-treated kidneys were often closer to the surface relative to those in control kidneys (**Movie S5**), indicating a potential change in the local position of the inductive front. Staining for the shallow marker Netrin (NTN1) revealed that though stromal infiltration was intact, NTN1 expression persisted deeper in the nephrogenic zone in *Rock1/2/Mylk*-shRNA conditions, indicating delayed differentiation from a shallow to deep cell state (**Fig. 6G**). Bulk RNA-seq on FACS-sorted transduced stroma confirmed this, showing depletion of myofibroblast and other deep stromal markers upon *Rock1/2*-shRNA treatment relative to scramble-shRNA control virus (**Fig. 6H**). Again, the latter included genes associated with the vasculature^55^, including *Cdh5* (VE-cadherin), *Plvap*, and *Sox17*. ToppGene analysis of downregulated genes confirmed over-representation of vasculature/pericyte development, smooth muscle, and SRF signatures. This suggests a mechanical origin for the observed loss of differentiation of pericytes (autonomously) and peritubular capillaries (non-autonomously) upon stromal YAP/TAZ knockout^54^. Together the data indicate that stromal cell actomyosin activity is necessary for stromal differentiation and pacemaking of adjacent nephron progenitor commitment to early nephrons.

## Discussion

Nephron progenitors are stratified from naive to primed with depth into the nephrogenic zone^34,45^. Paradoxically, these cells migrate freely within their cap compartment, forming a dynamic equilibrium among states/layers^42^. These features are partially shared by other stem cell niches that form ‘proliferative conveyors’ governed by neutral drift dynamics^104–110^, such as hair follicles and intestinal crypts. However, nephrogenic niches must form discrete nephrons rather than a continuous stream of differentiating cells. Our data reveal a rhythmic component to niche behavior, where the sub-population composition of the cap alternates between a renewal and differentiation bias in synchrony with the branching program of the adjacent ureteric bud tip. This is accompanied by a parallel rhythm in surrounding stroma. Though retrograde transport of cells in both streams relative to the advancing nephrogenic zone was predicted by lineage-tracing^58^, the two streams have a tighter rhythmic coordination than previously recognized. We found that ‘just in time’ nephron progenitor differentiation and proliferation/cell cycle rhythms are coupled, prompting future investigation into whether this is required for proper early nephron composition^111^ or size^112^, as recently reported for somite formation^112^.

Turning to the underlying clock, none of the signals that coordinate niche cell compartments upstream of nephron condensation are explicitly rhythmic^15–18,25,28,90,113–117^, and no direct mechanism for feedback from branching morphogenesis of the ureteric bud has been established. This process is an obvious candidate for the pacemaker, since it is a largely independent, autonomously rhythmic input so long as growth factors including GDNF are present^118,119^. However, there was a puzzling lack of evidence for direct rhythmic signaling between this compartment and nephron progenitors in our spatial sequencing data, relative to the stroma. FOXD1+ stroma and its daughter lineages contribute to nephron progenitor renewal:differentiation balance^16,28,76–81^. We find that the ratio of naive to differentiating stromal cells local to the niche changes rhythmically over the UB branching cycle. Our data propose that stromal ribbon tension induced by branching-derived niche pressure drives these compositional changes, particularly through myofibroblast differentiation and vascular recruitment. This is necessary for nephrogenesis pacemaking, entraining the nephrogenesis rhythm to the ureteric bud branching rhythm. Indeed, stromal differentiation along the corticomedullary axis is known to coordinate smooth muscle patterning, vascular recruitment, and ureteric bud tree elaboration^54,91,92,120^; non-autonomous effects on nephron progenitors are an additional category^76,91^. The latter are most likely mediated by signals in our list of paracrine/ECM factors that correlate with stromal nuclear stretch. Alternatively, rhythmic stromal tension could control nephron pacing through a parallel, independent mechanism in nephron progenitors. For example, there is an intriguing overlap between somite and nephron progenitors in their rhythmic condensation dynamics, pathways (Wnt and Notch/FGF-MAPK^121–123^), and markers^121^ (*Hes1*, *Bcl2l11*, *Has2*, *Tnfrsf19*, *Id1*, *Ptpn11*, *Dusp6*, and *Myc*; see also somite development gene set in **Fig. 3D, Fig. S3E**). The relatively longer period between nephron condensation events could be explained by an interplay between niche pressure and stromal compliance^86^, which could convert oscillations into excitable pulses gated by YAP activity^124^. In this model, stromal differentiation would separately coordinate vascularization with the forming nephron. Whether the mechanical pacemaking role of the stroma depends on its differentiation remains to be determined.

Our findings reveal that mechanical feedback enables the kidney to appropriately pace nephron formation events relative to the ureteric bud branching program. This feedback is distinctive among similar stem cell niches, discretizing cell commitment along an otherwise continuous conveyor. This may lend robustness to tissue composition during exponential organ growth.

## Acknowledgements

We thank Kieran Short, Zev Gartner, Lukasz Bugaj, and Louis Prahl for helpful discussion. We thank Kyle McCracken and Pedro Medina for advice on kidney organoid air-liquid interface culture and immunofluorescence, and Carol Gao for assistance with organoid culture. This work was performed in part at the Penn Medicine Center for Molecular Studies In Digestive and Liver Diseases, Molecular Pathology and Imaging Core (MPIC, funded under NIH center grant P30-DK050306). This research was partially supported by the NSF through the University of Pennsylvania Materials Research Science and Engineering Center (MRSEC, DMR-2309043). Cell sorting was performed on a BD FACSAria Fusion maintained by the Penn Cytomics and Cell Sorting Resource Laboratory and obtained in part through NIH grant 1S10OD026986. This work was supported by the NSF GRFP (J. Liu), Predoctoral Training Program in Developmental Biology T32HD083185 (SHG, JMV), Trainee Pilot Award from Center for Engineering MechanoBiology (CEMB), an NSF Science and Technology Center, under CMMI: 15-48571 (AZH), NIH NIDDK R01DK110792 and R01DK054364 (APM, NOL), NIH NIGMS MIRA R35GM133380 (AJH), NIH NIDDK R01DK132296 (AJH) and R01DK140070 (AJH), NSF CAREER award 2047271 (AJH), and Penn Center for Precision Engineering for Health (CPE4H) pilot grant (AJH). We acknowledge support relating to iPSC lines from NIH NIDDK P50DK133943, and Open Philanthropy funds from the Silicon Valley Community Foundation and the Good Ventures Foundation to K.S. No federal funds were used for the study of human kidney development.

## Author contributions

S.H.G., S.N.D., J.M.V., A.H., and A.J.H. designed experiments. A.E.S., L.S., J. Levinsohn, A.P.M, N.O.L., L.W., K. Sasaki, and K. Susztak provided kidney tissues and/or image data. S.H.G., S.N.D., J.M.V., A.H., G.Y.L., J. Liu, K.B., G.Q., C.M.P., A.E.S., and L.S. performed experiments. S.H.G., S.N.D., I.G., J.M.V., A.H., R.S.K., M.J.Y., K.Y., and A.J.H. analyzed data. S.H.G., I.G., and A.J.H. created code and/or computational models. S.H.G., S.N.D., and A.J.H interpreted data and wrote the manuscript. All authors reviewed the manuscript.

## Materials & correspondence

Requests for further information, resources, and reagents should be directed to Alex Hughes.

## Data and code availability

Microscopy data reported in this paper will be shared by the lead contact upon request. Raw and processed spatial sequencing, scRNAseq, and bulk RNA sequencing data files have been deposited in the Gene Expression Omnibus (GEO) under the series accession number GSEXXXX. Other processed data files including further spatial sequencing files, and original code files have been deposited in Mendeley Data (DOI: 10.17632/v9z6bbdp6d.1) and are publicly available. Any additional information required to reanalyze the data reported in this paper is available from the lead contact upon request.

## Methods

### Embryonic kidneys

Mouse protocols followed NIH guidelines and were approved by the Institutional Animal Care and Use Committees of the University of Pennsylvania and the University of Southern California. Mice were housed in standard ventilated cages (single occupancy) in a conventional rodent facility with 12 h light cycle and free access to water and food. For wild-type kidney experiments, embryos were collected from timed pregnant CD-1 mice that were 8-10 weeks old upon receipt (Charles River Laboratories, RRID:IMSR_CRL:022). Embryonic kidneys were dissected in chilled Dulbecco’s phosphate buffered saline (DPBS, MT21-31-CV, Corning)^125^. For *Wt1*-reporter kidney experiments, timed matings between *Wt1*^GFPCre/+^ (The Jackson Laboratory #010911) and wild-type animals were conducted to obtain embryos at E13.5. *Wt1*^GFPCre/+^ embryos express an EGFPCre fusion protein driven by *Wt1* promoter/enhancer elements^51^. While the endogenous *Wt1* allele is inactivated, the loss of one allele has minimal impacts on embryonic kidney development^49,50^. Embryo genotype was confirmed by EGFP expression using confocal microscopy. Embryo ages were confirmed by limb staging^126^. The sex of embryos was not determined and not taken into consideration in the study design.

For *Foxd1*-reporter kidney experiments, *Foxd1*^GFPCre/+^; *Rosa26*^mTmG/+^ embryos were harvested at E11.5-E12.5 from timed matings. Kidneys were dissected and cultured overnight at 37°C on a Transwell filter (Corning) in FluoroBrite DMEM (Life technologies, A18967–01) supplemented with 10% fetal calf serum, 1% pen/strep, and 1X Glutamax (Thermofisher). Filter inserts were transferred to 35mm MatTek glass bottom dishes in customized metal holders and imaged over 24-72 hr using a Leica SP8 system using an HC FLUOTAR L 25x/0.95 water immersion objective. The water immersion was maintained through a Leica water cap with a continuous water supply and drainage system allowing for flow of water. CO_2_ and humidity levels were maintained using a stage-top Ibidi control system.

Human fetal kidneys were obtained from donors who had provided informed consent and underwent elective abortion at the University of Pennsylvania. All experimental procedures were approved by the Institutional Review Board at the University of Pennsylvania (#832470). Embryo ages were determined through ultrasonographic measurement of crown-rump length. Human kidney tissue was dissected in chilled RPMI-1640 (Gibco) and processed at 4°C by fixing in 10% neutral-buffered formalin overnight. Tissues were dehydrated using 50% followed by 70% ethanol (EtOH) for 2 hr each before storage in 70% EtOH, all at 4°C. Tissues were then cut into ∼5 mm blocks using a razor blade.

### Confocal fluorescence imaging

Imaging was performed using a Nikon Ti2-E microscope equipped with a CSU-W1 spinning disk (Yokogawa), a white light LED, laser illumination (100 mW 405, 488, and 561 nm lasers and a 75 mW 640 nm laser), a Prime 95B back-illuminated sCMOS camera (Photometrics), motorized stage, 4x/0.2 NA, 10x/0.25 NA, 20x/0.5 NA, and 40x/0.9 NA lenses (Nikon), and a stagetop environmental enclosure with temperature and CO_2_ control (OkoLabs).

### Kidney whole-mount immunofluorescence

Mouse embryonic kidney staining was performed as previously described^127^, using protocols adapted from Combes *et al.* and O’Brien *et al.^117,128^*. Briefly, dissected mouse kidneys were fixed in ice cold 4% paraformaldehyde in DPBS for 20 min, washed three times for 5 min per wash in ice cold DPBS, blocked for 2 hr at room temperature in PBSTX (DPBS + 0.1% Triton X-100) containing 5% donkey serum (D9663, Sigma), incubated in primary and then secondary antibodies in blocking buffer for at least 48 hr at 4°C, alternating with 3 washes in PBSTX totaling 12-24 hr. The minimum duration of primary and secondary antibody incubations and washes depended on the age of the kidney, as previously described^128^. In some experiments, stained mouse kidneys were cleared for 2 days in ScaleA2 (4 M urea + 0.1% Triton X-100 + 10% glycerol) followed by 2 days in ScaleB4 (8 M urea + 0.1% Triton X-100)^129^, and imaged in ScaleA2.

Human embryonic kidney cortical sections were manually cut using a razor blade. Sections were then rehydrated in a series of decreasing ethanol concentrations for 1 hr each (70%, 35%, 15%) and stored in 1x DPBS. Samples were then blocked in a solution of 5% donkey serum (D9663, Sigma) in 1x DPBS + 0.1% Triton X-100 (PBS-Tx) for 24 hr at 4°C. Samples were then incubated with antibodies described below diluted in blocking buffer for at least 72 hr at 4°C, then washed in 3 exchanges of ice cold PBS-Tx at 4°C for at least 3 hr each, then incubated in secondary antibodies described below for at least 48 hr at 4°C, followed by a repeat of the PBSTX washing steps described above. Prior to imaging, samples were resuspended in FocusClear (CelExplorer Labs Co., FC-101) to aid visualization.

Primary antibodies and dilutions included rabbit anti-SIX1 (1:200, 12891, Cell Signaling Technology, RRID: AB_2753209), goat anti-RET (1:200, AF1485, R&D Systems, RRID: AB_354820), rabbit anti-SIX2 (1:600, 11562-1-AP, Proteintech, RRID: AB_2189084), goat anti-ITGA8 (1:20, AF4076, R&D Systems, RRID: AB_2296280), mouse anti-E-cadherin (1:50, clone 34, 610404, BD Biosciences, RRID: AB_397787), Goat anti-PDGFRA (1:500, AF1062, R&D Systems, RRID:AB_2236897), rabbit anti-phospho-Smad1/5 (1:800, 9516, Cell Signaling Technology, RRID:AB_491015), rabbit anti-Ki67 (0.5 µg ml^-1^, ab15580, Abcam, RRID:AB_443209), mouse anti-MEIS1/2/3 (0.5 µg ml^-1^, 39795, Active Motif, RRID:AB_2750570), rabbit anti-Cyclin D1 (1:200, ab16663, Abcam, RRID:AB_443423), rabbit anti-PAX8 (1:250, 10336-1-AP, Proteintech, RRID: AB_2236705), rabbit anti-YAP (1:200, 11562-1-AP, Proteintech, RRID: AB_2189084), rabbit anti-MYL12A (pS19) (1:100, ab2480, abcam, RRID:AB_303094). Secondary antibodies (all raised in donkey) were used at 1:300 dilution and included anti-rabbit AlexaFluor 647 (A31573, ThermoFisher, RRID: AB_2536183), anti-rabbit AlexaFluor 555 (A31570, ThermoFisher, RRID: AB_2536180), anti-rabbit AlexaFluor 488 (A21206, ThermoFisher, RRID: AB_2535792), anti-mouse AlexaFluor 555 (A31572, ThermoFisher, RRID: AB_162543), and anti-goat AlexaFluor 488 (A11055, ThermoFisher, RRID: AB_2534102). In some experiments, samples were counterstained in 300 nM DAPI (4’,6-diamidino-2-phenylindole; D1306, ThermoFisher) diluted in blocking buffer for 2 hr at room temperature, followed by 3 washes in PBS.

Stained kidney samples were imaged in wells created with a 2 mm diameter biopsy punch in a ∼5 mm-thick layer of 15:1 (base:crosslinker) polydimethylsiloxane (PDMS) elastomer (Sylgard 184, 2065622, Ellsworth Adhesives) set in 35 mm coverslip-bottom dishes (FD35-100, World Precision Instruments). For mouse kidney rhythmic marker validation, average immunofluorescence was quantified for cap mesenchyme regions of interest segmented using Ilastik^130^ v1.4.1b6. Segmented outlines were manually refined in FIJI as required.

### 3D mouse embryonic kidney explant culture

E13.5 mouse embryonic kidney explants were cultured in 3D in 50:50 matrigel:collagen 1 domes as previously described^48^. Briefly, kidneys were incubated at 37°C with fluorophore-conjugated anti-EpCAM or anti-E-cadherin for 2 hr and washed once in PBS before culture. Live labels were FITC-labeled anti-EpCAM/CD326 (11-5791-82, Invitrogen) and AF660-labeled anti-CD324/E-cadherin (50-3249-82, Invitrogen) at 1:250 dilution in culture media. To prepare the hydrogel solution, rat tail collagen I (354236, Corning) was neutralized and diluted on ice with 1 M NaOH, 10X PBS and DI water to achieve a 1 mg ml^-1^, pH 7 solution. The neutralized collagen and human ESC-qualified Matrigel (354277 lot 17823003, Corning) were then mixed at a 1:1 ratio v/v on ice resulting in a 0.5 mg ml^-1^ collagen I/50% Matrigel composite. Labeled kidneys were transferred to 12 mm internal-diameter PDMS rings using a gelatin-coated p20 pipette tip truncated with a razor blade.

Excess liquid was removed and 10 µl of hydrogel solution was added to suspend the kidney within a droplet. The outside of the PDMS rings was filled with PBS for humidity control, and the culture plate was incubated at 37°C for 30 min to set the hydrogel. Phenol red-free DMEM media (A1443001, Thermo Fisher) with 10% fetal bovine serum (FBS, MT35-010-CV, Corning) and 1x pen/strep (100 IU ml^-1^ penicillin, and 100 μg ml^-1^ streptomycin, 100x stock, 15140122, Invitrogen) was then added to the inside of the PDMS ring after gelation and prior to imaging. 10x confocal *z* stack timelapses were collected every hour at 15 µm step size after pre-defining *xy* positions.

### SFPR1 inhibition in 3D mouse embryonic kidney culture

E13.5 wild-type CD1 kidneys were cultured as described above without use of the live anti-epithelial markers or time-lapse microscopy. Immediately after gelation, kidneys were administered DMEM with 10% FBS, 1% pen/strep including either way316606 (20 µM, Tocris, 4767) for SFRP1 inhibition or equivolume DMSO for the control.

Kidneys were cultured for 72 hr prior to fixation and immunonofluorescence. Kidneys were stained with Donkey antibodies towards rabbit anti-CITED1 (1:250, Invitrogen PA5-65541, RRID:AB_2664550), mouse anti-SIX2 (1:300, Proteintech 66347-1-Ig, RRID:AB_2881727), and rat anti-ECAD (1:300, Abcam ab11512, RRID:AB_298118). Secondary antibodies (raised against donkey and used at 1:300 dilution) included anti-rabbit AlexaFluor 488 (A21206, ThermoFisher, RRID: AB_2535792), anti-mouse AlexaFluor 555 (A31572, ThermoFisher, RRID: AB_162543), and anti-goat AlexaFluor 647 (A32849, ThermoFisher, RRID:AB_2762840). Kidneys were counterstained using 2.5 µg ml^-1^ DAPI in DPBS and washed once in DPBS before imaging using confocal microscopy with a 20x objective (see **Kidney whole-mount immunofluorescence**). Cap mesenchyme and ureteric bud tip ROIs from niches in-plane with the x-y plane were manually segmented using FIJIs polygon selection tool. ROIs were segmented using the z-plane where UB tip lumen was first visible to keep relative depth consistent between niches. Cap mesenchyme z-depth was measured from segmented niches by manually identifying the shallowest (z_i_) and deepest (z_f_) in-focus planes of SIX2 signal. Depth was calculated using the following formula: (z_f_ - z_i_) x step size, with step size equal to 2 µm. A custom CellProfiler^131^pipeline was used to measure DAPI, SIX2, CITED1, and ECAD channel fluorescence from the manually segmented ROIs. CellProfiler output was analyzed in R for statistical testing and plotting.

### Live nephrogenesis tracking

Mouse kidney explant culture timelapses were registered in xyzt using manually annotated landmarks in FIJI. Ticker-tape data consisted of nephron stage annotation at each timelapse frame for UB tips for which at least one nephron initiated and for which there was a least one transition from post- to pre-branching over the course of the timelapse. Discrete rather than continuum branch staging was estimated as described in the main text due to tips being at various orientations.

The frame(s) at which nephron condensation were first perceptible were then associated with the corresponding branching stage ticker-tape. Further processing included producing aggregated ticker-tapes in which nephron formation events were plotted at the fraction of the corresponding branching stage that they formed in. In this case the 3 branching stages were idealized as equal thirds of the branching life-cycle. We also produced plots of branching events per 48 hr estimated as # stage transitions / 3 for the subset of annotated tips, and cumulative distributions of nephrons formed relative to the idealized ticker-tape timelines.

### HCR RNA-FISH

E17 mouse embryonic kidneys dissected in RNase-free PBS were assayed by whole-mount hybridization chain reaction (HCR v3.0) single-molecule RNA fluorescence in-situ hybridization (smFISH) similar to published protocols^132–136^ and consistent Molecular Instruments Inc. protocols. Probe sets were designed using in-house software (AnglerLite) and are provided in **Supplementary Files**). Buffers and hairpin solutions were procured from Molecular Instruments, Inc. Kidneys were transferred to 1.5 ml Eppendorf tubes and fixed in 4% PFA overnight at 4°C. Kidneys were washed 3 x 5 min in PBST and dehydrated on ice by serial 10 min incubations in 25%, 50%, 75%, and 100% methanol:PBST washes before incubation at -20°C for > 16 hr. Kidneys were rehydrated on ice by serial 10 min incubations in 25%, 50%, 75%, and 100% PBST:methanol, followed by one 10 min PBST wash at room temperature. Kidneys were further permeabilized for 1 hr in 2% SDS, 10 µg ml^-1^ proteinase K in PBST. Kidneys were washed 3 x 5 min in PBST, post-fixed 20 min in 4% PFA at room temperature, washed 5 x 5 min in PBST, and incubated in probe hybridization buffer for 5 min. Kidneys were pre-hybridized with pre-warmed probe hybridization buffer for 30 min at 37°C. 100 µl of 16 nM probe solution was then prepared from a 1 µM stock and pre-warmed to 37°C for 15 min before probing kidneys overnight (>12 hr) at 37°C. Probe wash buffer was pre-heated to 37°C and used to wash kidneys for 4 x 30 min at 37°C, followed by 2 x washes with 5x sodium chloride-sodium citrate buffer tween buffer (SSCT) at room temperature. Amplification buffer was equilibrated at room temperature while 2 µl of 3 µM stocks of hairpins 1 and 2 were snap-cooled from 95°C for 90s to 20°C for 30 min in the dark. Kidneys were incubated with pre-warmed amplification buffer for 30 min at room temperature before incubating with 100 µl amplification buffer containing 50x dilutions of each hairpin at room temperature overnight (12-16 hr). Excess hairpins were removed by washing in 5x SSCT at room temperature 2 x 5 min, 2 x 20 min, and 1 x 5 min, followed by 3 x 5 min washes in PBST.

Kidneys were imaged by confocal fluorescence microscopy at 20x and 40x as for antibody assays (see **Kidney whole-mount immunofluorescence.)**

### Spatial transcriptome sequencing

The 10x Xenium Prime 5k enables *in situ*, imaging-based detection of ∼5000 prescribed transcripts in tissue slices. ∼100 extra custom gene probesets were manually curated based on literature search to capture canonical and pathway markers characteristic of cell types and signaling processes operating in the nephrogenic zone that were missing from the default 5000 gene list. Freshly dissected mouse kidneys were partially flattened by applying ∼20% strain normal to the surface plane. This was achieved by sandwiching kidneys between two 1” x 3” standard microscope slides spaced apart using shims cut from 750 µm-thick plastic (Grainger) and held together using rubber bands during fixation for 2 hr in 4% paraformaldehyde (PFA) at 4°C. Human fetal kidney samples stored in 70% EtOH were not flattened. These samples were subjected to further EtOH exchanges at 75%, 90%, 95%, and 100% for 2 hr each at 4°C, washed 3 times in excess PBS, and embedded in paraffin. Tissue preparation, sectioning, deparaffinization/decrosslinking, and Xenium Prime assay followed manufacturer protocols (10x Genomics). Briefly, 5 µm kidney sections were prepared and transferred to Xenium slides using a microtome and flotation bath. Slides were dried and transferred to a desiccator overnight. For deparaffinization, slides were incubated at 60 °C for 2 hr, cooled to room temperature, and then processed through xylene washes and an ethanol series ending with nuclease-free water. After rehydration, FFPE crosslinks were reversed using a decrosslinking buffer followed by two washes in PBST. Xenium Prime barcoding assay steps were then performed, comprising priming hybridization, RNase treatment and polishing, probe hybridization, ligation, amplification, cell segmentation staining (ATP1A1/CD45/ECAD for cell boundaries; 18S rRNA and aSMA/Vimentin for cell interior), autofluorescence quenching, and DAPI staining. Barcode decoding was then performed through automated iterative cycles of fluorescent probe hybridization, imaging, and probe removal on the Xenium Analyzer instrument. For each cycle and image plane, the onboard pipeline detected fluorescent puncta, decoded them into gene identities, and assigned a per-transcript basecalling quality ‘Q score’. Cell boundaries were computed by morphology-based segmentation of DAPI images using a neural-network model, enabling assignment of decoded transcripts to individual cells.

### Pseudobulked spatial sequencing data analysis

All ureteric bud, nephron progenitor, and nephron lineage cells associated with each niche were manually annotated using lasso outlines in Xenium Explorer 3 to create ROIs. Stromal/ureteric bud cells in close proximity to the cap mesenchyme were associated with niches using the FNN package by finding the nearest 2 stromal/ureteric bud cell neighbors to each NP cell (by Euclidean distance between cell centroids) and removing duplicate cell IDs. Cell IDs, K-means cluster number (cell type), cell area, and ROI outline vertex coordinates were exported for further processing in R. Transcript-level data were provided in Xenium output for all cells in parquet format. For each ROI, we performed separate analyses by K-means cluster number. Quality control removed transcripts with Q score <20. We pseudobulked data by computing raw per-gene count tallies across cells falling within each ROI. In parallel, we estimated ROI area as the sum of the per-cell areas (µm^2^). Outputs for each ROI included a gene-by-count table and total area. We next selected ROIs falling within a similar predicted tissue depth range by filtering ROIs based on a ‘depth metric’ created from expression of height-associated genes. A Seurat object^137^ was first created from Q score-filtered per-cell per-gene counts data for all cells in a given slice. These data were log-normalized (NormalizeData) and scaled (ScaleData) to create gene *z*-scores. *Z*-scores were exported as an RDS file with the x,y coordinates of each cell. We defined a scalar depth metric as a weighted sum of height-associated gene *z*-scores. Height-associated genes (*TTC28*, *MEST*, *CITED1*) were previously identified from principal component analysis of pseudobulked ROI expression data, see below. The depth score was computed for each cell as the sum of z-scores of the height associated genes multiplied by the following weights: TTC28 = -1, MEST = -1, CITED1 = -2. We then used mba.surf to create a continuous interpolated surface from the metric values and cell coordinates, which was then Gaussian smoothed and exported as a contour map. We manually selected contours that bracketed ROIs whose depths were anatomically consistent with the approximate midplane of the nephrogenic zone and filtered out ROIs whose centroids (computed from ROI outline vertex coordinates) did not fall between these contours. We next returned to the original pseudobulked count data and filtered out genes having a non-zero count in fewer than 20% of the in-contours ROIs. We then performed between-ROI Q3 normalization. Specifically, for each ROI we computed the 75th percentile of its gene counts, defined the global scaling factor as the median of these ROI-specific Q3 values, and rescaled each ROI by dividing counts by its own Q3 and multiplying by the global median Q3. Median gene expression per ROI scaled with ROI area, which normalization successfully compensated for (**Fig. S2B**). Q3 normalized counts were allocated to the counts argument in CreateSeuratObject and log transformed using a log2(count + 1) function. To reduce noise, we filtered genes by average expression across ROIs, retaining only features with mean Q3-normalized counts of >4-7 per ROI. We then identified the top 1,500 highly variable genes using Seurat FindVariableFeatures with the VST method and created *z*-scores using ScaleData. RunPCA was used to perform principal component analysis on the variable features. Pheatmap using average linkage with correlation distance for both rows and columns was used for unsupervised clustering of niches using the top 100 variable genes overall or of those in the first principal component for stroma. We performed differential expression analysis on the top 200 variable genes between ROIs in different branching life-cycle stages using gene-wise linear models with Benjamini-Hochberg correction for multiple comparisons (FDR). The models controlled for the effect of tissue slice if more than one slice contributed ROIs to a given analysis.

### Branching continuum score

The AlphaShape^138^ algorithm (Ken Bellock) was used to create closed polygons from the ureteric bud and nephron progenitor cell centroids associated with each niche in Python 3.14.0. Polygon coordinates across niches at the same developmental stage and in the same Xenium replicate were registered to a common cartesian origin. 55 geometric features were calculated for each of the two polygons per niche (**Table S2**). The feature matrix was scaled such that each feature had zero mean and unit standard deviation. Principal component analysis (PCA) was performed and the first principal component was linearly scaled to a minimum value of 0 maximum of 100, yielding the branching continuum.

### Validation of branching continuum score

Niches were analyzed and processed similarly to Xenium niches to create branching continuum scores, but for every third frame of a confocal fluorescence timelapse of an E11.5 Foxd1GC/+; R26mTmG/+ mouse kidney explant imaged between 24 and 72 hr of culture time (**Movie S4**). Input polygons here were derived from manual segmentation of the ureteric bud and nephron progenitor compartments of two parent niches and their associated daughter niches in Fiji.

### Rhythm visualization

We curated groups of differentially expressed genes at FDR < 0.15 associated with naive/differentiation/proliferation cell states for nephron progenitors and shallow/deep states for stroma (**Table S3**). Genes were also pulled from the top variable gene heatmap in the case of the proliferation state. We mapped LOESS-fitted mean *z*-scores of these gene groups to a line radius to create radial plots. Boundary points at continuum scores of 0 and 100 were assigned arbitrarily high weights during fitting to result in closed loops. For each cell compartment in the niche, we created per-cell heatmaps of gene groups by first defining per-gene weights as the inverse of the mean Q3 normalized gene expression across ROIs. ‘Relative expression’ was then represented for each cell as the sum of per-gene weights normalized by a cell’s area. For rhythmic gene heatmaps, the z-score vs. ROI branching continuum data were fitted with a fixed-wavelength sine model for each of the top 1,500 highly variable genes, i.e. *z*-score(continuum) = A.sin(B_fixed_.continuum + C) + D, where A is amplitude, C is phase (radians), D is vertical offset, and B_fixed_ = 2*π* / fitlength is the angular frequency fixed by the user. We chose fitlength = 73 since > 95% of ROIs fell within a branching continuum range of 0-73. A, C, and D were left as fitting parameters. Initial values were set from the data range (A_o_ = (z_max_ - z_min_) / 2, A_o_ = 0, D_o_ = mean(z)). Per-gene models were fitted by non-linear least squares (nls) with a maximum of 1000 iterations. To unify orientation, fits having A < 0 were re-expressed with positive amplitude and phase shifted by *π* radians. For each gene, we computed the residual sum of squares (SSE) for both the sine model and a constant-mean null hypothesis model (i.e. no rhythm in gene expression over branching life-cycle). We then performed an F-test using the statistic:

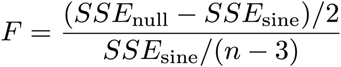

with *n* = number of ROIs, numerator degrees of freedom = 2, and denominator degrees of freedom = *n* - 3. Sine fits of rhythmic genes having *p* < 0.2 were ordered by the branching continuum score of their peak value (‘peak continuum’) and visualized using pheatmap.

### GO/KEGG/reactome gene set and ToppGene transcription factor binding site over-representation analyses of rhythmic genes

We performed a circular sliding-window gene set over-representation analysis on genes ordered by peak continuum. A sliding window of fixed width (25 branching continuum units) was advanced in 3 unit increments. The window was robust to looping of the branching continuum score from 100 back to 0 units. Genes having peak continuum values in the current window were extracted and tested for over-representation in KEGG pathways (enrichKEGG), GO biological process terms (enrichGO), and reactome pathways (enrichPathway). All analyses used Benjamini–Hochberg (BH) multiple-testing correction and default background sets. Significantly over-represented pathways/terms having FDR < 0.05 in the results tables associated with the series of windows were manually inspected and curated for relevance to nephron progenitor/stromal development and mechanics. -log10(FDR) vs. peak continuum window center was then plotted for these curated pathways/terms. We followed a similar approach using the transcription factor binding site analysis of ToppGene for genes in windows 1, 5, 10, and 15 (peak continuum centers of 12.5, 24.5, 39.5, and 54.5). Significantly over-represented binding site IDs having FDR < 0.05 were assigned to naive/shallow, differentiation/deep, or proliferation categories based on inspection of associated gene lists and on the peak continuum range of the window in which they were detected.

### Human iPSC-derived nephron progenitors

Nephron progenitor cell ‘puck’ organoids were generated from SIX2^EGFP^ transgenic reporter iPSC line (*SIX2*-T2A-*EGFP*, Murdoch Children’s Research Institute / Kidney Translational Research Center, Washington University Nephrology) and MAFB^BFP^;GATA3^mCherry^ transgenic reporter iPSC line^139^ according to published protocols^140–142^. Briefly, iPSCs were maintained in standard tissue-culture treated 6-well plates coated with hESC-qualified matrigel (1:100, #354277, Corning) in stem cell maintenance medium plus supplement (mTeSR+ kit, StemCell Technologies 100-0276). Lines were passaged as single cells using Accutase (StemCell Technologies, 07920) once colonies reached 60-80% confluence, and Accutase was neutralized using mTeSR+ at an excess of 6:1 (v/v). For differentiation, maintenance iPSCs were passaged as described above and seeded onto standard tissue-culture treated 6-well plates coated instead with human recombinant laminin 521 (10 µg well^-1^, #77003-05, Biolamina) at a density of 5,200 cells cm^-2^. iPSCs were maintained in mTeSR+ for the first 24 hr ahead of differentiation (day -1). The following day (day 0), wells were washed with DPBS and differentiation towards primitive streak identity was induced using stage 1 media (TeSR-E6 Medium plus supplements (TeSR-E6+) (STEMCELL Technologies, 05946) and 7µM CHIR-99021 (Tocris, 4423)), and stage 1 media was refreshed every 48 hr until day 5. On day 5, wells were washed with DPBS and media was changed to stage 2 media (TeSR-E6+ with 1 µg ml^-1^ heparin (Sigma-Aldrich H4784) and 200 ng ml^-1^ FGF9 (R&D Systems 273-F9-025)). Stage 2 media was refreshed every 24 hr until day 10 to progress differentiation first through an intermediate mesoderm stage before reaching nephron progenitor identity (iNPCs). On day 10, iNPCs were lifted using Accutase and TeSR-E6+ was used to quench digestion at an excess of 3:1 (v/v). iNPCs were resuspended in 1 ml TeSR-E6+ and counted. A volume of iNPC suspension equivalent to 3*10^5^ cells organoid^-1^ was transferred to a 1.5 ml eppendorf and pelleted by centrifuging for 3 min at 300g. Supernatant was then removed and the cell pellet was resuspended with TeSR-E6+ to create a dense cell slurry at ∼3 x 10^5^ cells µl^-1^. Nephron organoids were formed by manually spotting up to eight 1 µl droplets of the slurry onto a single 6-well plate 0.4 µm polyester transwell membrane (CellTreat, 230607) that was prewet with stage 1 media.

Nephron organoids underwent a CHIR ‘pulse’ for 2 hr at 37°C. Organoids were then washed with DPBS and media was exchanged with stage 2 media for a ‘chase’. Nephron organoids were ‘chased’ for 2 hr at 37°C. For select conditions, the CHIR pulse was combined (in pulse media) or staggered (in chase media) with 0.1 µM TTNPB (Tocris 0761), 4 µM TRULI (MedChem Express HY-138489 or Sigma-Aldrich SML3634), or equivolume DMSO (Sigma-Aldrich, 276855). Nephron organoids were washed with DPBS following the ‘chase’ and media was exchanged with stage 2 media. Stage 2 media was refreshed after 24 hr. On day 12, nephron organoids were washed with DPBS and the media was switched to TeSRE6+ without additional factors. TeSR-E6+ was refreshed every 48 hr until reaching the experiment endpoint.

### Organoid immunofluorescence

Organoids were fixed in 4% paraformaldehyde for 15 min. Fixed cells were then washed twice for 10 min in PBSG (DPBS + 7.5 g/L glycine) and once in DPBS. Organoids were permeabilized in 0.5% Triton-X-100 for 30 min at room temperature and blocked for 1hr at room temperature in IF Wash (DPBS + 1g/l Bovine Serum Albumin + 0.2% Triton-X-100 + 0.04% Tween-20) + 10% donkey serum. Organoids were then incubated overnight at 4°C in appropriate dilution of primary antibody in IF Wash + 10% donkey serum, washed three times for 1 hr in IF wash at room temperature, and incubated in secondary antibody at overnight at 4°C in IF wash + 10% donkey serum. Organoids were again washed three times in IF wash for 1 hr. Organoids were counterstained in 1 µg ml^-1^ DAPI in DPBS and washed once in DPBS before imaging using confocal microscopy (see **Kidney immunofluorescence imaging)**. Primary antibodies and dilutions were rabbit anti-SIX2 (1:400, 11562-1-AP, Proteintech, RRID: AB_2189084), rabbit anti-ECAD (1:300, 3195, Cell Signaling Technology, RRID: AB_2291471), goat anti-JAG1 (1:150, AF599, R&D Systems, RRID: AB_2128257), goat anti-GATA3 (1:20, AF2605, R&D Systems, RRID: AB_2108571), sheep anti-NPHS1 (1:40, AF4269, R&D Systems, RRID: AB_2154851), mouse anti-YAP (1:50, sc-101199, Santa Cruz, RRID: AB_1131430), and biotinylated LTL (1:300, B-1325-2, Vector Laboratories). Donkey secondary antibodies were used at 1:300 dilution: anti-rabbit AlexaFluor 647 (A31573, ThermoFisher, RRID: AB_2536183), anti-goat AlexaFluor 488 (A11055, ThermoFisher, RRID: AB_2534102), anti-sheep AlexaFluor 555 (A21436, ThermoFisher, RRID: AB_2535857), and DyLight 405-Streptavidin (016-470-084, Jackson ImmunoResearch).

Day 10 organoids were fixed 2 or 4 hr after spotting and stained as described above. FIJI was used to manually segment nephron progenitor (SIX2+) nuclei using the DAPI channel and cytoplasms by extending the outer DAPI edge to the midway point of neighboring nuclei. Mean YAP intensity was measured for each ROI and background subtracted using image-wide mean YAP intensity at a distant, out of focus plane. Background subtracted averages were used for calculating YAP nucleus:cytoplasmic ratio.

For day 12 organoids, nephron progenitor (SIX2+) and early nephron (JAG1+) % area relative to DAPI+ area was determined by thresholding *z*-stack montages in FIJI. For day 25 organoids, glomeruli (NPHS1+) % area was determined by thresholding *z*-stack montages while proximal tubule (LTL+) % area, distal tubule (only ECAD+) % area, and connecting segment (nuclear GATA3+ and ECAD+) % area relative to organoid area (determined by manual annotation of background signal) were determined by manual annotation of z-stack montages followed by thresholding. Analyses were performed on a single, central z-plane for each organoid.

### Organoid RNA isolation and quantitative PCR

Organoids were collected for RNA extraction with 8 organoids pooled for each condition. Following the manufacturer protocols, we used the RNeasy Mini Kit (Qiagen 74104) to isolate RNA and the High-Capacity RNA-to-cDNA Kit (Applied Biosystems 4387406) to generate cDNA libraries. For qPCR, we used the PowerUp SYBR Green Master Mix (Applied Biosystems A25742). We used an Applied Biosystems 7300 thermocycler set to manufacturer suggested times and temperatures for PowerUp SYBR Green Master Mix. 25 ng of cDNA were used in each reaction along with the appropriate qPCR primers at 500 nM. qPCR primers for *CRABP2*, *CTGF*, *CYP26A1*, *CYR61*, *GAPDH*, *HPRT*, and *RARB* were used (**Table S4**). DeltaDeltaCt values were calculated using either *GAPDH* or *HPRT* as housekeeping genes. Samples were run in triplicate for qPCR analysis.

### Single cell sequencing prep

E17 wild-type CD1 kidneys were pooled to generate NZC pellets (see **Mouse kidney primary nephrogenic zone cell isolation**). NZC cell pellets were resuspended to 10^6^ cells ml^-1^ in 1x DPBS, and cell suspensions were kept on ice until cDNA library preparation. cDNA library preparation, quality control, and sequencing were performed with help from Children’s Hospital of Philadelphia’s Single Cell Technology (identifying info) and High-throughput Sequencing (identifying info) Cores. Single-cell RNA-seq was performed using the Chromium GEM-X Single Cell 3’ Kit v3 (10x Genomics) following the manufacturer’s protocol. In brief, approximately 10,000 cells were captured per sample and partitioned into Gel Bead-in-Emulsions (GEMs), where cell lysis and barcoded reverse transcription of mRNA occurred. cDNA was then amplified, fragmented, and subjected to 3’ library construction with unique sample indices. cDNA libraries were generated in bulk and quality control was performed using the High Sensitivity DNA Kit (Agilent) and 2100 bioanalyzer (Agilent) ahead of sequencing. Once quantified, the dual-indexed libraries were diluted, pooled in equal molarity, and sequenced in Illumina NovaSeq6000 on a SP-100 flow cell (Illumina). Paired-end sequencing was performed as per 10X Genomics recommendations as 28x10x10x90. The pool was denatured and diluted to a loading concentration of 350pM, and PhiX control was added at 1%; cluster PF was recorded at 76% and Q30 > 93%.

### scRNA-seq data analysis

#### Mouse E17 NZC feature count matrix generation

The Kallisto-Bustools workflow (kb-python; Kallisto version 0.50.1 and Bustools version 0.40.0) was used to index pseudo-alignment and paired-end read FASTQ transcript mapping^143^. The ‘kallisto index’ call was used for index generation from the mouse reference transcriptome cDNA fasta file (Mus_musculus.GRCm39.cdna.all release-111; ensemble.org). The default kb-python mouse transcript to gene map was downloaded using the ‘kb ref’ call with the default (-d) argument set to ‘mouse’. The ‘kb count’ call was used for collapsing reads to unique molecular identifiers (UMI), mapping UMIs to transcripts and cells, and generating a cell x feature matrix. Input for the ‘kb count’ call were Illumina read 1 and read 2 FASTQ files, the mouse reference transcriptome index, default kb-python transcript to gene map, capture technology specified as 10x version3 (‘10XV3’), and output format specified as ‘cellranger’. The default kb-python Chromium 10x-v3 capture technology whitelist was used for barcode filtering.

#### Seurat object pre-processing

Empty droplets were estimated from the output cell x feature count matrix using the *emptyDrops* function from the DropletUtils package (v1.24.0, Bioconductor DOI:10.18129/B9.bioc.DropletUtils). Cells with scores less than or equal to 0.05 were kept for downstream analysis. In short, the Seurat library (v5.2.1)^144,145^ was used to import the filtered gene expression matrix to R and for downstream analysis. Seurat objects were filtered to exclude features in fewer than 3 cells and cells with over 20% of transcripts assigned to mitochondrial RNA, fewer than 1000 unique features, and greater than 20,000 transcript counts. The filtered dataset contained 9530 cells with an average of 3144 unique genes and an average of 9300 total counts detected per cell.

Gene expression matrices generated from scRNA-seq of isolated nephrogenic zone cells from two week 17 human fetal kidneys were used for integration with human week 20 Xenium datasets^40^. These matrices were accessed from the Gene Expression Omnibus (GEO, NCBI) under accession code GSE112570. Seurat objects were filtered as described in *Lindstrom et al*^40^. In brief, cells with fewer than 1000 unique features, greater than 5% of transcripts assigned to mitochondrial RNA, and with Good-Turing estimates of observed expression (S) under 0.7 were filtered out. The Good-Turing estimate is described by S = 1−n_1_/N, where n_1_ is the number of genes with one mapped read and N is the total number of reads in the cell. Seurat objects from kidneys 1 and 2 were then merged ahead of downstream analyses with the filtered dataset containing 7350 cells with an average of 1966 unique genes and an average of 6386 total counts detected per cell. The Seurat library *Read10X_h5* function was used to load Xenium output cell count matrices with the corresponding ‘Gene Expression’ matrix being used as input for each Seurat object. Seurat objects were filtered by removing cells with fewer than 50 unique features and removing features in fewer than 100 cells. Additionally, cells with greater than 2 identified nuclei based on Xenium segmentation were removed.

#### Dataset Integration

Mouse and human nephrogenic zone scRNA-seq datasets were merged with Xenium mouse E17 slide 2 and human week 20 slide 1, respectively. Merged datasets were then processed with the following Seurat functions using default arguments: *NormalizeData*, *FindVariableFeatures*, *ScaleData*, and *RunPCA*. Dataset integration was performed through canonical correlation analysis (CCA) on the joint PCA latent space using Seurat’s *IntegrateLayers* function.

*Cluster Identification and QC*: Following integration, the first 30 dimensions of the CCA latent space were used as input for Seurat’s *FindNeighbors* function to generate a shared nearest neighbor (SNN) graph. Seurat’s *FindClusters* function was then used to identify clusters at resolutions of 0.5 and 0.3 for the mouse and human datasets, respectively. Resolutions were selected based on clustering of expected cell types in the shared datasets using a manually curated panel of cell type marker genes found in the Xenium panel (**Fig.S25/26**). Datasets were then separated and re-processed with *FindVariableFeatures* and *ScaleData* functions ahead of cluster DGE analysis. Wilcoxon rank sum test was used for DGE analysis using Seurat’s *FindAllMarkers* function. Sub-cluster identities were defined after manual exploration or marker genes with published nephron and stromal lineage datasets from Lawlor *et al.^42^* and England *et al.^53^*, respectively. To refine cell type identities further, stromal and nephron progenitor sub-clusters were then re-integrated and re-clustered while isolated nephron and ureteric bud clusters were re-clustered without further integration due to low cell number mapping to these identities in the nephrogenic zone cell datasets. Refined identities were determined using previously published references.

Clusters were screened with several quality control metrics prior to downstream analysis: doublet probability was assessed by the *computeDoubletDensity* function from the scDblFinder package (v1.18.0, Bioconductor DOI: 10.18129/B9.bioc.scDblFinder), while mean cell area, nucleus area, and mean transcript count were obtained from Xenium output parquet files. Clusters 7 and 9 from the initial mouse integration analysis were filtered out after QC screening; cells from cluster 7 showed consistently high doublet probability scores with marker gene expression indicating contribution of both NP and stromal lineages while cells from cluster 9 were considerably smaller and had fewer transcripts than remaining clusters (**Fig. S25**). No clusters were removed from the human xenium analysis (**Fig. S26**).

#### Niche identity analysis

Cells from depth-filtered niches were used for niche composition analyses, and niche assignments were kept consistent with the pseudo-bulked analysis described above. Sub clusters with insufficient cell numbers contributing to the total depth-filtered population were additionally removed from the analysis. First, UMAPs were generated for each general cell type (NP, stroma, and UB) after default re-processing with kidney slice identity being regressed out during gene expression scaling for mouse datasets. A custom function to measure scatter plot point density (https://slowkow.com/notes/ggplot2-color-by-density/) was used to plot relative UMAP density. Density was measured as the number of cells occupying each grid divided by the total number of cells in the plot for a uniform 10 by 10 grid along UMAP dimensions 1 and 2. Chi-squared testing was used to evaluate similarity of cell type composition associated with discrete life-cycle stages. Wilcoxon rank rum testing with Bonferroni Hochberg multiple comparison adjustment was used to evaluate change in individual niche composition associated with discrete life-cycle stages. The ureteric bud population was not confined to the two cell layer adjacent to the cap mesenchyme as was the case in the pseudo-bulked analysis described above.

#### Deep vs Shallow Gene Set Scoring

Clusters 1 and 4 from Xenium’s output k-means clustering (k=4) from the mouse E17 slide 1 dataset were used as input for Seurat’s *AddModuleScore* function. A shallow gene set score was generated with nbin set to 3, ctrl set to 100, seed set to 1, and features set with the following genes: *Foxd1*, *Ntn1*, *Smoc2*, *Sfrp1*, *Cyp1b1*, *Arap1*, *Adgrl2*, *Igfbp5*, *Nr2f1*. A deep gene set score was generated using consistent arguments except for features set with: *Pdgfrb*, *Id2*, *Snai2*, *Cxcl12*, *Dkk1*, *Pcna*, *Shisa3*, *Bmper*, *Fzd1*, *Peg10*, *Dlc1*. Clusters 1,2,4,7,10, and 11 from the initial cluster identification of the human wk20 Xenium slide 1 dataset were used as input for shallow and deep gene set analyses as described above. The human shallow geneset overlapped with the mouse shallow geneset with *Nr2f1* removed and *TGFBI*, *SULF2*, and *DNAH11* added. The human deep geneset overlapped with the mouse deep geneset with *Shisa2*, *Fzd1*, *Peg10*, and *Dlc1* removed and *ID3*, *IRS4*, *THBS1*, *GATA6*, and *TNC* added. Human genesets were type adjusted to match human gene annotations. Cells were assigned ‘shallow’ classification if the cell’s shallow score was greater than its deep score, assigned ‘deep’ classification if the cell’s deep score was greater than its shallow score, and assigned ‘intermediate’ classification if shallow and deep scores were the same. Classification metadata was input into Xenium Explorer 3 for inset generation (**Fig. 5F**, **Fig. S22F**).

#### CONCORD

Python (v3.12) was used to run CONCORD (v0.9.5, ref. ^73^), a recently developed scRNA-seq integration software that uses contrastive learning to identify biologically meaningful structures in high dimensional data, for mapping the mouse E17 nephrogenic zone cell dataset and to integrate the Xenium human wk20 slide1 dataset with human wk17 nephrogenic zone cell dataset. Seurat objects were first converted to single cell experiment (SCE) objects in R using Seurat’s *as.SingleCellExperiment* function. The anndata2ri (v1.3.2, https://github.com/theislab/anndata2ri) and rpy2 (v3.5.17, https://pypi.org/project/rpy2/) packages were used in the python environment to import and convert SCE objects to python compatible anndata objects. CONCORD was run using default settings in both instances. For the mouse E17 NZC run, the top 5000 variable genes were identified by running *select_features* function with flavor set to ‘seurat_v3’ and included in the mapping. For the human dataset integration, all overlapping genes were included in the mapping and the domain_key argument was set to distinguish the datasets. The *run_umap* function with arguments n_components set to 3, n_neighbors set to 30, min_dist set to 0.1, and metric set to ‘euclidean’ was used on CONCORD dimensions 1-32 to obtain a 3-dimensional UMAP reduction for the mouse mapping. Output anndata objects were then saved as h5ad files using the anndata*.write* function. The readH5AD function (zellkonverter v1.14.1, <u>theislab.github.io/zellkonverter/</u>) was used to load and convert anndata objects into R as SCE objects and Seurat’s *as.Seurat* function was used to convert SCE objects into seurat objects. Nephron progenitor and nephron clusters were isolated for plotting. The Xenium dataset was additionally isolated for the wk20 human plot and a new 3D UMAP reduction was generated from the latent CONCORD dimensions 1-32 using *RunUMAP* with default parameters and n.components set to 3. Cell cycle phase identification was determined by the *CellCycleScoring* function with S and G2M gene sets taken from Seurat’s cc.genes.updated.2019 list. The *plot_ly* function (plotly v4.10.4, https://plotly.com/r/) set with type ‘scatter3d’ was used to generate interactive 3D scatter plots using CONCORD UMAP dimensions 1-3 and colored either by refined cluster ID or cell cycle phase assignment. Plots were manually oriented to highlight relevant angles of the 3D UMAP space.

#### Cell-cell communication ligand-receptor database generation

CellChat^7^ (v2.2.0) was used for inferential cell-cell communication analysis of depth-filtered cells from mouse E17 (slide 2) and human week 20 (slide 1) datasets. Clusters with insufficient cell numbers contributing to the overall population were removed prior to analysis. Additionally, niches that did not include stroma were removed from the analysis. Default mouse and human CellChat ligand-receptor databases were updated to include non-overlapping L-R pair annotations from the celltalk database (v1.0, https://github.com/ZJUFanLab/CellTalkDB) for mouse and celltalk and cellphone (v5.0.0, https://www.cellphonedb.org/) databases for human CCC analyses. L-R pairs added from the CellTalk and CellPhone databases had column names changed to be consistent with the CellChat database and additional columns removed prior to merging. All added CellTalk L-R pairs were given an associated pathway name of “NA_celltalkDB.interaction” and signaling annotation of “Secreted Signaling”.

Pathway names for L-R pairs added from the CellPhone database were taken from the classification column adjusting to match the style and naming provided by CellChat. Pathways without an associated classification were given an associated pathway name of “NA_cellphoneDB.interaction”. CellPhoneDB L-R pairs were given a signaling annotation of “Secreted Signaling”. New L-R pairs that overlapped between the CellTalk and CellPhone databases were taken from the CellPhone database due to added pathway association information. The final mouse L-R database included 4505 pairs (3379 and 1126 from CellChat and CellTalk, respectively) and the final human L-R database included 6841 pairs (3233, 1596, and 2012 from CellChat, CellPhone, and CellTalk, respectively).

#### Within dataset cell-cell communication inference

Separate cellchat objects were generated for cells associated with each discrete life-cycle stage by subsetting the Xenium seurat objects according to depth status and discrete life-cycle stage annotation of the assigned niche. Cellchat objects were grouped by refined cluster identities, had datatype set to ‘spatial’, had the coordinates argument set with a matrix comprising of each cell’s centroid x and y coordinates acquired from the Xenium output cells.parquet file, and the spatial factors argument set with a data frame containing a ratio variable equal to 1 indicating Xenium measurements are in µm and a tolerance variable equal to 5 indicating that cell boundaries are 5 µm radially from the centroid. The *subsetData* function was used to remove genes not found in the L-R database from the analysis. The *identifyOverExpressedGenes* function used Wilcoxon rank sum testing with Bonferroni multiple comparisons adjustment to find differentially expressed genes between all cell type clusters within each dataset. The *identifyOverExpressedInteractions* found significant L-R pairs between all sender-receiver combinations as defined by the ligand being significantly upregulated in the sender population and receptor being significantly upregulated in the receiving population. Due to the low sequencing depth of the Xenium datasets, the *smoothData* function was employed to smooth gene expression based on protein neighbors in experimentally mapped protein-protein networks provided by CellChat. The *computeCommunProb* function was used to identify significantly expressed L-R interactions between sender and receiver pairs within the dataset. The computeCommProb function was set with the following arguments: type set to “truncatedMean”, trim set to 0.1, raw.use set to false, population.size set to true, distance.use set to true, interaction.range set to 50 µm, scale.distance set to 5, contact.dependent set to true, contact.range set to 10, and nboot set to 500. The *filterCommunication* function with min.cells set to 10 was used to remove communications occurring in fewer than 10 cells per sender/receiver cluster pairing.

#### Differential cell-cell communication analysis

CellChat objects generated as described above were merged for differential cell-cell communication analyses using the *mergeCellChat* function (mouse E17 pre- vs post-branch and human wk20 pre- vs mid-branch). The *updateClusterLabels* function was used to arrange cluster positions on circle plots and *subsetCellChat* function was used to reduce analyses to sender and receiver clusters of interest. Circle plots showing net differential interaction weight between sender and receiver cluster pairs were generated with the *netVisual_diffInteraction* function with the measure argument set ‘weight’ and the sources.use and targets.use arguments set depending on analysis (**Fig.4F**, **Fig.S18F**, **Fig.S19E**, **Fig.S20E**). A custom built function was used to identify differential interaction weight between specific L-R pairs across all sender-receiver cluster pairs analyzed. In brief, the *subsetCommunication* function was used on the “net” slot to isolate L-R interaction weights in all sender-receiver cluster pairs per branching stage. The *full_join* function (dplyr) was used to rejoin the branching stage “net” slots on all columns minus L-R pair ‘prob’ and ‘pval’ columns which were kept separate using dataset identifiers. Differential interaction weight was calculated for individual L-R interactions within each sender-receiver cluster pairing by using the *group_by* function (dplyr) on sender and receiver clusters and then subtracting prob_dataset2 from prob_dataset1. L-R interactions not found in one dataset had probability values set to 0 ahead of measurement.

#### Human week 20 nephron progenitor stream analyses

The full human wk20 slide 1 dataset was used for nephron progenitor analyses. Cell cycle phase identification was determined by the *CellCycleScoring* function with S and G2M gene sets taken from Seurat’s cc.genes.updated.2019 list. Centroid to centroid euclidean distances were measured using coordinates provided by Xenium cells.parquet output files for all cell pairs between testing sets. Refined NP clusters were evaluated against targets from refined UB and nephron clusters. Matrices were collapsed by finding the minimum distance to a cell of the target identity (column) for each NP cell (row). The minimum distance matrix was further filtered to remove distances greater than 15 µm or 75 µm depending on the analysis (**Fig.S21**).

### Stromal nuclear morphology analysis

For week 20 human, the Xenium ‘Level 1’ DAPI morphology image was processed in FIJI using in-house macros; breaking this image into 512 x 512 pix tile stacks and finding in-focus planes for each stack using Focus LP^146^ (Helmut Glunder). In-focus planes were then montaged and all nuclei segmented using Cellpose^147^ (v4.0.7) after human-in-the-loop model training on 5 representative in-focus planes spanning the *xy* extent of the montage. Nuclei masks were exported to FIJI for aspect ratio and area calculation. These morphology data were then associated with Xenium cell IDs through a nearest neighbor search based on Euclidean distance between cell centroids in R (FNN package). Running average aspect ratio was quantified for a window size of 100 cells after ordering cells by their centroid *x* coordinate. For E17 mouse, variation in stromal nuclear morphology was not as large in Xenium slices from kidneys fixed in the sandwich system as in traditionally fixed kidneys, perhaps due to differences in PFA penetration vs. cell relaxation timescales. We therefore segmented nuclei in whole-mount immunofluorescence images of freshly dissected E17 kidneys using the same Cellpose approach as above. In this case, stromal cells were manually associated with the nearest cap mesenchyme population and duplicate associations were allowed in the case that a stromal cell contacted two caps. Despite its lower dynamic range, Cellpose segmentation of the Xenium DAPI morphology image was still used in analyses requiring correlation of gene expression and mouse stromal nuclear aspect ratio (see below). Peripheral stromal cells above niches were directly segmented on a per-niche basis from whole-mount image stacks in mouse. This was not possible for human since surface slices containing the same set of niches corresponding to those in the Xenium slices were not available. Instead, peripheral stromal nuclei at slice edges were analyzed to approximate their average aspect ratio and area values. Per-cell aspect ratio heatmaps were generated by shading Xenium and Cellpose nuclear outlines for human and mouse respectively using in-house code. Per-niche pseudobulked Q3-normalized gene expression was fit to per-niche average stromal nuclear AR via a linear model to determine R^2^ values for each gene. Data sorted by R^2^ were split into genes with positive or negative correlations, and genes having R^2^ > 0.05 in each category were analyzed using ToppGene pathway and transcription factor binding site analyses. Significantly over-represented pathway/binding site IDs having FDR < 0.05 were manually inspected and curated for relevance to nephron progenitor/stromal development and mechanics.

### Mouse kidney stromal pMLC, phalloidin, and YAP analyses

pMLC/phalloidin fluorescence vs. depth in mouse embryonic kidney stromal ribbons was quantified from average projections of xz-plane confocal sections (i.e. orthogonal to the *en face* plane) local to each ribbon at 0.2 µm spacing. Fluorescence vs. depth traces were background-subtracted using the fluorescence at the shallowest point in depth, which was outside the tissue boundary. These traces were normalized by their value at an anatomical reference position, namely the *z* plane glancing the cortical surface of the ureteric bud tips adjacent to a given stromal ribbon. A similarly-processed trace for an average projection of xz-plane sections spaced at 2 µm spanning the entire confocal field of view was then subtracted from each ribbon trace to yield an ‘enrichment’ metric. The enrichment traces were aligned to the same reference position across niches. These steps correct for decreasing fluorescence with depth due to light scattering and quantify any pMLC enrichment in ribbons that is higher/lower than expected on average at a given depth across ureteric bud/nephron progenitor/stromal compartments.

YAP nuclear:cytoplasm ratio did not require the same normalization procedure since the per-cell ratio intrinsically accounts for decreasing fluorescence with tissue depth. Cellpose was used on the DAPI channel for nuclei segmentation. Niche ROIs were manually generated in FIJI using the polygon selection tool, and *xy* coordinates from each niche’s vertices were written into csv files for cell-niche association in R. A custom FIJI macro was used to filter Cellpose output segmentations, only keeping nuclei having partial overlap with binarized masks generated from niche ROIs. Remaining nuclei ROIs were loaded as objects into CellProfiler using the loadmaskobjects.py plugin (Egor Zindy). Cytoplasms were defined as annuli extended up to 3 pixels around each nucleus and stopping at neighboring cytoplasms. Binarized niche masks were also input into CellProfiler and converted back into individual niche objects using a watershed threshold. Nuclei centroids were used to relate cells with niches. YAP intensity, MEIS123 intensity, and shape parameters measured in nucleus, cytoplasm, and niche objects were output from CellProfiler. CellProfiler output data was analyzed and plotted in R. In brief, the st package (v 1.0.19, https://r-spatial.github.io/sf/) was used for cell-niche association by measuring nuclei centroid occupancy within a niche’s boundary. FIJI output coordinates were used to define niche boundaries for cell assignments. Nuclei occupying multiple niches were assigned to each niche.

Stromal cells were detected by setting a manual threshold on MEIS1/2/3 marker fluorescence for every z plane. Cells having median nuclear intensity at or above the threshold and a nucleus:cytoplasmic median MEIS1/2/3 intensity ratio above 1.25 were classified as ‘stroma’. YAP background was calculated on a per niche basis as the niche’s mean intensity minus two standard deviations. Nuclear and cytoplasmic YAP intensities were background subtracted using niche assignments determined in CellProfiler prior to nucleus:cytoplasmic ratio measurements. Stromal cells within the first or last 2.5 percentiles for nucleus area were removed. Remaining cells with a cytoplasmic:nucleus area ratio under 0.2 were additionally removed prior to analysis.

### Mouse kidney primary nephrogenic zone cell isolation

Wild-type CD1 E17-E18 embryonic kidneys were dissected, pooled per litter, and kept on ice in DPBS until digestion. For NZC single cell RNA-seq, a digestion solution made up of 0.25% pancreatin and 0.1% collagenase in DPBS was applied for 15 min in a cell culture incubator set to 37°C and 5% CO_2_. Samples were placed on a shaker at 100 rpm for constant agitation. Digestion was quenched using FBS and released DNA was digested using 6 units of TURBO DNase (2 U/μl). Cell suspensions were transferred to 1.5 ml eppendorf tubes and centrifuged at 300g for 5 min. Cells were then prepared for sequencing (see *single cell sequencing prep*). For NZC cell substrate adhesion assays, a digestion solution of 0.25% trypsin-EDTA (Invitrogen, 25200056) was applied up to 30 minutes in a cell culture incubator set to 37°C and 5% CO_2_. Samples were placed on a shaker at 100 rpm for constant agitation. Every 10 minutes, solutions were triturated with a p1000 taking care not to suck up kidneys, cell suspensions transferred to an eppendorf containing FBS at 10% digest volume for quenching trypsin, and a new volume of 0.25% trypsin-EDTA was added to the kidneys for continued digestion. Quenched samples were kept on ice until 30 minutes elapsed. After all cell suspensions were collected, cells were centrifuged at 300g for 5 min. Cells were then prepared for substrate adhesion assays (see *Mouse kidney primary cell substrate adhesion immunofluorescence and qPCR/bulk RNA-sequencing)*.

### Mouse kidney primary cell substrate adhesion immunofluorescence and qPCR/bulk RNA-sequencing

NZC cell pellets were washed by resuspension in FACS media (DPBS with 2% FBS, 1% pen/strep and 10 µM Y27632), re-pelletted, and then resuspended at 10^6^ cells 200 uL^-1^ in Alexa Fluor 488 conjugated goat anti-PDGFRA (1:200, FAB1062G, R&D Systems, RRID:AB_3645435) diluted in FACS media. Cell suspensions were incubated on ice for 1 hr, washed with FACS media, and resuspended at 10^6^ cells 200 uL^-1^ in donkey anti-goat AlexaFluor 647 (1:200, A32849, ThermoFisher, RRID: AB_2762840) diluted in FACS media. Cells were incubated in secondary for 1 hr on ice and then washed with FACS media. Labeled cells were passed through a 40 µm cell strainer and enriched using BD FACSAria Fusion (BD Biosciences). The PDGFRA positive fraction were seeded at 2.5 x 10^5^ cells well^-1^ of a 12-well polystyrene plate with one 18mm PDMS coated glass coverslip per well. PDMS was adsorbed with either 50 µg ml^-1^ (high-adhesion) or 10-500 ng ml^-1^ (low-adhesion) bovine plasma fibronectin (F1141-5MG, Sigma) for at least 3 hr in a tissue culture incubator set to 37°C and 5% CO_2_. Cells were grown in DMEM with 10% FBS and 1% pen/strep. After 14-18 hr, cells were either prepared for immunofluorescence or bulk-sequencing analyses. The rounding/clustering phenotype observed for the low-adhesion immunofluorescence analysis was lost in subsequent experiments using the same concentration of fibronectin (500 ng ml^-1^). However, the phenotype was regained after a further reduction in fibronectin concentration (10 ng ml^-1^), and this reduced concentration was used for follow-up bulk-sequencing experiments.

#### Immunofluorescence

Cells were fixed using 4% paraformaldehyde for 5 min at room temperature. Fixed cells were then washed twice in PBSG for 5 min and once in DPBS. Samples were incubated for 1 hr in IF wash with 5% donkey serum for permeabilization and blocking. Primary and secondary incubation was overnight in IF wash + 5% donkey serum. Samples were washed for 1 hr in IF wash following antibody incubation. Samples were counterstained in 1 µg ml^-1^ DAPI in DPBS and washed once in DPBS before imaging using confocal microscopy (see *Kidney immunofluorescence imaging*).

Primary antibodies and dilutions were rabbit anti-YAP (1:200, PA1-46189, Invitrogen, RRID: AB_2219137), mouse anti-ɑ-Smooth Muscle Actin Alexa Fluor 488 (1:300, 53-9760-82, Thermo Fischer Scientific, RRID:AB_2574461), and goat anti-PDGFRA (1:200, AF1062, R&D Systems, RRID:AB_2236897). Donkey secondary antibodies were used at 1:300 dilutions: anti-rabbit Alexa Fluor 555 (A32794, Thermo Fisher Scientific, RRID:AB_2762834) and anti-goat Alexa Fluor 647 (A32849, Thermo Fisher Scientific, RRID:AB_2762840).

#### bulk RNA-sequencing

RNA was kept at -80°C immediately after isolation (see *Organoid RNA isolation and quantitative PCR*) until all samples were processed. RNA was subjected to concentration measurement, cDNA library preparation, and mRNA poly-A capture 3’ bulk sequencing through a commercial service (Plasmidsaurus) using Illumina NovaSeq sequencing and 3’ end counting. Quality of the resulting fastq files was assessed using FastQC v0.12.1. Reads were then quality filtered using fastp v0.24.0 with poly-X tail trimming, 3’ quality-based tail trimming, a minimum Phred quality score of 15, and a minimum length requirement of 50 bp. Quality-filtered reads were aligned to the reference genome using STAR aligner v2.7.11 with non-canonical splice junction removal and output of unmapped reads, followed by coordinate sorting using samtools v1.22.1. PCR and optical duplicates were removed using UMI-based deduplication with UMIcollapse v1.1.0. Alignment quality metrics, strand specificity, and read distribution across genomic features were assessed using RSeQC v5.0.4 and Qualimap v2.3, with results aggregated into a comprehensive quality control report using MultiQC v1.32. Gene-level expression quantification was performed using featureCounts (subread package v2.1.1) with strand-specific counting, multi-mapping read fractional assignment, exons and three prime UTR as the feature identifiers, and grouped by gene_id. Final gene counts were annotated with gene biotype and other metadata extracted from the reference GTF file. Sample-sample correlations for sample-sample heatmap and PCA were calculated on normalized counts (TMM, trimmed mean of M-values) using Pearson correlation. Differential expression analysis was performed with edgeR v4.0.16 using a paired analysis by mouse litter, including filtering for low-expressed genes with edgeR::filterByExprwith default values. Density plots along the logFC axis were determined for the top 7000 variable genes using kernel density estimation.

### Mouse kidney explant adeno-associated virus knockdown and validation in NIH/3T3 cells

Adeno-associated virus serotype 2 (AAV2) vectors were commercially produced and concentrated to 10^13^ genome copies ml^-1^ by VectorBuilder. AAV2 vectors contained two miR30 short hairpin siRNA sequences downstream of a CAG promoter and mCherry fluorescent reporter. The miR30 shRNA sequences were designed to target two regions on non-muscle myosin light chain kinase (*Mylk*) (AAV2-mCh-shMylk), one region on Rho Associated Coiled-Coil Containing Protein Kinase 1 (*Rock1*) and one region on Rho Associated Coiled-Coil Containing Protein Kinase 2 (*Rock2*) (AAV2-mCh-shRock1/2), or randomized Rock1 and Rock2 target sequences (AAV2-mCh-shScramble).

#### 3D explant transduction and live nephrogenesis tracking

Explant cultures differed from the protocol described earlier (see *3D mouse embryonic kidney explant culture*) in the following ways: prior to gelation, kidneys were transfected by incubating for 3-4 hr at 37°C in DMEM with 10% FBS, 1x pen/strep, ∼2 x 10^12^ GC ml^-1^ of appropriate AAV2-mCh-shRNA and live labeled anti-EpCAM/CD326 FITC (1:250, 11-5791-82, Invitrogen). Post transduction, kidneys were washed using DMEM with 10% FBS and 1x penicillin/streptomycin. Kidneys from the same condition were grouped for imaging with up to 9 kidneys used per well on a 6-well glass coverslip bottom plate (80637, Ibidi). Kidneys were encapsulated individually without the use of a PDMS ring and PBS to maintain humidity during gelation. Kidneys were re-stained using anti-EpCAM/CD326 FITC (1:250) in DMEM with 10% FBS and 1x pen/strep for 1-2 hr at 37°C followed by 1-2 hr wash in DMEM with 10% FBS and 1% pen/strep 16-24 hr post-encapsulation. Media was then exchanged with phenol red-free DMEM media with 10% FBS and 1x pen/strep prior to imaging as described in 3*D mouse embryonic kidney explant culture*. Nephron initiation was annotated as in **Live nephrogenesis tracking** above.

#### Transduced kidney dissociation and RNA isolation

After 72 hr 3-9 kidneys per condition were transferred to a 1.5 ml eppendorf for cell isolation. Samples were washed with DPBS and digested using a 1:1 (v/v) mixture of 1% type I collagenase (Sigma-Aldrich, C0130-100MG) and 0.25% trypsin-EDTA (Invitrogen, 25200056). Digestion was carried out in a cell culture incubator set to 37°C and 5% CO_2_ up to 30 min with intermittent trituration every 10 min. Samples were placed on a shaker at 100 rpm for constant agitation. Digestion was quenched using FBS. Cell suspensions were pelleted by centrifuging at 300g for 5 min. Cells were resuspended in FACS media (DPBS with 2% FBS and 1% pen/strep) and then enriched through FACS into mCherry-positive and mCherry-negative populations (top 25% and bottom 50% of mCherry signal, respectively). Positive and negative fractions were then pelleted, washed with DPBS and pelleted a second time ahead of RNA isolation. RNA isolation was performed using a Qiagen RNeasy kit following manufacturers instructions. RNA was kept at -80°C immediately after isolation.

#### Bulk RNA-sequencing

RNA was subjected to mRNA poly-A capture 3’ bulk sequencing through a commercial service (Plasmidsaurus) using Illumina NovaSeq sequencing, 3’ end counting, and data processing as in **Mouse kidney primary cell substrate adhesion immunofluorescence and qPCR/bulk RNA-sequencing**. Outcomes are consistent with analyses using manual fastq mapping and transcript abundance quantification through kallisto. The mouse reference transcriptome (Mus_musculus.GRCm39.cdna.all release-111; ensemble.org) with the mCherry cDNA sequence (GenBank Accession #AY678264.1) manually appended was used to build a transcript to gene index through Kallisto *index* call. The custom built index was then used for transcript to gene abundance estimates using the kallisto *quant* call with optional arguments set for single-end reads, estimated average transcript length of 250, and estimated transcript length standard deviation of 30.

#### 3D explant cryopreservation and immunofluorescence

Tissues were incubated in 30% sucrose for 24 hr, embedded in OCT, frozen on dry ice, and stored at -80°C until sectioning. Cryosections were collected at 20 µm thickness. For staining, slides were air-dried for 60 minutes at room temperature and all subsequent steps were performed in a humidified chamber. Sections were blocked for 1 h in 1x immunofluorescence (IF) wash buffer containing 5% donkey serum, followed by overnight incubation at 4°C with primary antibodies diluted in the same blocking buffer. Slides were washed three times for 10 min each in 1x IF wash at room temperature, incubated with secondary antibodies for 2 h at 37°C, and washed again three times for 10 min. Primary antibodies included mouse anti-SIX2 (1:400, 66347-1-Ig, Proteintech, RRID:AB_2881727), goat anti-PDGFRA (1:400, AF1062, R&D Systems, RRID:AB_2236897), and rabbit anti-RFP (1:1000, ab62341, Abcam, RRID:AB_945213). Secondary antibodies (raised against donkey and used at 1:500 dilutions) included anti-mouse AlexaFluor 488 (A21202, ThermoFisher, RRID:AB_141607), anti-rabbit AlexaFluor 555 (A31570, ThermoFisher, RRID: AB_2536180),, and anti-goat AlexaFluor 647 (A32849, ThermoFisher, RRID:AB_2762840).Nuclei were counterstained with 2.5 µg ml^-1^ DAPI in DPBS for 10 min, rinsed once in DPBS, and mounted in 85% glycerol / 15% deionized water ahead of confocal imaging using a 20x objective (see **Kidney immunofluorescence imaging**).

#### Analysis of live mCh and immunofluorescence image data

Images were circumferentially ‘unwrapped’ and fluorescence line profiles were produced as in **Stromal cell transport model** below. For the transduction specificity analysis, a single middle z-plane was chosen per kidney. Cellpose^147^ (v4.0.7) was run on the DAPI channel for each kidney to generate cell segmentations. Cellpose output segmentations were input into FIJI as ROIs and used to create a binarized segmentation mask using the roimanager ‘combine’ function followed by ‘create mask’. Cap mesenchyme and ureteric bud tip niche ROIs were manually segmented using the polygon selection tool and binarized masks of niches were generated in a similar fashion. A custom CellProfiler pipeline was used to convert cell and niche masks to objects, reduce cell object boundaries by 1 pixel to define nuclei, expand nuclei boundaries up to 3 pixels and remove associated nucleus pixels to define cytoplasm, measure nuclear and cytoplasmic channel signal intensity, and measure nucleus shape for each cell. Niche object boundaries were expanded to up to 20 pixels to associate adjacent stroma, and a cell was related to an expanded niche if its centroids fell within the niche’s extent. CellProfiler output files were analyzed in R. Nuclear DAPI intensity, area, and eccentricity were used to filter out poor or erroneous segmentations or cells with rounded nuclei. Manual SIX2 and PDGFRɑ intensity thresholds were used for cell identity classification. Cells with median nuclear intensity above the SIX2 threshold were defined as ‘NP’, cells with median nuclear or cytoplasmic intensity above the PDGFRɑ threshold were defined as ‘stroma’, and all remaining cells were classified as ‘other’. The *fitdist* function (fitdistrplus v1.2-2, 10.18637/jss.v064.i04) using maximum likelihood estimation was used to fit a gamma distribution to the distribution of median nuclear mCherry intensities. The transduction threshold was set as the mCherry intensity at the 50th quantile of the estimated gamma distribution, which was determined using the *qgamma* function (stats v4.4.0) with shape and rate parameters taken from the *fitdist* output. Cells with median nuclear mCherry intensity above the transduction threshold were classified as ‘transduced’ with all remaining cells classified as ‘non-transduced’.

#### 3T3 transduction and western blotting

NIH/3T3 mouse embryonic fibroblasts (gift of Lukasz Bugaj, UPenn) expressing H2B-Venus were transduced using concentrated AAV2-shRNA. Cells were seeded into T25 flasks at 2 x 10^6^ cells per flask and transduced 4 hr post-seeding, allowing for cell attachment to the plate. Cells were transduced for 4 hr at 37°C with AAV2 carrying knockdown constructs at ∼ 7 x 10^11^ GC ml^-1^ in DMEM with 10% FBS and 1% pen/strep (see **Mouse kidney explant adeno-associated virus knockdown**). After 4 hr, media was exchanged with fresh DMEM with 10% FBS and 1% pen/strep. Cells collected at the 72 hr time point had an additional media exchange after 48 hr. At 24, 48, and 72 hr time points, cells were sorted on mCherry intensity using a BD FACSAria Fusion to collect mCherry-positive and negative populations. After 24 hr, two distinct mCherry populations were identifiable with the brighter population (∼50% of cells) collected as mCherry-positive and the dimmer population (∼40% of cells) collected as mCherry-negative cells. Technical issues arose during sorting of the 24 hr shScramble sample preventing collection and use in our validation. Distinct mCherry populations were not visible at 48 and 72 hr time points, so we elected to collect cells within the top and bottom 25% of mCherry intensity as positive and negative fractions, respectively. Sorted fractions were pelleted post sorting, resuspended in DPBS, re-pelleted, and then lysed with RIPA buffer (MedChem Express HY-K1001) containing protease inhibitor (1:100, MedChem Express HY-K0010) on ice for 15 min. Lysates were centrifuged at 14000 rcf for 30 min at 4°C after which the supernatant was collected and maintained at -80°C until use.

Lysates prepared for ROCK1 and ROCK2 evaluation were first denatured by adding equivolume Laemmli buffer (Bio Rad #1610737) with 2-mercaptoethanol (700 mM, Sigma M7154) and then heating samples to 95°C for 20 min. Lysates prepared for MYLK evaluation were diluted to equivolume with Laemmli buffer only without further perturbation. Samples were loaded onto a 4-15% gradient polyacrylamide gel (#4561081, Bio Rad) with 15 µL of sample per lane. Protein size was evaluated against the Precision Plus Protein Kaleidoscope Protein Standard (#1610375, Bio Rad). Gels were run using Tris/Glycine/SDS running buffer (#1610732, Bio Rad) at 120V for 1 hr. Protein was transferred from the polyacrylamide gel onto a nitrocellulose membrane (#170-4270, Bio Rad) using Trans-Blot Turbo transfer buffer (#10026938, Bio Rad). The transfer was performed using a Trans-Blot Turbo Transfer system (Bio Rad 1704150EDU) for 7 min at 13A/25V. Membranes were blocked using casein (#37583, Thermo Scientific) for 2 hr at room temperature. Donkey antibodies were used for primary and secondary labeling and diluted in DPBS, 0.1% tween-20, 1% bovine serum albumin. Primary antibodies were incubated overnight at 4°C and included rabbit anti-ROCK1/2 (1:1000, ab45171, Abcam, RRID:AB_2182005), mouse anti-MYLK (1:1000, SAB4200808, Sigma, RRID:AB_3076644), rabbit anti-RFP (1:1000, ab62341, Abcam, RRID:AB_945213), and mouse anti-ɑ-TUBULIN (1:1000, 3873, Cell Signaling Technologies, RRID:AB_1904178). Secondary antibodies were incubated for 1 hr at 37°C and included goat anti-rabbit IgG IRDYE 680RD (1:10000, 926-68071, LICORbio, RRID:AB_10956166) and goat anti-mouse IgG IRDYE 800CW (1:10000, 926-32210, LICORbio, RRID:AB_621842). Membranes were washed three times for 1 hr at room temperature using the DPBS with 0.1% tween-20 post antibody incubation. Membranes were imaged using a LICORbio Odyssey 9120 instrument.

### Niche composition stochastic models

Niche cell composition dynamics were simulated using stochastic three-state models describing nephron progenitor cells in states A, B, or C. In both models, individual cells undergo state changes and divisions according to a continuous-time Markov process simulated with a Gillespie algorithm^148^ implemented in R. All cells divide symmetrically at rate *k*_4_ while preserving their current state. State transitions between A and B occur with first-order kinetics: in the ‘accumulation’ model, A → B and B → A transitions occur with constant rates *k*_1_ and *k*_2_, respectively.

The B → A transition is supported by lineage tracing^42^. B → C occurs irreversibly at rate *k*_3_, but only within repeating competence windows of duration *t*_trigger_ immediately preceding fixed ‘purge’ times spaced by *t*_gap_, at which all C cells are removed from the system (simulating a PTA leaving the niche/branching/creation of a new PTA site). In the ‘rhythmic’ model, B → A and B → C occur continuously at constant rates *k*_2_ and *k*_3_, respectively. A → B occurs continuously at a rhythmically varying rate *k*_1_(*t*) = *k*_1,mean_[1 + *k*_1,A_.sin(2π*t*/*T* + φ)], where *k*_1,mean_ is the mean value of *k*_1_, *k*_1,A_ is an amplitude expressed as a fraction of the mean value, *T* is rhythm period, and φ is phase. C cells are again purged periodically every *t*_gap_. For both models, initial conditions for simulations were *n*_0_ cells distributed across A, B, and C according to user-defined fractions. We ran 50 replicate simulations per case up to final time *t*_max_, recording cell numbers at regular intervals Δ*t*. From these trajectories, we quantified % niche composition relative to the combined progenitor pool (A + B) for A and B, and relative to the total population (A + B + C) for C. Parameter values estimated from literature sources and assumptions are listed in R code files provided in Mendeley Data (see **Data and code availability**).

### Stromal cell transport model

Stromal transport parameters were fitted from AAV2-mCh-shRNA mouse embryonic kidney explant 3D culture timelapses. Initial and final timepoints at a plane ∼50 µm below the explant surface were unwrapped using a custom macro in FIJI based on bicubic resampling of manually annotated kidney outline polygons and the ‘Straighten…’ function. This created fluorescence ‘curtains’ having image axes tangent (*y*) and orthogonal (*x*) to the explant surface. Shallow-to-deep fluorescence profiles were generated by averaging pixel intensities along *y* using the FIJI function plot profile. Profiles were manually cropped and background-subtracted. Infiltration of stromal cells was modelled using a 1D reaction-diffusion scheme that estimates transport via cell diffusion away from a fixed boundary that produces cells by local proliferation. The evolution of cell fluorescence *C*(*x*,*t*) (AFU) along the corticomedullary axis (x ≥ 0) was described by:

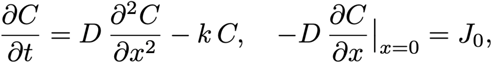

where *D* (µm^2^ h^-1^) is the effective diffusivity associated with cell transport, *J*_0_ (AFU.µm h^-1^) models cell flux due to proliferation at the boundary, and *k* (h^-1^) is a first-order term that captures uniform changes in fluorescence due to e.g. photobleaching of the mCherry reporter. A boundary condition *C*(*x*→∞,*t*) → 0 enforces decay far from the cell fluorescence source. Pairs of spatial profiles *C*(*x*,0) and *C*(*x*,*t*_obs_) were used to estimate *D*, *J*_0_, and *k* by nonlinear least-squares fitting in R. Custom registration, unwrapping, and transport modeling FIJI macros/R code are provided in Mendeley Data (see **Data and code availability**).

### Coding, rendering and graphics

Confocal fluorescence images, sections, and time-coded/other projections were processed in FIJI. Niche branching animation key frames were drawn manually and tweened in Topaz Video AI (Topaz Labs). R code and FIJI macros were created with the assistance of ChatGPT versions 3.5-5.1. All implemented code was closely validated by an expert.

### Statistical analysis

One-way analysis of variance (ANOVA) with correction for multiple comparisons using Tukey’s honestly significant difference test was performed in MATLAB using anova1.m and multcompare.m functions.

