## Supplementary Information for "A mechanical pacemaker sets rhythmic nephron formation in the kidney"

#### Supplementary Notes

**Note S1. Protein-level validation confirms rhythmic naive, priming, and differentiation signatures in the cap mesenchyme.** We sought to validate a subset of the spatial sequencing hits in the nephron progenitor compartment from unsupervised hierarchical clustering and differential expression analysis of human week 20 and mouse E17 kidney slices using immunofluorescence in mouse E17 kidneys.

Wnt/β-catenin is thought to promote nephron progenitor renewal at low activity, and differentiation at high activity[1,2]. ITGA8 is a Wnt/β-catenin target associated with naive, renewing nephron progenitors[3–5]. *ITGA8*/*Itga8* was enriched in post-branching nephron progenitor niches in human/mouse Xenium data, approximately matching enrichment of ITGA8 protein staining at the post- to pre-branching transition (**Fig. S14A**). This supports a period of nephron progenitor renewal late in the branching life-cycle.

Autocrine BMP7 primes nephron progenitors, sensitizing them to subsequent commitment induced by Wnt/β-catenin[6]. BMP7 signals via pSMAD1/5 to activate the target gene *Id1* (ref. [6,7]), which was enriched in pre-branching niches in human and mouse Xenium data (**Fig. S14B**). pSMAD1/5 protein staining was similarly enriched in pre-branching niches, supporting priming of nephron progenitors early in the branching life-cycle.

We next turned to the Wnt/β-catenin targets CCND1 and PAX8 that mark nephron progenitor differentiation[3,4], as well as MKI67, which marks proliferative cells in all active phases of the cell cycle, especially S-phase[8] (**Fig. S14C**). Each of these was generally enriched in pre-branching nephron progenitor niches in human and mouse Xenium data, approximately matching enrichment at the pre-branching stage or post- to pre-branching transition at the protein level. An exception was that the human *CCND1* Xenium data did not match the trend of the mouse Xenium and IF data, perhaps due to low expression at the tissue depth range analyzed. These data indicate Wnt-driven nephron progenitor differentiation and proliferation during early- to mid-branching, which is thought to drive their condensation via mesenchymal-to-epithelial transition[9], and later to establish proximodistal polarity in the early nephron[10].

Together the validation data reinforce the notion that nephron progenitors alternatingly favor differentiation/renewal early/late in the branching cycle, respectively.

**Note S2. Interpretation of branching life cycle correlation with Hippo/Yap, Wnt/β-catenin, and retinoic acid signaling in nephron progenitors.** In the mouse nephron progenitor compartment, upregulated genes in Wnt and retinoic acid receptor signaling pathway sets were significantly over-represented in the pre-branching phase, while those in a Hippo signaling pathway set were significantly over-represented in post-branching and early pre-branching phases (**Fig. S3E**). Specific genes contributing to these lists included those consistent with increased Wnt/β-catenin signaling (*Lrp6*, *Wnt4*, *Myc*; as well as nephron progenitor differentiation-specific target genes[3,4,11] *Ccnd1*, *Aldh1a2*, *Lhfpl2* from differential expression, DE), increased retinoic acid signaling[12] (*Aldh1a2*, *Rxra*), and increased Yap signaling (*Tead3*, *Myc*, *Ccnd2*; as well as nephron progenitor-specific target genes[13] *Cited1*, *Traf1*, and *Capn6* from DE) in these phases. We validated that YAP nuclear localization peaked during post-branching in E17 mouse kidney nephron progenitors by whole-mount immunofluorescence, which is compatible with the timing of transcript-level changes noted above (**Fig. S14A**). Wnt/β-catenin target gene timing is similarly validated in **Fig. S14C** (**Note S1**). The relative timing of these pathways is less clear in human week 20 nephron progenitors, perhaps due to the two distinct and potentially temporally staggered routes of nephron progenitor differentiation (**Fig. 4, Fig. S21**) that are averaged in the pseudobulk analyses here. Even so, upregulated genes in a canonical Wnt signaling pathway activity set were significantly over-represented in pre-branching, extending into mid-branching (specific genes included *DVL1/2*, *CTNNB1* consistent with increased activity; as well as the nephron progenitor differentiation-specific target gene[3,14] *PAX8* from DE). Upregulated genes in Hippo signaling pathway and response to retinoic acid gene sets were both significantly over-represented in mid-branching (**Fig. 3D**). Specific Hippo signaling pathway genes in mid-branching included those consistent with increased activity (*WWTR1, TGFB3, GLI2)*. However, specific response to retinoic acid genes in mid-branching included *FGFR2* consistent with decreased activity[12]; and *RXRA* from DE was differentially expressed in pre-branching rather than mid-branching, consistent with increased response to retinoic acid in pre-branching. Together the data indicate that Wnt/β-catenin and retinoic acid signaling activities are likely higher in nephron progenitors in pre-branching niches, while YAP activity is likely higher in those in mid-/post-branching niches.

**Note S3. Rhythmically regulated pathways in the human niche are sufficient to modify interpretation of Wnt differentiation signaling in human stem-cell derived nephron progenitors.** We turned to human iPSC-derived nephron progenitor organoids[15–17] to validate the effects of rhythmic pathways in nephron progenitors, while excluding non-autonomous effects from the UB and stroma. A 'pulse' of Wnt/β-catenin activation via the GSK3β inhibitor CHIR 99021 is commonly used here to increase differentiation efficiency of SIX2+ nephron progenitors[15,18]. We reasoned that combining transient Wnt activation with simultaneous or staggered perturbation of candidate signaling pathways from our spatial sequencing screen would cause differences in nephron progenitor renewal vs. differentiation if rhythmic signaling was relevant to maintaining this balance. We focused on YAP and retinoic acid signaling due to the rhythmic Hippo and retinoic acid signaling signatures we uncovered during gene set over-representation analysis (**Fig. 3D, Fig. S3E**). Despite some species-specific differences, iPSC-derived nephron progenitors roughly transit through the same states when differentiated (transcriptionally and in protein marker expression) as nephron progenitors in the human and mouse niches[15,18–21]. Increased YAP signaling *in vivo* and *in vitro* appears to drive nephron progenitor renewal at the expense of differentiation, and vice versa[13,22–27]. TRULI has recently been found to recruit YAP to the nucleus of iPSC-derived nephron progenitors and extend their long-term renewal in 2D culture, while retaining their nephron differentiation potential[24]. We verified that TRULI increases YAP nuclear recruitment and target gene expression (*CTGF*, *CYR61*) in day 10 human iPSC-derived nephron progenitors by immunofluorescence and qPCR respectively (**Fig. S15**). We then found that TRULI increased the representation of SIX2+ nephron progenitors in organoids at day 12 only when it temporally overlapped the 2 hr CHIR pulse at day 10, but not when it chased the CHIR pulse for 2 hr (**Fig. S16A-D**). However, YAP activation suppressed differentiation of cells to the JAG1+ renal vesicle state at day 12 in either case (**Fig. S16D**), which also translated to lower nephron formation at day 25 (**Fig. S16E,F**). Changes in nephron composition among the conditions were minimal (**Fig. S17**). These data indicate that the basal YAP state in nephron progenitor organoids permits Wnt-induced differentiation, while YAP activation converts the same Wnt signal into a renewal signal when the two temporally overlap. This interaction may relate to the paradoxical requirement of Wnt/β-catenin for both nephron progenitor renewal and differentiation[28], namely that a rhythmic YAP signaling context could enable the niche to alternately favor each state given a consistent Wnt/β-catenin input *in vivo*.

While little is known about the role of retinoic acid after specification of nephron progenitors, microarray and qPCR analysis of RA-treated mouse embryonic kidney explants saw up-regulation in early nephron differentiation markers including *Pax8*, *Lhx1* and *Wnt4* relative to untreated controls[29]. Retinoic acid synthesis enzymes are also expressed in early rat kidney connecting segments and renal vesicles[30]. Retinoic acid promotes proximal segment fate at the expense of distal segment in the zebrafish pronephros, but a similar role in the segmentation of mammalian nephrons has not yet been established[31,32]. Huang *et al.* found that the retinoic acid analogue TTNPB[33,34] is necessary for polarized renal vesicle formation in hPSC-derived UB/NP mosaic organoids[35]. We verified that TTNPB increases retinoic acid target gene expression (*RARB*, *CRABP2, CYP26A1*) in iPSC-derived nephron progenitors by qPCR (**Fig. S15C**). Staggered or overlapping TTNPB exposure around the 2 hr CHIR pulse at day 10 reduced the SIX2+ cell fraction at day 12 and restricted SIX2 expression to spatially discrete colonies within organoids (**Fig. S16G-I**). Neither treatment timing affected the fraction differentiated to the JAG1+ state at day 12 (**Fig. S16I**), reflecting either off-target differentiation of SIX2+ cells or transit through to later differentiation states. Nephron formation by day 25 was robust for both TTNPB treatment timings (**Fig. S16J,K**). GATA3+ connecting segments appeared and nephrons were notably more spatially discrete, with frequent formation of polarized tubules with correct anatomical connectivity of connecting segment-distal tubule-proximal tubule-podocyte cells (**Fig. S16J,K)**. LTL+ proximal tubule segments were also observed more frequently, suggesting that retinoic acid indeed has a similar proximalizing effect here as in the zebrafish. These data indicate that retinoic acid signaling may increase progenitor exit from the SIX2+ state, restrict the spatial scale of Wnt-induced differentiation and promote an appropriate bias of fates during subsequent nephron segmentation.

**Note S4. Cell cycle rate correlates with progressive recruitment of nephron progenitors in UB-adjacent and distant streams.** Lindstrom *et al.* proposed a progressive recruitment model of nephron progenitor induction and contribution to the early nephron, where early-induced cells contribute to distal nephron cells, and late-induced cells contribute to proximal nephron/glomerular cells[36]. We saw similar evidence in our spatial sequencing data, with an added observation of higher cell cycle rate in the early-induced distal stream. First, cells in our 'primed' Xenium cluster are geometrically closer to UB than 'committing' cluster cells, supporting the existence of distinct UB-adjacent vs. distant induced nephron progenitor streams as proposed by Fausto *et al.*[37] (**Fig. S21A**). The primed cluster had almost all cells in S/G2M, while the committing cluster had a larger proportion of cells in G1 (**Fig. S21B**). Both primed and committing cells were geometrically nearby early nephron cells, but a higher proportion of primed cluster cells were nearby connecting segment/distal cells, while a higher proportion of committing cluster cells were nearby proximal/glomerular cells (**Fig. S21C**). Committing cluster cells were found at the outer border of renal corpuscles in deeper areas of a representative human week 20 Xenium slice, suggesting that they join the proximal pole of the forming nephron later than other cells, matching Lindstrom's progressive recruitment model (**Fig. S21D**). These data imply that the close-to-UB, fast-cycling, early-induced primed cell stream contribute to the distal pole of new nephrons; while the far-from-UB, slow-cycling, late-induced committing cell stream contribute to the proximal pole of new nephrons (**Fig. S21E**). The faster cycling of distal early nephron cells was also noted by Georgas *et al.*[11].

**Note S5. Rhythmic, stromally expressed SFRP1 preserves nephron progenitor naivety in mouse embryonic kidney explant cultures.** We treated mouse embryonic kidneys with the SFRP1 inhibitor WAY-316606 (SFRP1i) for 72 hr in 3D explant culture (**Fig. S23**). SFRP1i led to a reduction in cap mesenchyme and ureteric bud cross-sectional areas as well as cap mesenchyme depth. Staining for the naive NPC marker CITED1 showed that its expression shifted from a uniform distribution in the cap mesenchyme to a radial gradient upon SFRP1i treatment, with expression retained only adjacent to the ureteric bud tip. This resulted in an increased interquartile range (IQR) in CITED1 fluorescence intensity within individual niches. These data suggest that SFRP1 preserves nephron progenitor naivety.

### Supplementary Figures

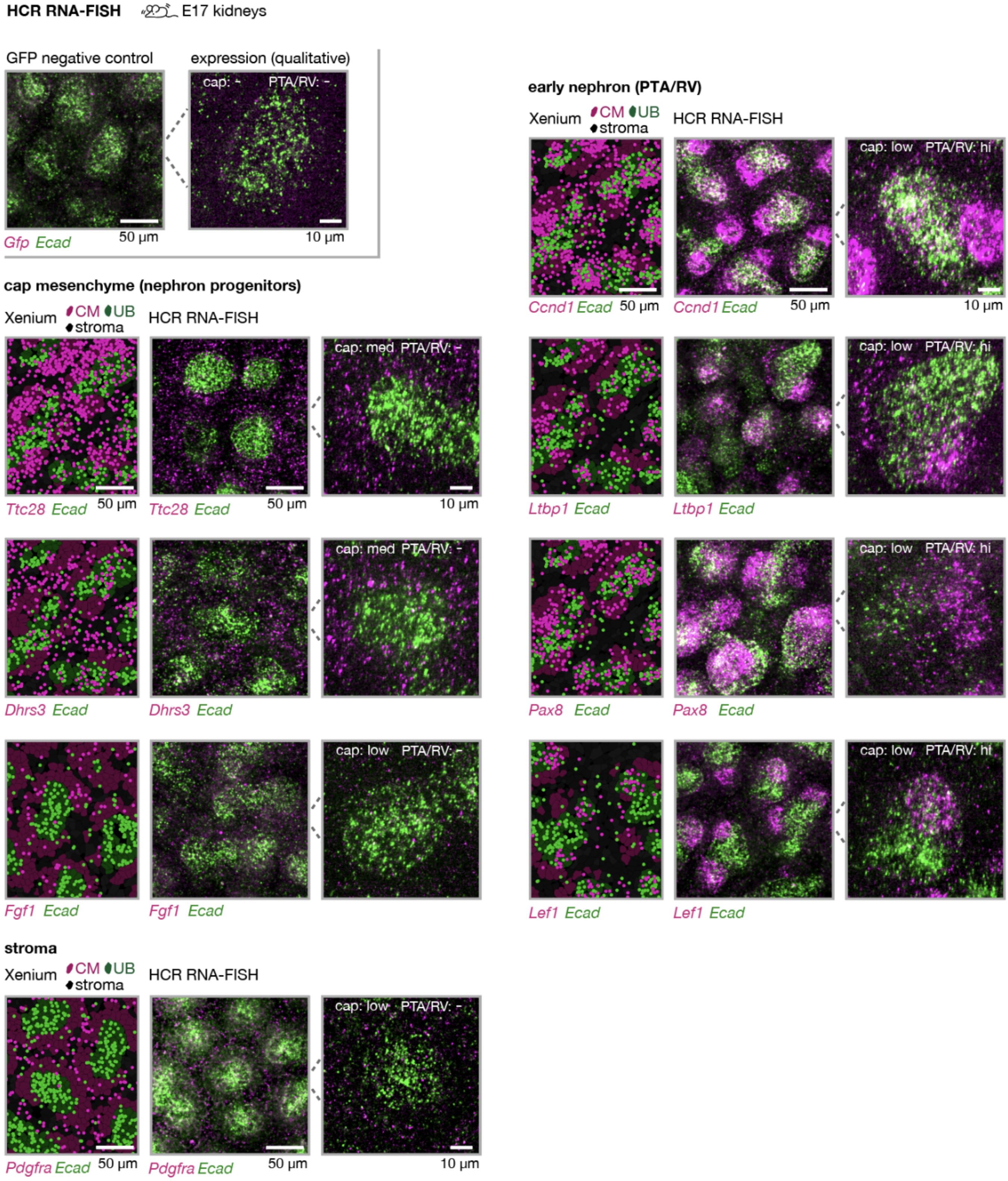


**Fig. S1: Validation of Xenium transcript localization by HCR RNA-FISH.** Representative Xenium and 20x/40x whole-mount confocal fluorescence micrographs of E17 kidneys at approximate mid-planes of ureteric bud epithelial tips and surrounding nephrogenic niches. Qualitative expression levels in cap mesenchyme (‘cap’) and pre-tubular aggregates/renal vesicles (‘PTA/RV’) are noted, relative to *Gfp* negative control, listed as low, medium (med), high (hi), or no expression observed using the chosen probe set (-).

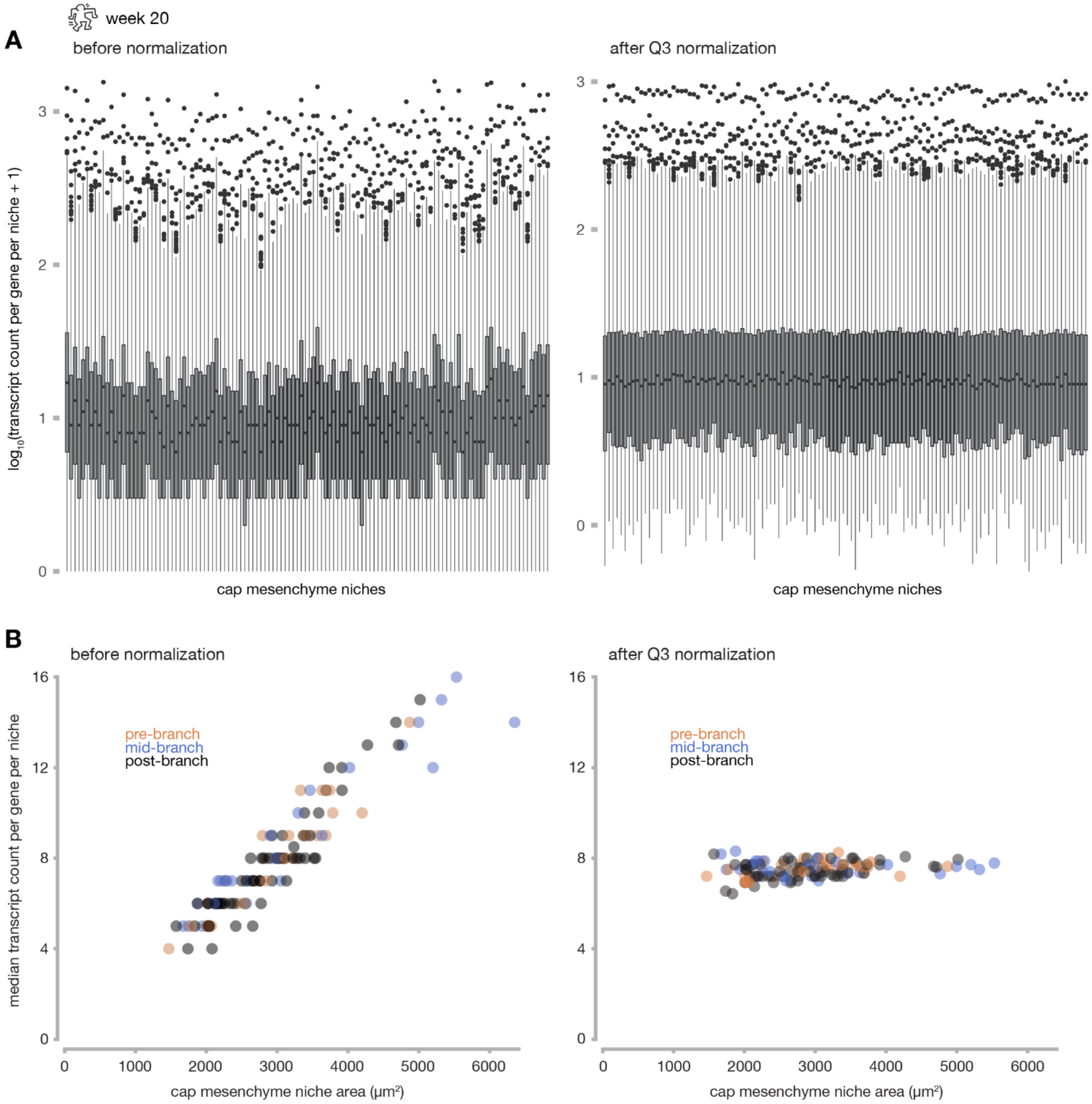


**Fig. S2: Normalization accounts for variation in niche size.** Human week 20 kidney slice, $n$ = 120 niches. Data are typical of results in all human and mouse datasets. (**A**) *Left*, Box plots of log-transformed counts for pseudobulked transcripts in the cap mesenchyme compartment (i.e. summed over nephron progenitor cells within each niche). *Right*, Similar box plots after Q3 normalization of data. (**B**) Median expression vs. niche area (sum of areas of nephron progenitor cells) before (*left*) and after (*right*) normalization.

195
196

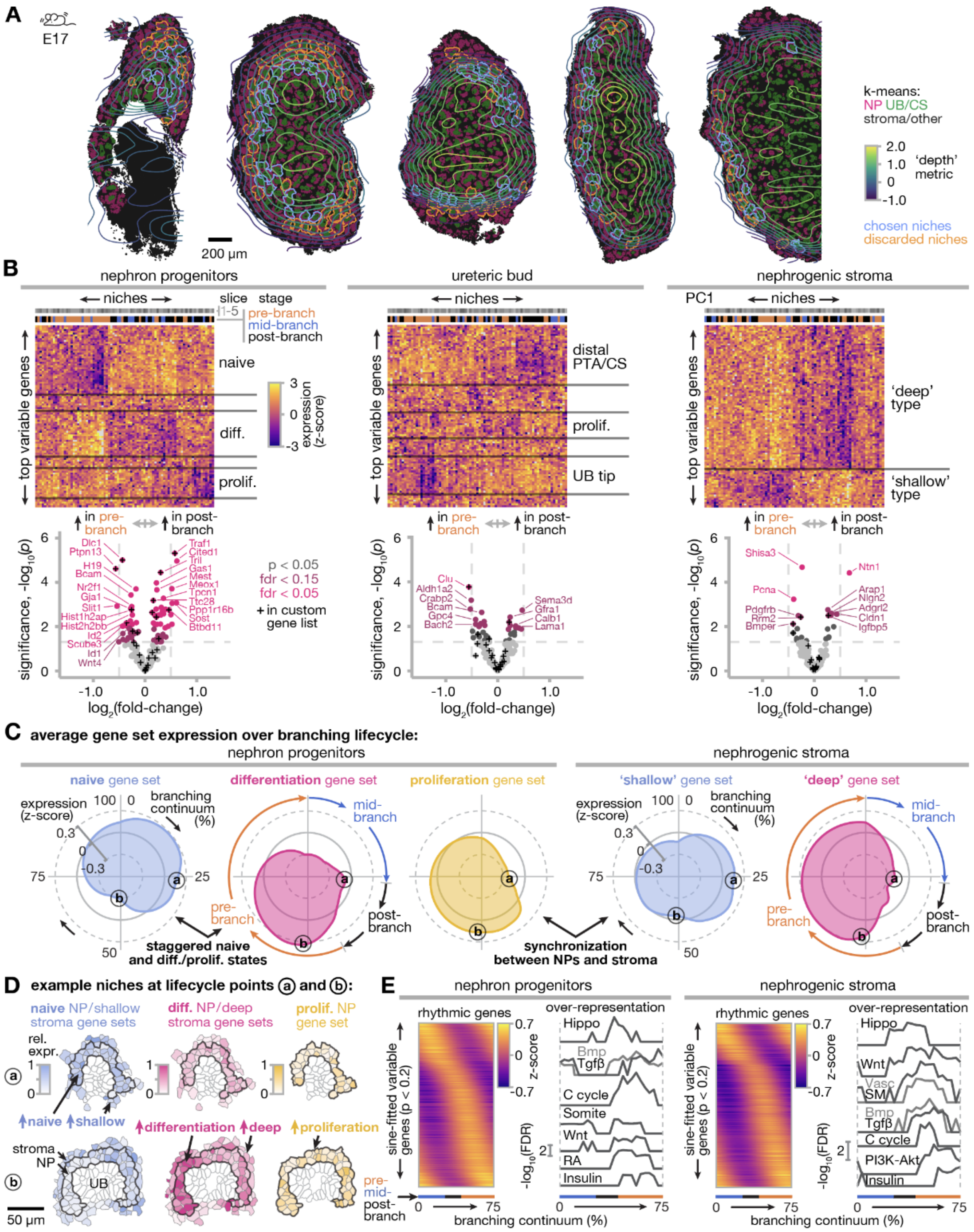


**Fig. S3: Nephron progenitor and stromal compartments of the mouse nephron-forming niche vary transcriptionally in renewal and differentiation pathways depending on ureteric bud branching life-cycle stage.** (**A**) Contours of the depth metric overlaid on k-means Xenium view, highlighting 'chosen' niches sharing a similar anatomical position in depth. NP: nephron progenitor, UB/CS: ureteric bud/connecting segment. (**B**) Unsupervised hierarchical clustering heatmaps (top 100 variable genes, *top*) and differential expression plots (top 200 variable genes, *bottom*) for pseudobulked transcript expression across niches at different branching lifecycle stages ($n$ = 81 niches across 5 slices from 5 different kidneys, see **Fig. S11** for full analysis). Blocks of genes are qualitatively annotated based on preponderance of genes previously associated with the indicated cell states. As part of Xenium analysis, we pre-selected 100 genes (of the 5106 total) known to be relevant to early cell decision making, which are indicated with crosses. (**C**) Radial plots of z-scores for gene sets differentially expressed in (B) and qualitatively associated with the indicated cell states for nephron progenitor and stroma cell compartments. Radial distance from the center is expression level, angular distance clockwise from the 12 o'clock position indicates branching continuum. (a) and (b) labels correspond to rows in (D). (**D**) Heatmaps of gene set expression for representative niches at single-cell resolution in renewal (a) and differentiation (b) phases. (**E**) Heatmap of expression vs. branching continuum for rhythmic genes ordered by peak position fitting a sine wave model at $p < 0.2$ and line plots of GO/KEGG/reactome analyses performed using a sliding window over the same genes (Hippo, hippo signaling pathway; Bmp, BMP signaling pathway; Tgfβ, TGF−beta signaling pathway; C cycle, Cell cycle; Somite, somite development; Wnt, Wnt signaling pathway; RA, retinoic acid receptor signaling pathway; Insulin, insulin receptor signaling pathway; Vasc, kidney vascular development; SM, regulation of smooth muscle cell proliferation; PI3K-Akt, PI3K-Akt signaling).

197

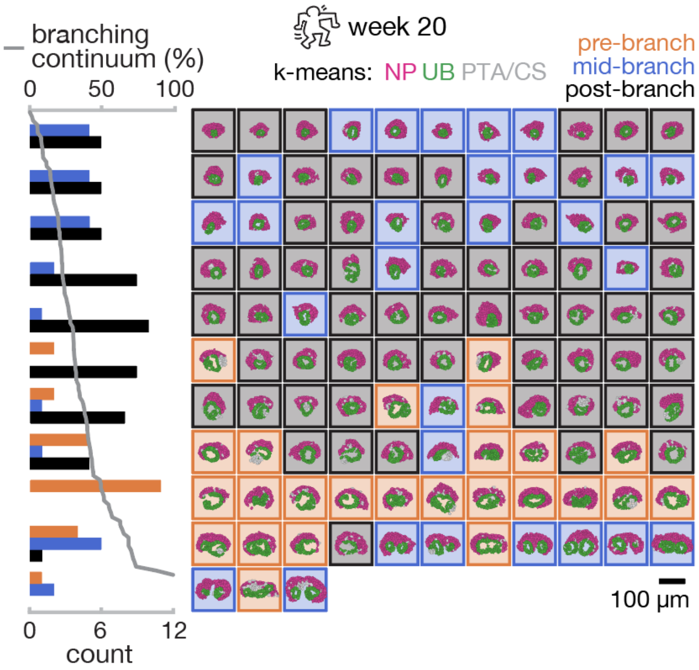


**Fig. S4: Discrete and continuous branching stage annotations for human week 20 kidney.** Histogram of discrete branching stage counts per row of a matrix of niches arranged left-to-right and top-to-bottom by increasing branching continuum. Niche outlines are colored by discrete branching stage. All nephron progenitor and ureteric bud cells associated with each niche are shown, colored by Xenium k-means cluster. NP: nephron progenitor, UB: ureteric bud, PTA/CS: pretubular aggregate/connecting segment.

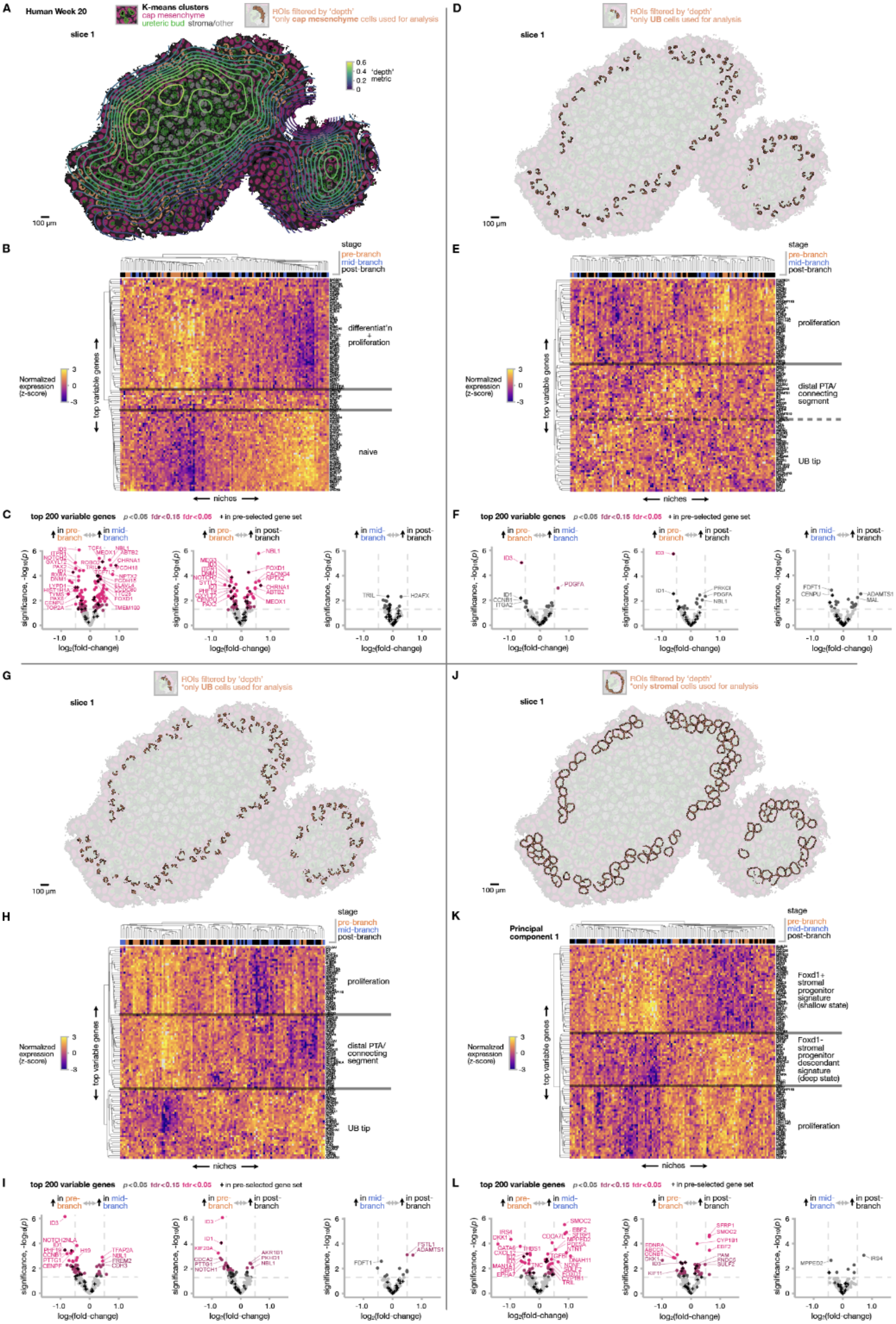


**Fig. S5: Human week 20 nephrogenic niche selection and Xenium rhythmic gene expression analysis for niche cell compartments.** (**A**) Contours of the depth metric overlaid on k-means Xenium view, highlighting cap mesenchyme cells (nephron progenitors) in niches filtered for similar anatomical position in depth. (**B**) Unsupervised hierarchical clustering heatmap (top 100 variable genes) for pseudobulked transcript expression across niches at different branching lifecycle stages (*n* = 120 niches). Blocks of genes are qualitatively annotated based on preponderance of genes previously associated with the indicated cell states. (**C**) Differential expression plots (top 200 variable genes) for different niche branching lifecycle stage comparisons. As part of Xenium analysis, we pre-selected 91 genes (of the 5092 total) known to be relevant to nephrogenic niche development, which are indicated with crosses. (**D-F**) The same figure elements for ureteric bud cells neighboring the cap mesenchyme. (**G-I**) The same figure elements for ureteric bud cells/pretubular aggregate/connecting segment cells neighboring the cap mesenchyme. (**J-L**) The same figure elements for stromal cells lining the niche.

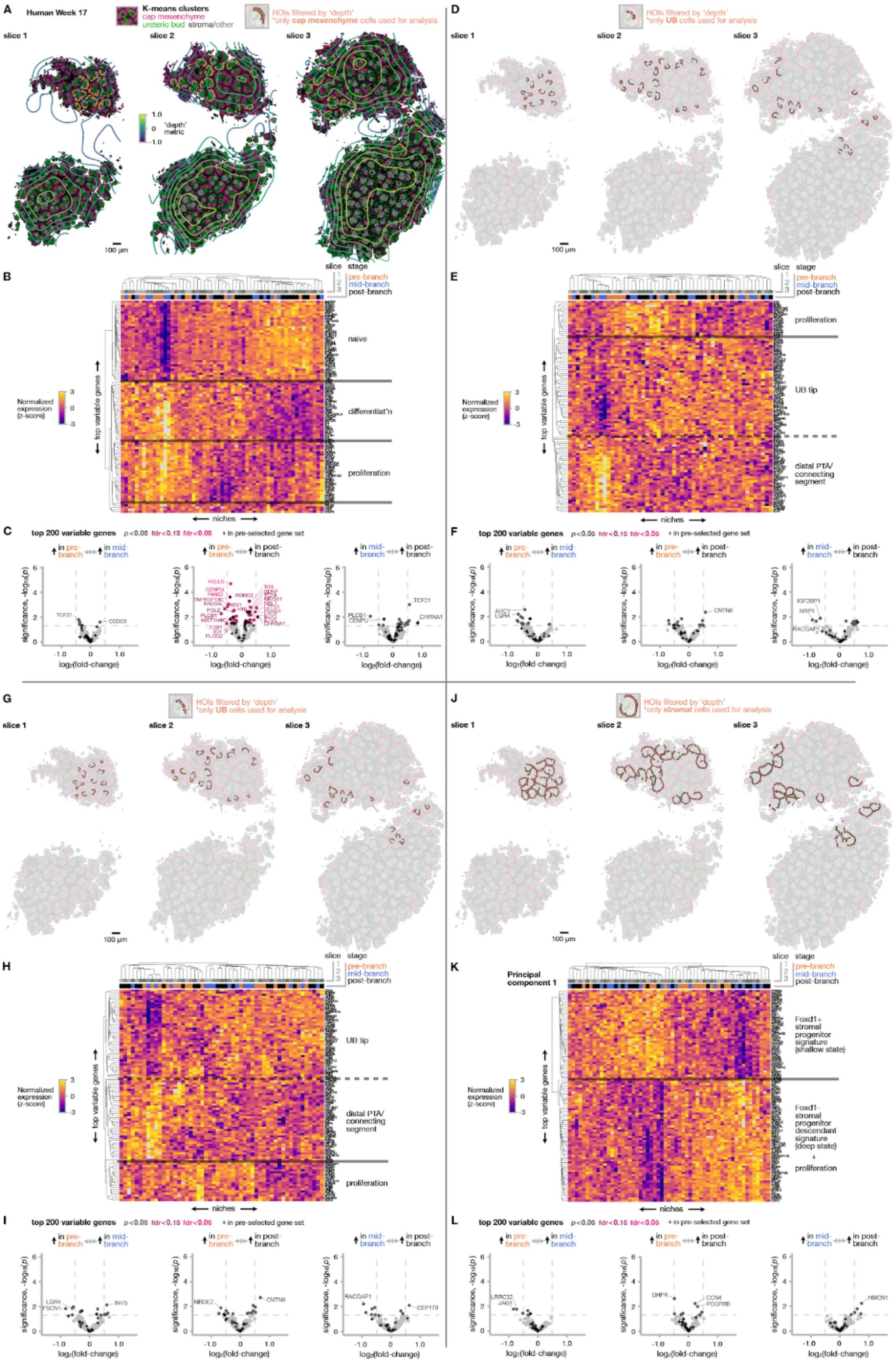


**Fig. S6: Human week 17 nephrogenic niche selection and Xenium rhythmic gene expression analysis for niche cell compartments.** (**A**) Contours of the depth metric overlaid on k-means Xenium view, highlighting cap mesenchyme cells (nephron progenitors) in niches filtered for similar anatomical position in depth. (**B**) Unsupervised hierarchical clustering heatmap (top 100 variable genes) for pseudobulked transcript expression across niches at different branching lifecycle stages ($n$ = 57 niches across 3 slices of the same kidney). Blocks of genes are qualitatively annotated based on preponderance of genes previously associated with the indicated cell states. (**C**) Differential expression plots (top 200 variable genes) for different niche branching lifecycle stage comparisons. As part of Xenium analysis, we pre-selected 91 genes (of the 5092 total) known to be relevant to nephrogenic niche development, which are indicated with crosses. (**D-F**) The same figure elements for ureteric bud cells neighboring the cap mesenchyme. (**G-I**) The same figure elements for ureteric bud cells/pretubular aggregate/connecting segment cells neighboring the cap mesenchyme. (**J-L**) The same figure elements for stromal cells lining the niche.

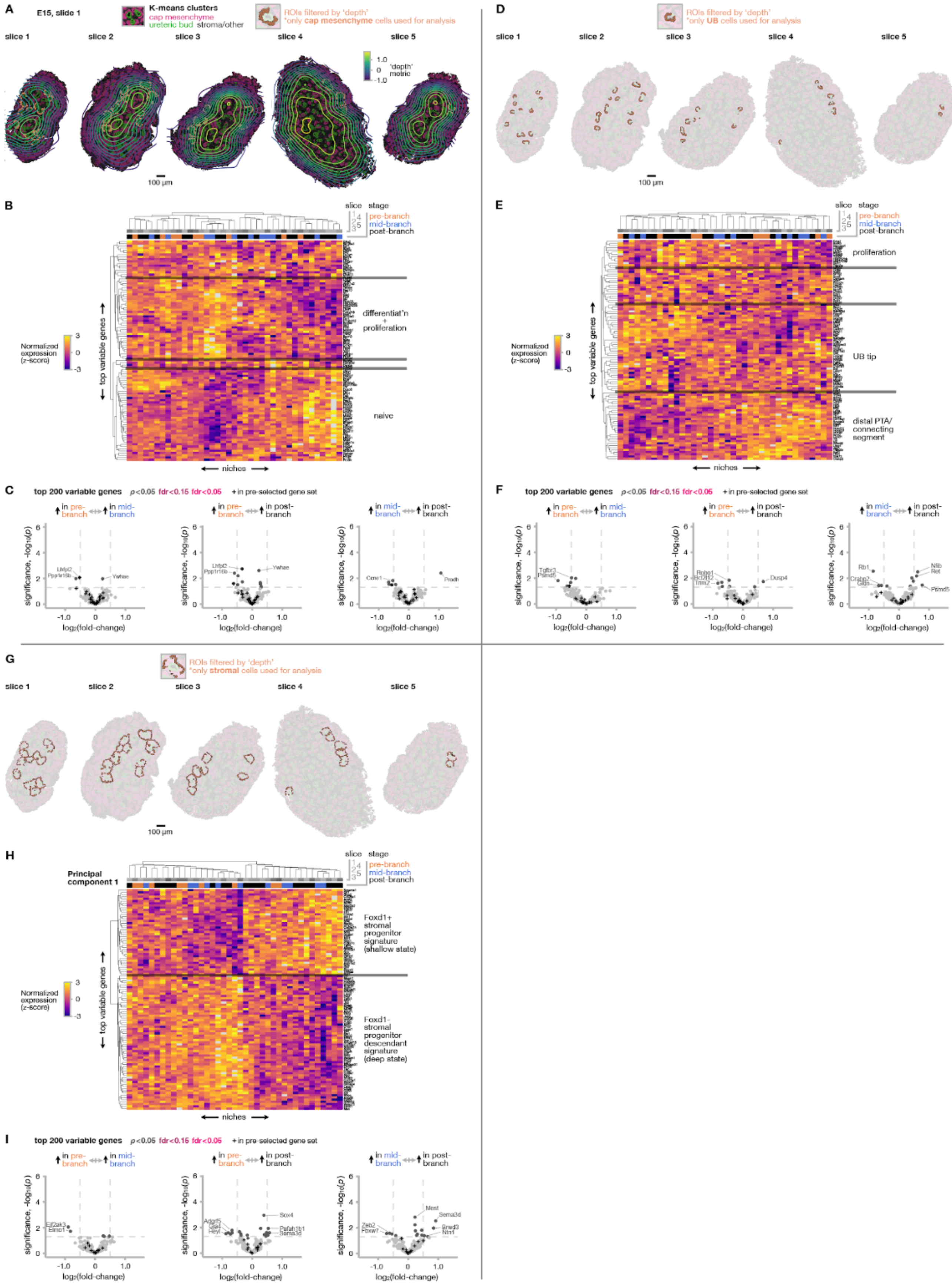


**Fig. S7: Mouse E15 (slide 1) nephrogenic niche selection and Xenium rhythmic gene expression analysis for niche cell compartments.** (**A**) Contours of the depth metric overlaid on k-means Xenium view, highlighting cap mesenchyme cells (nephron progenitors) in niches filtered for similar anatomical position in depth. (**B**) Unsupervised hierarchical clustering heatmap (top 100 variable genes) for pseudobulked transcript expression across niches at different branching lifecycle stages (*n* = 39 niches across 5 slices from 4 different kidneys). Blocks of genes are qualitatively annotated based on preponderance of genes previously associated with the indicated cell states. (**C**) Differential expression plots (top 200 variable genes) for different niche branching lifecycle stage comparisons. As part of Xenium analysis, we pre-selected 100 genes (of the 5106 total) known to be relevant to nephrogenic niche development, which are indicated with crosses. (**D-F**) The same figure elements for ureteric bud cells neighboring the cap mesenchyme. (**G-I**) The same figure elements for stromal cells lining the niche.

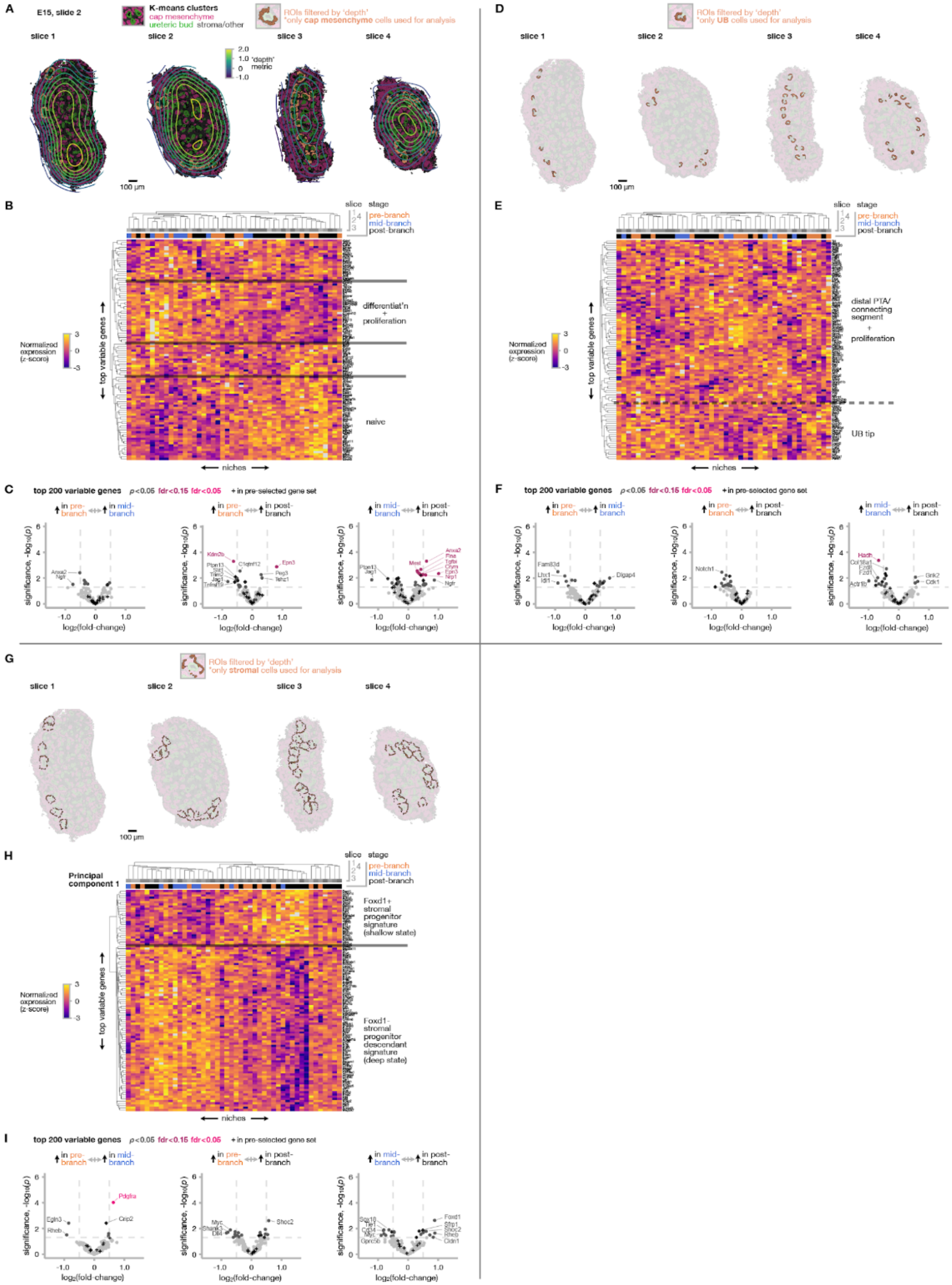


**Fig. S8: Mouse E15 (slide 2) nephrogenic niche selection and Xenium rhythmic gene expression analysis for niche cell compartments.** (**A**) Contours of the depth metric overlaid on k-means Xenium view, highlighting cap mesenchyme cells (nephron progenitors) in niches filtered for similar anatomical position in depth. (**B**) Unsupervised hierarchical clustering heatmap (top 100 variable genes) for pseudobulked transcript expression across niches at different branching lifecycle stages ($n$ = 46 niches across 4 slices from 2 different kidneys). Blocks of genes are qualitatively annotated based on preponderance of genes previously associated with the indicated cell states. (**C**) Differential expression plots (top 200 variable genes) for different niche branching lifecycle stage comparisons. As part of Xenium analysis, we pre-selected 100 genes (of the 5106 total) known to be relevant to nephrogenic niche development, which are indicated with crosses. (**D-F**) The same figure elements for ureteric bud cells neighboring the cap mesenchyme. (**G-I**) The same figure elements for stromal cells lining the niche.

206

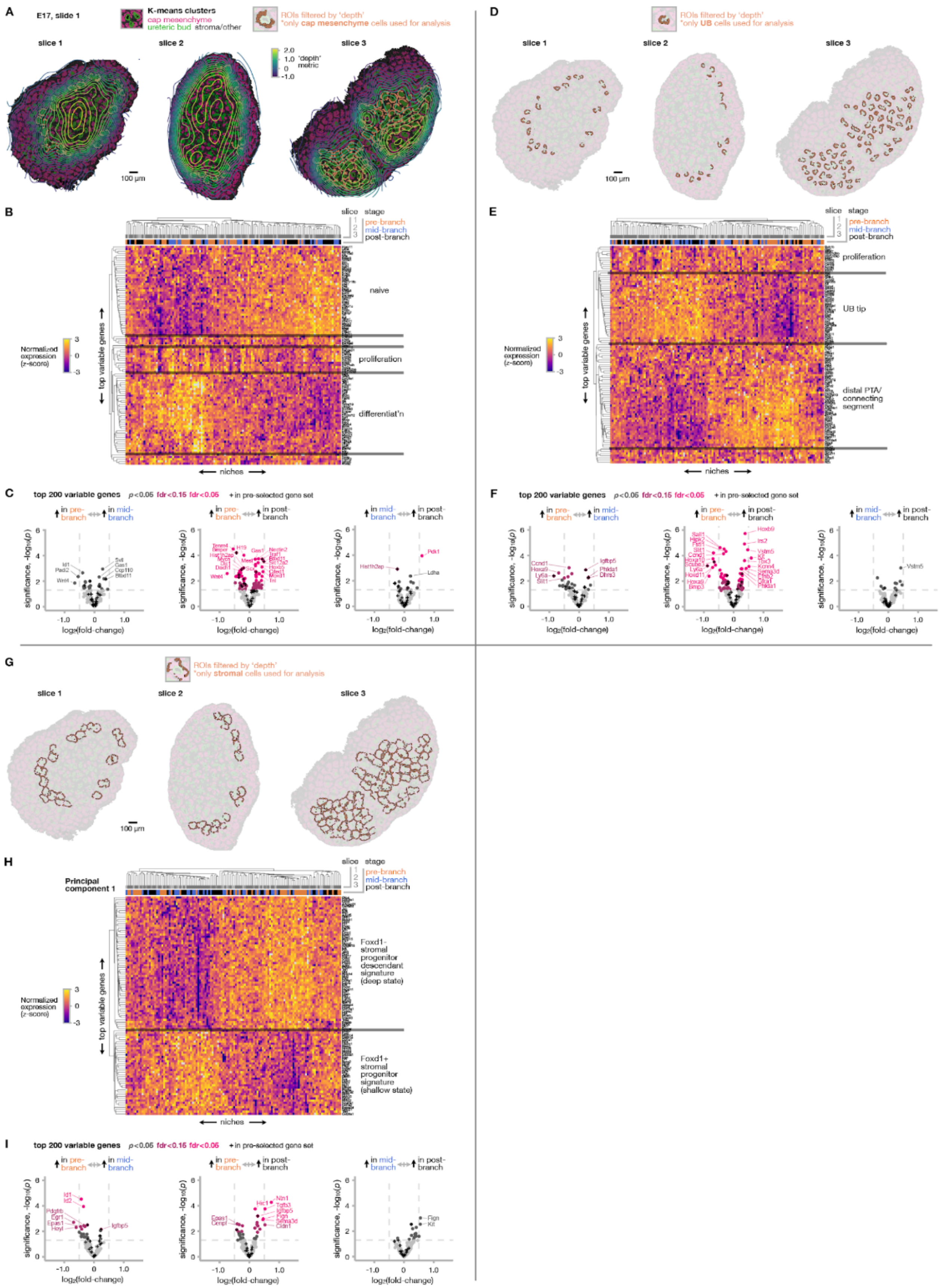


**Fig. S9: Mouse E17 (slide 1) nephrogenic niche selection and Xenium rhythmic gene expression analysis for niche cell compartments.** (**A**) Contours of the depth metric overlaid on k-means Xenium view, highlighting cap mesenchyme cells (nephron progenitors) in niches filtered for similar anatomical position in depth. (**B**) Unsupervised hierarchical clustering heatmap (top 100 variable genes) for pseudobulked transcript expression across niches at different branching lifecycle stages ($n$ = 120 niches across 3 slices from 3 different kidneys). Blocks of genes are qualitatively annotated based on preponderance of genes previously associated with the indicated cell states. (**C**) Differential expression plots (top 200 variable genes) for different niche branching lifecycle stage comparisons. As part of Xenium analysis, we pre-selected 100 genes (of the 5106 total) known to be relevant to nephrogenic niche development, which are indicated with crosses. (**D-F**) The same figure elements for ureteric bud cells neighboring the cap mesenchyme. (**G-I**) The same figure elements for stromal cells lining the niche.

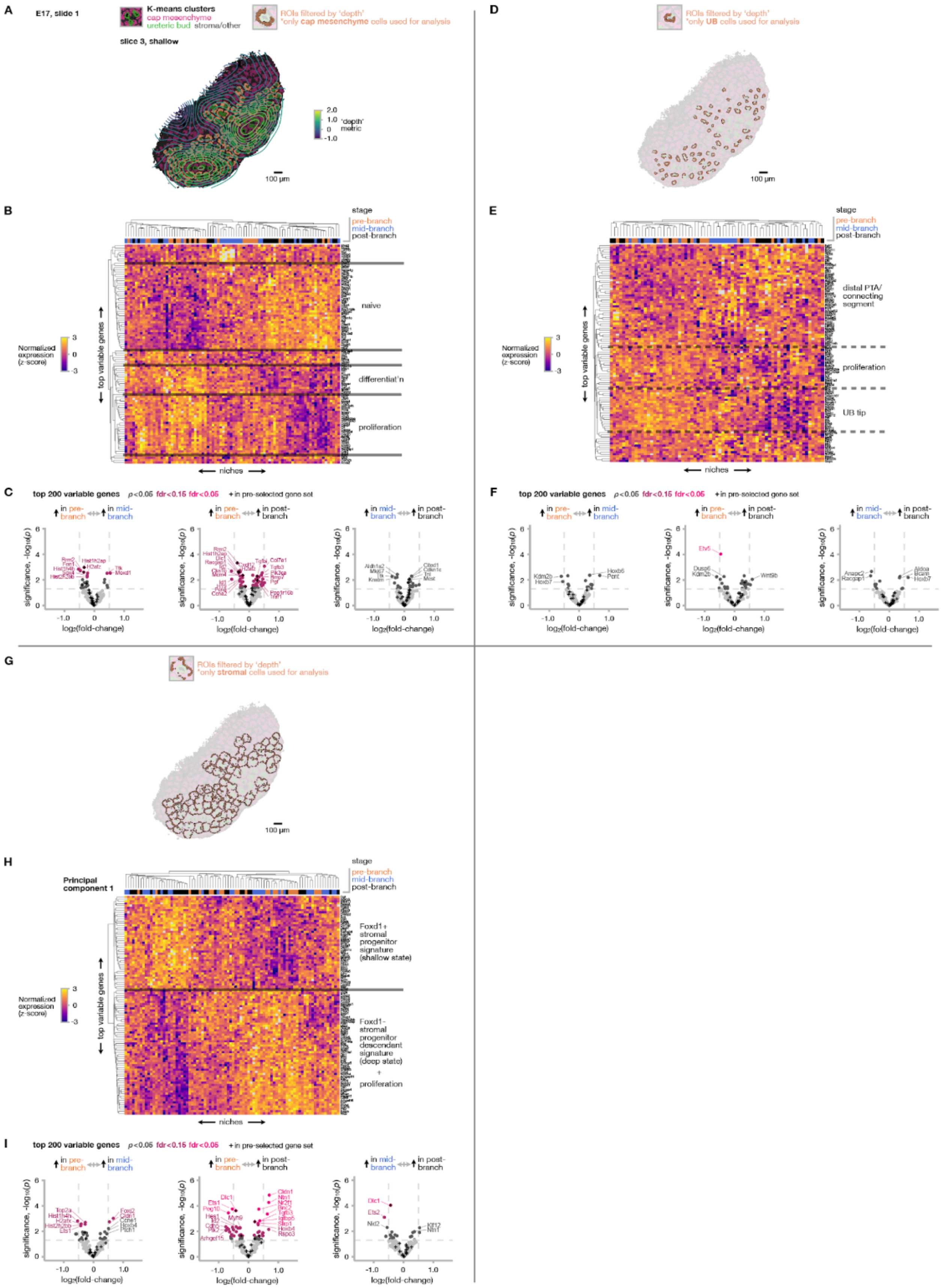

**Fig. S10: Mouse E17 (slide 1, shallower slice from the same kidney associated with Fig. S9 slice 3) nephrogenic niche selection and Xenium rhythmic gene expression analysis for niche cell compartments.** (**A**) Contours of the depth metric overlaid on k-means Xenium view, highlighting cap mesenchyme cells (nephron progenitors) in niches filtered for similar anatomical position in depth. (**B**) Unsupervised hierarchical clustering heatmap (top 100 variable genes) for pseudobulked transcript expression across niches at different branching lifecycle stages ($n$ = 84 niches). Blocks of genes are qualitatively annotated based on preponderance of genes previously associated with the indicated cell states. (**C**) Differential expression plots (top 200 variable genes) for different niche branching lifecycle stage comparisons. As part of Xenium analysis, we pre-selected 100 genes (of the 5106 total) known to be relevant to nephrogenic niche development, which are indicated with crosses. (**D-F**) The same figure elements for ureteric bud cells neighboring the cap mesenchyme. (**G-I**) The same figure elements for stromal cells lining the niche.

208

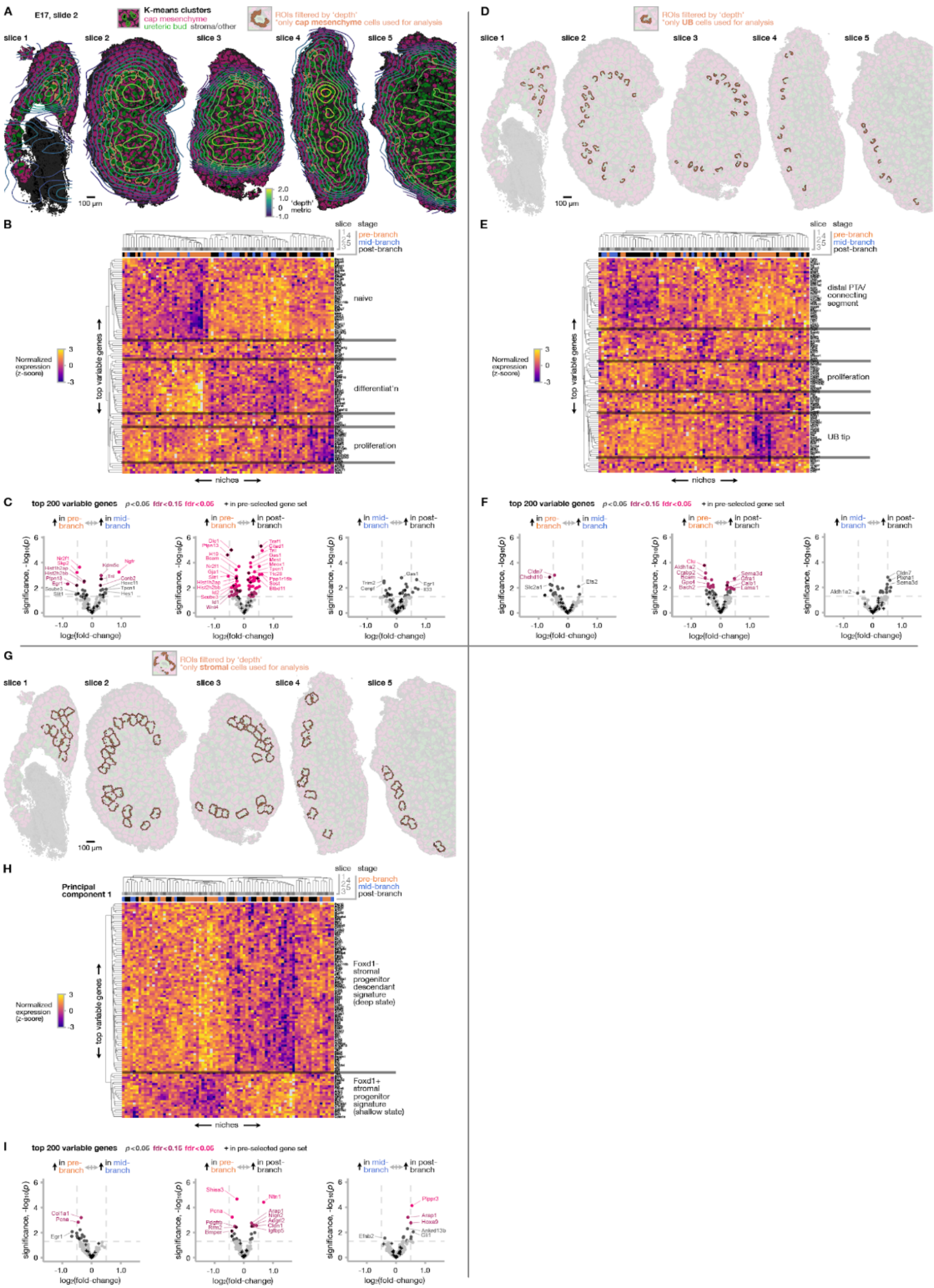


**Fig. S11: Mouse E17 (slide 2) nephrogenic niche selection and Xenium rhythmic gene expression analysis for niche cell compartments.** (**A**) Contours of the depth metric overlaid on k-means Xenium view, highlighting cap mesenchyme cells (nephron progenitors) in niches filtered for similar anatomical position in depth. (**B**) Unsupervised hierarchical clustering heatmap (top 100 variable genes) for pseudobulked transcript expression across niches at different branching lifecycle stages (*n* = 81 niches across 5 slices from 5 different kidneys). Blocks of genes are qualitatively annotated based on preponderance of genes previously associated with the indicated cell states. (**C**) Differential expression plots (top 200 variable genes) for different niche branching lifecycle stage comparisons. As part of Xenium analysis, we pre-selected 100 genes (of the 5106 total) known to be relevant to nephrogenic niche development, which are indicated with crosses. (**D-F**) The same figure elements for ureteric bud cells neighboring the cap mesenchyme. (**G-I**) The same figure elements for stromal cells lining the niche.

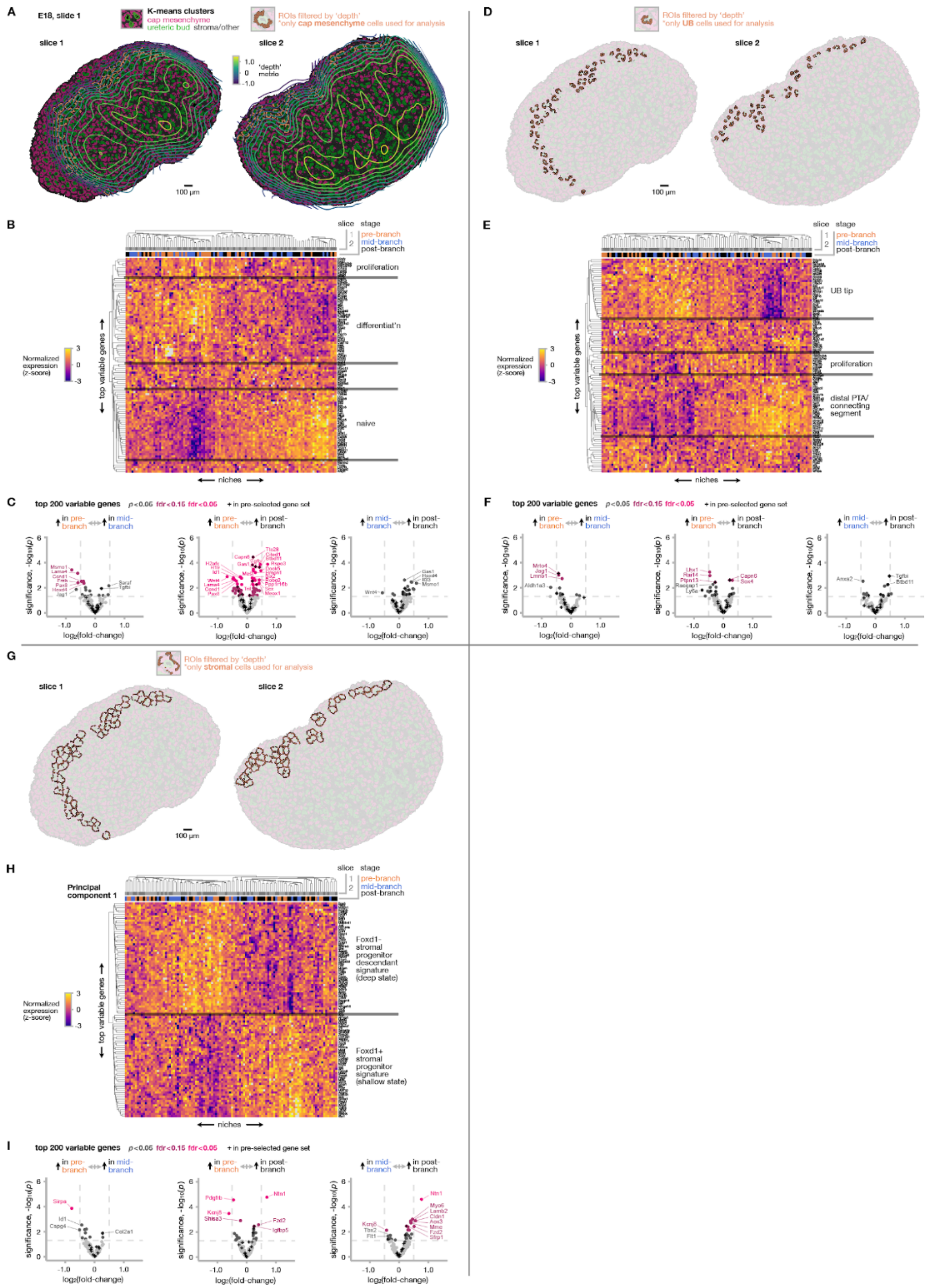

**Fig. S12: Mouse E18 (slide 1) nephrogenic niche selection and Xenium rhythmic gene expression analysis for niche cell compartments.** (**A**) Contours of the depth metric overlaid on k-means Xenium view, highlighting cap mesenchyme cells (nephron progenitors) in niches filtered for similar anatomical position in depth. (**B**) Unsupervised hierarchical clustering heatmap (top 100 variable genes) for pseudobulked transcript expression across niches at different branching lifecycle stages (*n* = 72 niches across 2 slices from 2 different kidneys). Blocks of genes are qualitatively annotated based on preponderance of genes previously associated with the indicated cell states. (**C**) Differential expression plots (top 200 variable genes) for different niche branching lifecycle stage comparisons. As part of Xenium analysis, we pre-selected 100 genes (of the 5106 total) known to be relevant to nephrogenic niche development, which are indicated with crosses. (**D-F**) The same figure elements for ureteric bud cells neighboring the cap mesenchyme. (**G-I**) The same figure elements for stromal cells lining the niche.

210

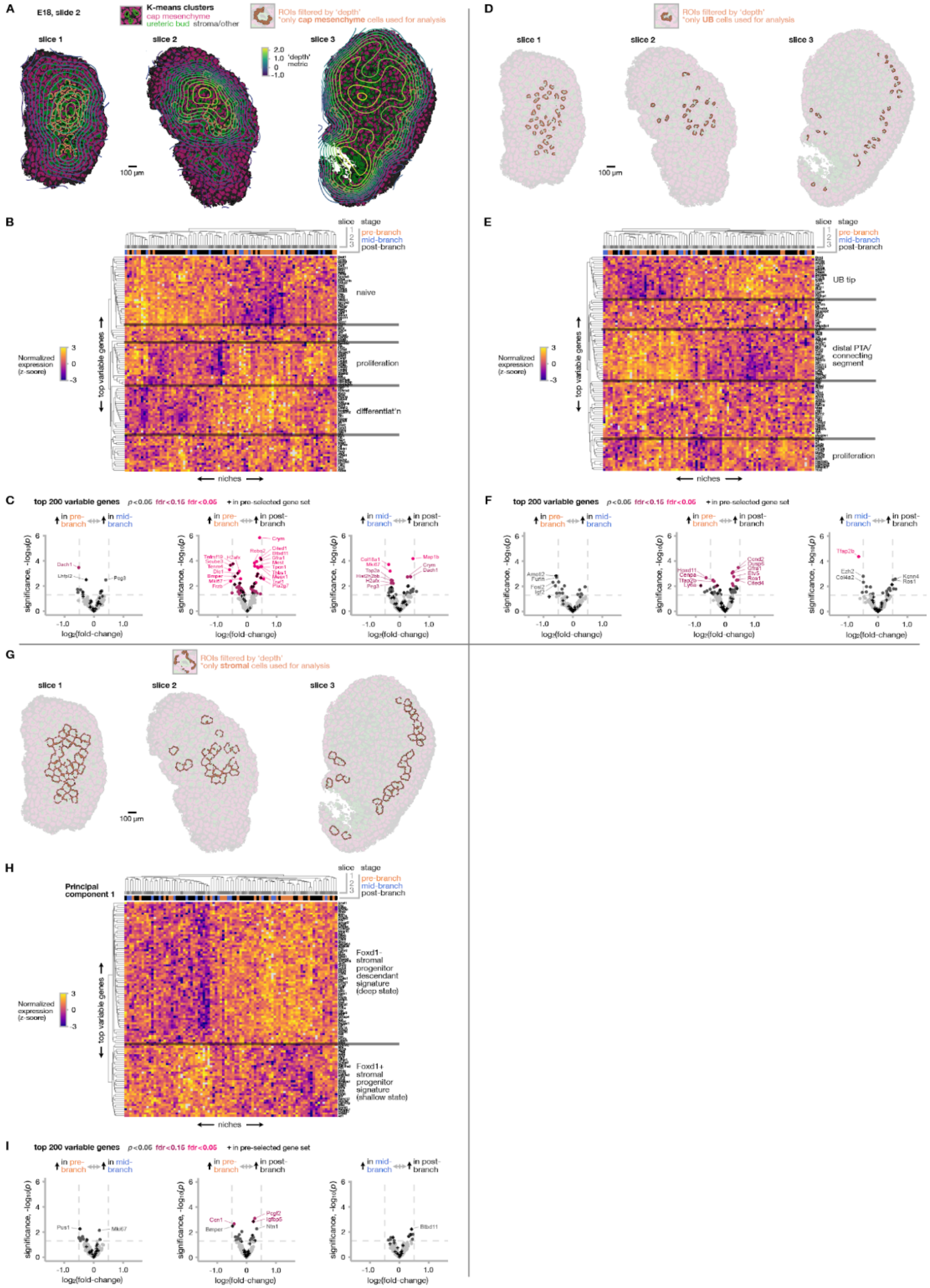


**Fig. S13: Mouse E18 (slide 2) nephrogenic niche selection and Xenium rhythmic gene expression analysis for niche cell compartments.** (**A**) Contours of the depth metric overlaid on k-means Xenium view, highlighting cap mesenchyme cells (nephron progenitors) in niches filtered for similar anatomical position in depth. (**B**) Unsupervised hierarchical clustering heatmap (top 100 variable genes) for pseudobulked transcript expression across niches at different branching lifecycle stages ($n$ = 91 niches across 3 slices from 3 different kidneys). Blocks of genes are qualitatively annotated based on preponderance of genes previously associated with the indicated cell states. (**C**) Differential expression plots (top 200 variable genes) for different niche branching lifecycle stage comparisons. As part of Xenium analysis, we pre-selected 100 genes (of the 5106 total) known to be relevant to nephrogenic niche development, which are indicated with crosses. (**D-F**) The same figure elements for ureteric bud cells neighboring
211 the cap mesenchyme. (**G-I**) The same figure elements for stromal cells lining the niche.

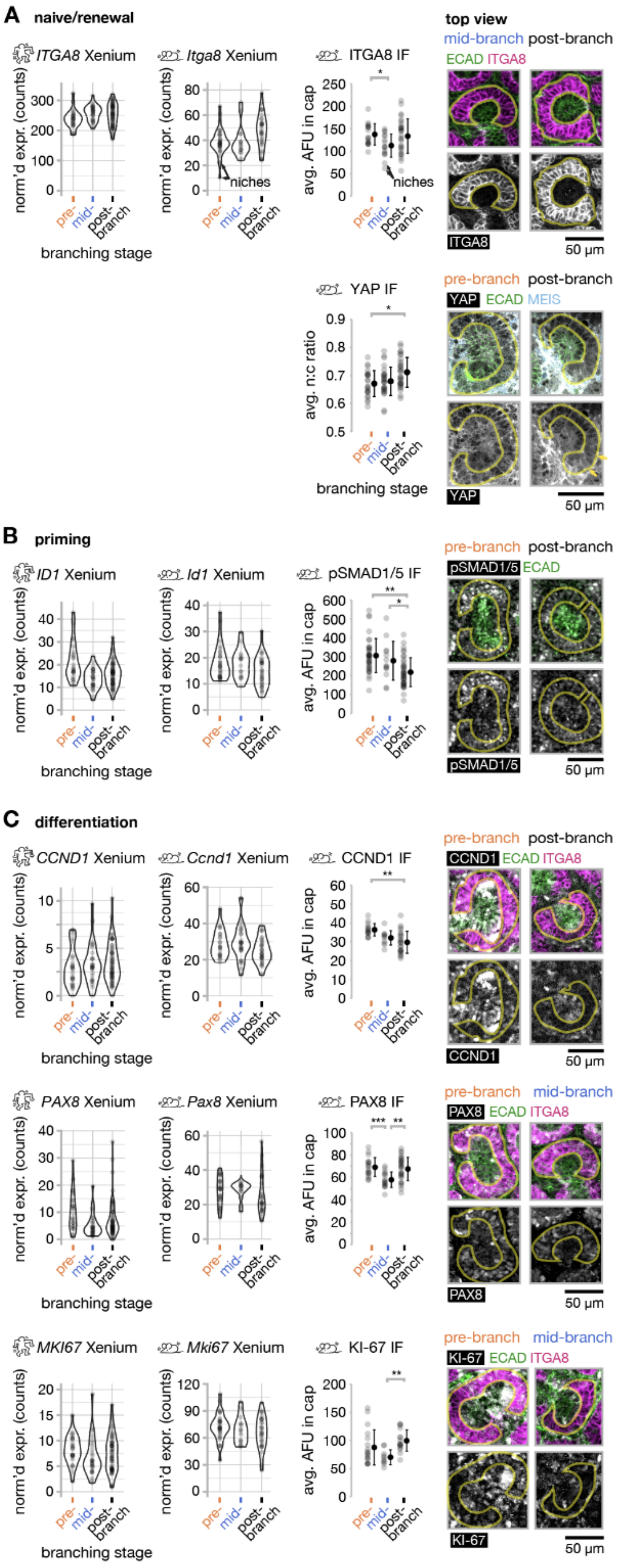


**Fig. S14: Rhythmic renewal and priming/differentiation markers in the cap mesenchyme over the niche branching life-cycle.** (**A-C**) Normalized Xenium expression of the indicated markers in human week 20 and mouse E17 cap mesenchymes, and average arbitrary fluorescence (AFU, or nuclear:cytoplasmic ratio for YAP) and example images of the same markers in mouse E17 cap mesenchymes by whole-mount confocal immunofluorescence at niche midplanes. Niches are grouped into pre-, mid-, and post-branching stages. Images at bottom show the isolated fluorescence channel for the marker of interest. Each point represents one cap mesenchyme niche, *n* > 12 niches per category, mean ± S.D. One-way ANOVA, Tukey's test, *$p < 0.05$, **$p < 0.01$, ***$p < 0.001$.

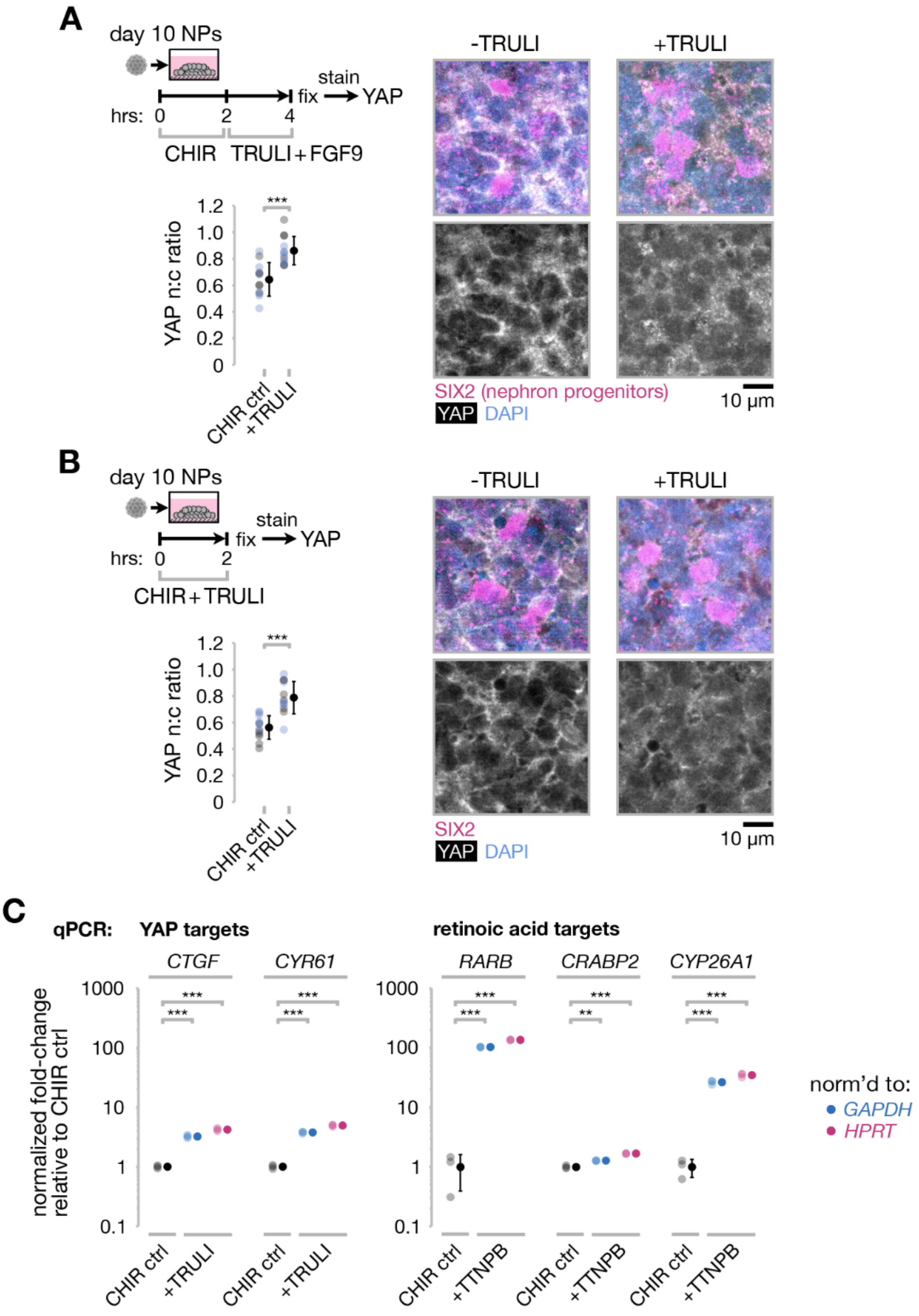


**Fig. S15: Validation of TRULI and TTNPB activity in human iPSC-derived nephron progenitor 'puck' organoids.** (**A**) *Left*, Plot of YAP nuclear-to-cytoplasmic (n:c) ratio (a metric for YAP activation) in day 10 nephron progenitor (NP) organoids fixed after staggered 7 µM CHIR and 4 µM TRULI treatments relative to CHIR-only controls ($n$ = 6 cells from each of two replicate pairs of organoids per comparison indicated by black and blue marker colors, mean ± S.D., unpaired $t$-test ***$p$ < 0.001). *Right*, representative immunofluorescence micrographs at approximate organoid mid-planes. (**B**) Similar data for overlapping 7 µM CHIR and 4 µM TRULI treatments relative to CHIR-only controls. (**C**) qPCR quantitation of YAP and retinoic acid transcriptional target expression in nephron progenitor organoids treated as in (B) for overlapping 7 µM CHIR and 4 µM TRULI or 0.1 µM TTNPB treatments relative to CHIR-only controls ($n$ = 3 reactions per condition, 8 organoids pooled per reaction, mean ± S.D., unpaired $t$-test **$p$ < 0.01, ***$p$ < 0.001). Expression was normalized to either *GAPDH* or *HPRT* housekeeping transcripts.

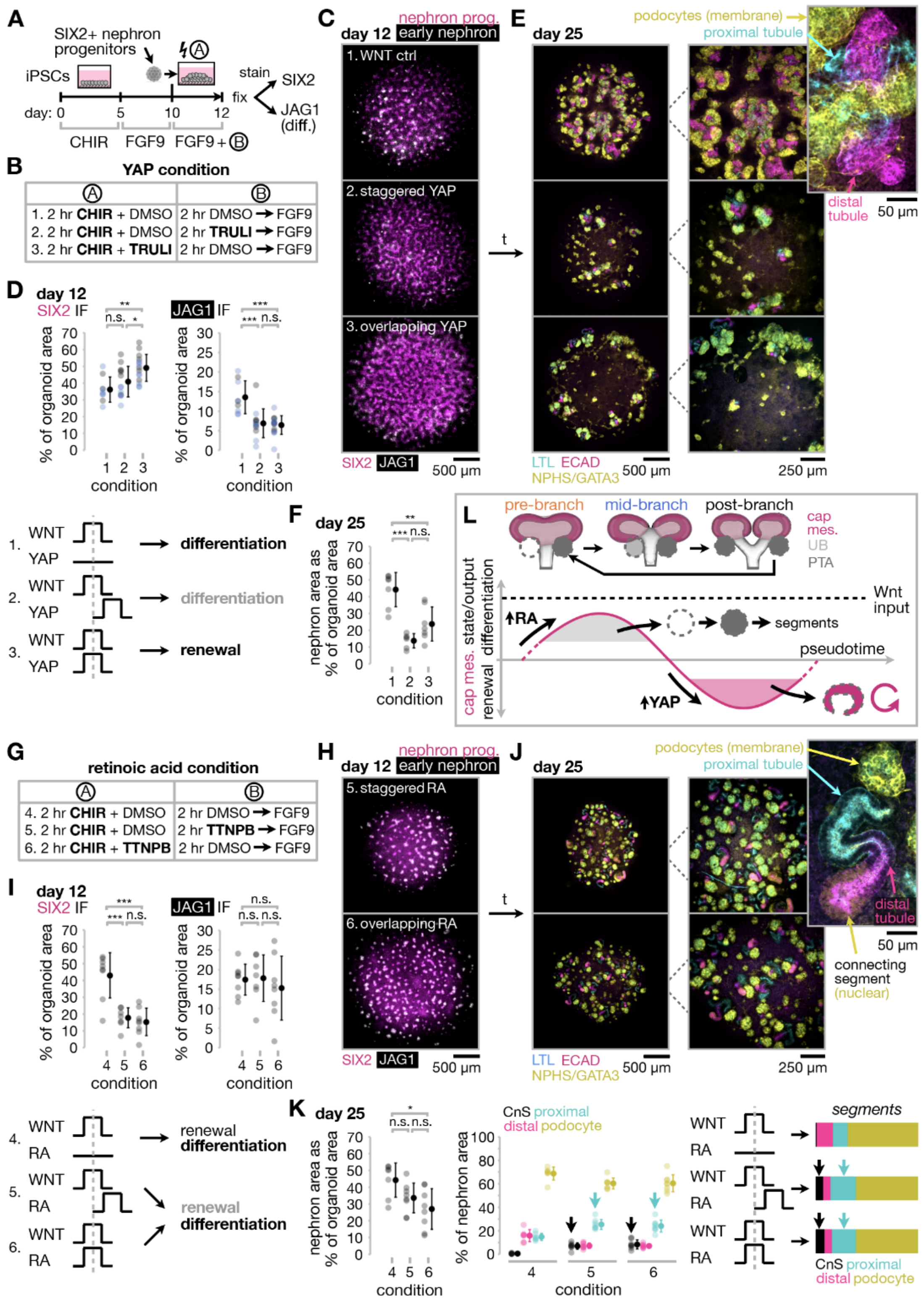


**Fig. S16: YAP and RA status modify iPSC-derived nephron progenitor interpretation of Wnt/β-catenin differentiation signaling in 'puck' organoids.** (**A**) Schematic workflow for nephron progenitor organoid differentiation and perturbation. (**B**) Table of perturbation conditions in the indicated phases of the differentiation schematic. Conditions 2 and 3 stagger or overlap the YAP agonist TRULI (4 µM) with standard 7 µM CHIR-induced differentiation in condition 1, respectively. (**C**) Confocal immunofluorescence images of representative $SIX2^{EGFP}$ iPSC-derived organoids at the day 12 endpoint for nephron progenitor (SIX2) and early nephron (JAG1) markers. (**D**) *Top*, Plots of marker expression as % of organoid area for two replicates, one for 300,000 cells per puck at day 10 (black markers, $MAFB^{BFP}$:$GATA3^{mCherry}$ iPSC line) and one for 150,000 cells per puck (blue markers, $SIX2^{EGFP}$ iPSC line), *n* > 2 pucks per condition per replicate, mean ± S.D. *Bottom*, schematic of experiment outcomes. (**E**) Confocal immunofluorescence images of representative $MAFB^{BFP}$:$GATA3^{mCherry}$ iPSC-derived organoids at the day 25 endpoint for nephron markers. (**F**) Plot of nephron cell area fraction relative to organoid area at day 25 for organoids in (E), *n* > 5 pucks per condition, mean ± S.D. (**G-J**) Similar data for $MAFB^{BFP}$:$GATA3^{mCherry}$ iPSC-derived organoids perturbed with the retinoic acid analogue TTNPB, *n* > 6 pucks per condition, mean ± S.D. (**K**) *Left*, Plot of nephron cell area fraction relative to organoid area at day 25 for organoids in (J). *Right*, plot and schematic of nephron composition by cell type. Arrows indicate increase in connecting segment (CnS) and proximal tubule cells in conditions treated with TTNPB. *n* > 6 pucks per condition, mean ± S.D. Statistics in D,F,I,K are one-way ANOVA, Tukey's test, *$p < 0.05$, **$p < 0.01$, ***$p < 0.001$. (**L**) Schematic model for differential interpretation of Wnt input by nephron progenitors depending on branching life-cycle-associated rhythms in YAP and RA signaling.

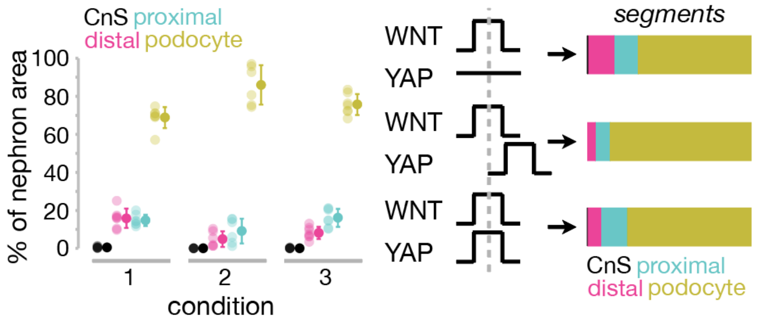


**Fig. S17: TRULI experiment organoid composition.** Plot and schematic of nephron composition by cell type for conditions 1-3 in **Fig. S16** at day 25, *n* > 5 pucks per condition, mean ± S.D.

216

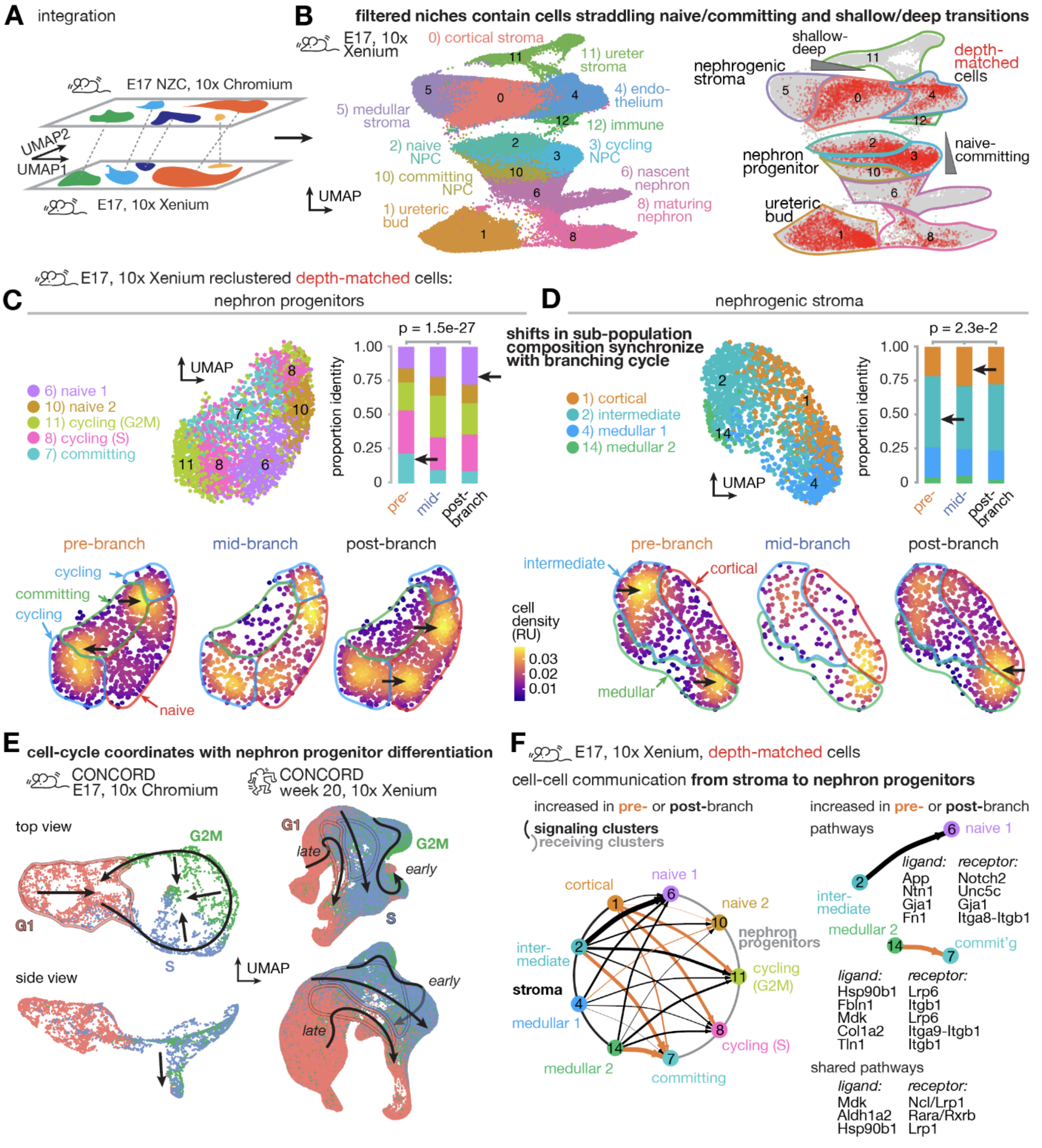


**Fig. S18: Mouse nephron progenitor and stromal rhythms reflect coordinated changes in niche sub-population composition.** (**A**) *Left*, Schematic of single-cell cluster label assignment to mouse E17 kidney Xenium data through integration. *Right*, UMAP representation of Xenium cells labeled by cluster and by membership in niches filtered for comparable anatomical location using depth metric contours. (**C,D**) Xenium cell UMAPs for re-clustered nephron progenitor and stromal lineages, and histograms and cell density heatmaps showing niche sub-population composition vs. discrete branching stage (1128, 527, and 1179 NP cells and 801, 232, and 756 stromal cells for pre-, mid-, and post-branching stages, respectively). Black arrows indicate notable differences in the data. Note that 'cortical', 'medullar', etc. labels relate to stroma in the nephrogenic zone, not kidney-wide. (**E**) UMAP representation of CONCORD latent space from nephron lineage identities in mouse E17 nephrogenic zone cell Chromium scRNA-seq and human week 20 Xenium spatial scRNA-seq datasets, respectively, colored by cell cycle phase. Arrows are qualitative trajectories. (**F**) CellChat prediction of cell-cell communication. *Left*, circle plot with signaling (sender) cell clusters at left and receiving clusters at right. Arrow thickness indicates relative differential interaction weight, arrow color indicates UB branching life-cycle stage with highest weight. *Right*, notable ligand/receptor pairs for selected cluster interactions.

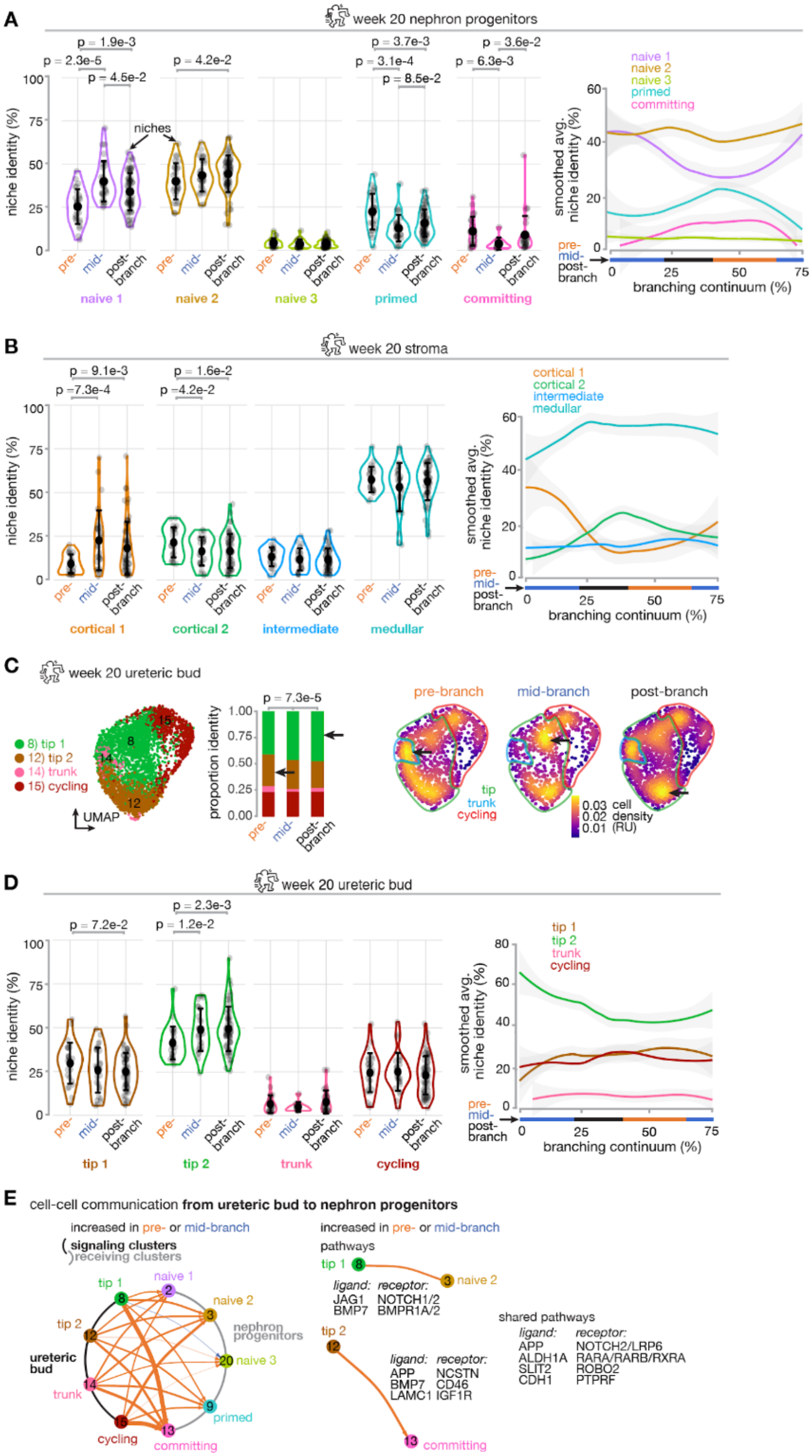


**Fig. S19: Human week 20 niche subpopulation composition and ureteric bud-to-nephron progenitor CellChat analyses.** (**A**,**B**) Dot plots and smoothed line plots of human week 20 niche sub-population composition vs. discrete branching stage and branching continuum score in nephron progenitor and stroma compartments where Xenium cells have been associated with their niches (points are niches, $n$ = 27, 29, 64 niches for pre-, mid-, post-branching stages across 5 kidneys, mean ± 95th CI, Wilcoxon rank sum testing). Note that 'cortical', 'medullar', etc. labels relate to stroma in the nephrogenic zone, not kidney-wide. (**C**) Xenium cell UMAPs for re-clustered mouse E17 ureteric bud lineage, and histogram ($n$ = 1237, 883, 1884 cells for pre-, mid-, post-branching stages) and cell density heatmaps showing niche sub-population composition vs. discrete branching stage. Black arrows indicate notable differences in the data. (**D**) Dot plots and smoothed line plots of human week 20 niche sub-population composition vs. discrete branching stage and branching continuum score in ureteric bud compartment where Xenium cells have been associated with their niches (points are niches, $n$ = 27, 29, 64 niches for pre-, mid-, post-branching stages across 5 kidneys, mean ± 95th CI, Wilcoxon rank sum testing). (**E**) CellChat prediction of cell-cell communication. *Left*, circle plot with signaling (sender) cell clusters at left and receiving clusters at right. Arrow thickness indicates relative differential interaction weight, arrow color indicates UB branching life-cycle stage with highest weight. *Right*, notable ligand/receptor pairs for selected cluster interactions.

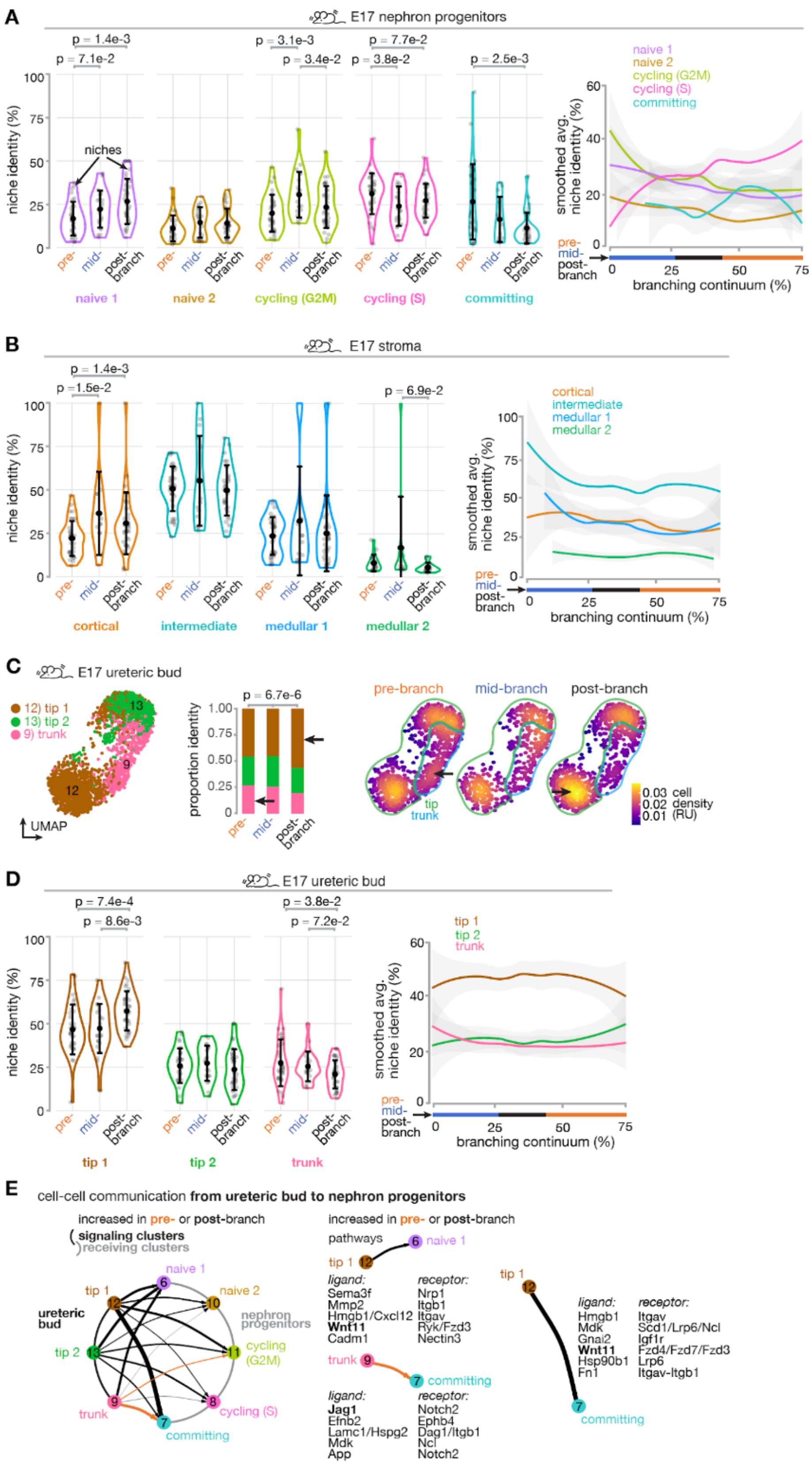


**Fig. S20: Mouse E17 niche subpopulation composition and ureteric bud-to-nephron progenitor CellChat analyses.** (**A,B**) Dot plots and smoothed line plots of mouse E17 niche sub-population composition vs. discrete branching stage and branching continuum score in nephron progenitor and stroma compartments where Xenium cells have been associated with their niches (points are niches, *n* = 34, 22, 36 niches for pre-, mid-, post-branching stages across 5 kidneys, mean ± 95th CI, Wilcoxon rank sum testing). Note that 'cortical', 'medullar', etc. labels relate to stroma in the nephrogenic zone, not kidney-wide. (**C**) Xenium cell UMAPs for re-clustered mouse E17 ureteric bud lineage, and histogram (*n* = 1027, 518, 919 cells for pre, mid, post) and cell density heatmaps showing niche sub-population composition vs. discrete branching stage. Black arrows indicate notable differences in the data. (**D**) Dot plots and smoothed line plots of mouse E17 niche sub-population composition vs. discrete branching stage and branching continuum score in ureteric bud compartment where Xenium cells have been associated with their niches (points are niches, *n* = 34, 22, 36 niches for pre-, mid-, post-branching across 5 kidneys, mean ± 95th CI, Wilcoxon rank sum testing). (**E**) CellChat prediction of cell-cell communication. *Left*, circle plot with signaling (sender) cell clusters at left and receiving clusters at right. Arrow thickness indicates relative differential interaction weight, arrow color indicates UB branching life-cycle stage with highest weight. *Right*, notable ligand/receptor pairs for selected cluster interactions.

220

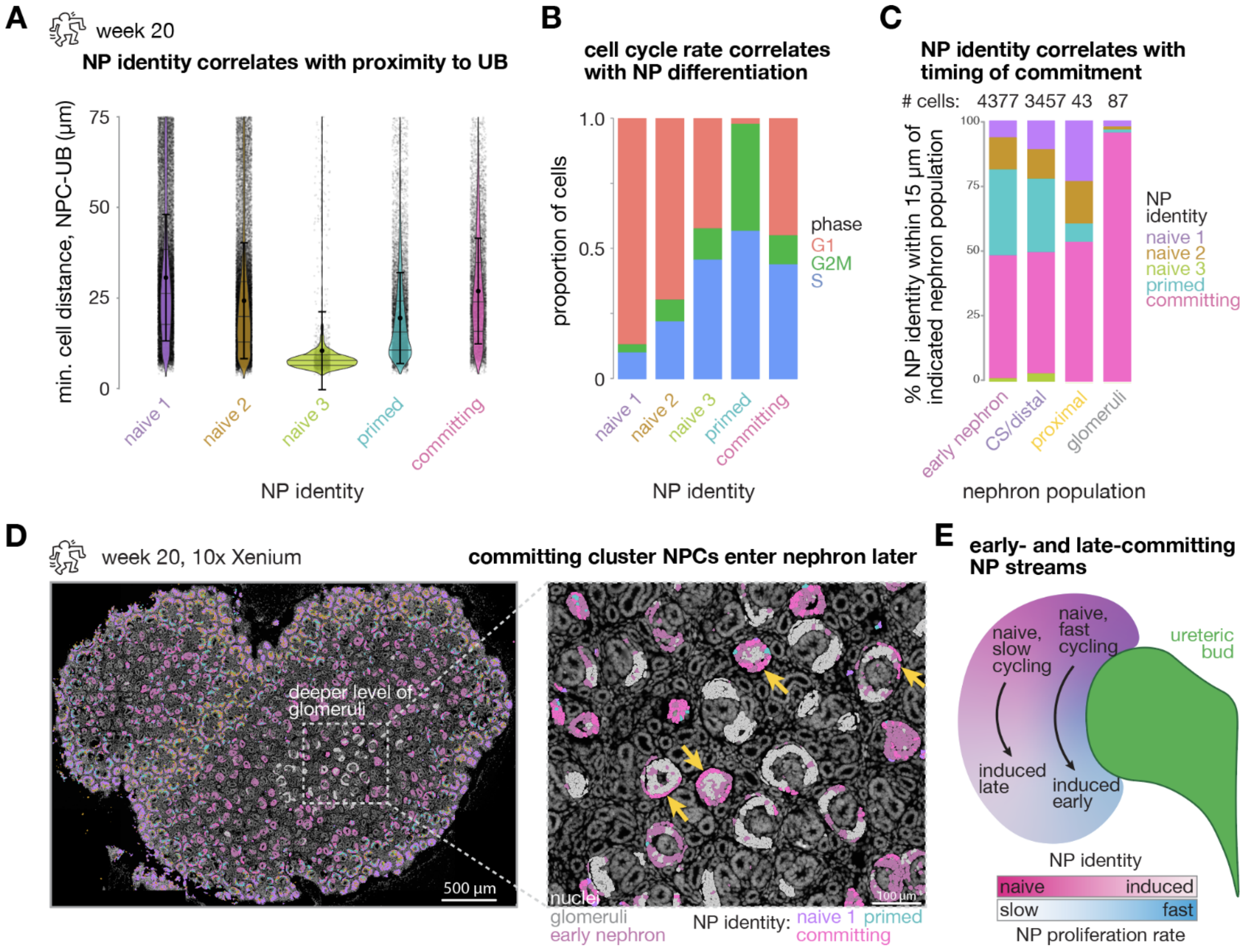


**Fig. S21: Cell cycle rate correlates with progressive recruitment of nephron progenitors in UB-adjacent and distant streams.** (**A**) Minimum cell distance to the nearest ureteric bud (UB) cell for nephron progenitor (NP) cells in the clusters indicated on the *x* axis in week 20 human kidney Xenium slice. (**B**) Predicted proportion of cells in the indicated cell cycle phases. (**C**) Niche composition within a 15 μm neighborhood local to cells in the nephron-lineage populations indicated on the *x* axis. (**D**) Week 20 kidney slice and zoom inset showing all cells by DAPI staining (gray), and colored overlays by cell identity. Yellow arrows indicate committing cluster cells peripheral to renal corpuscles. (**E**) Diagram of inferred nephron progenitor streams, temporal and geometric relationship in the niche, and coordination between differentiation and cell cycle rate.

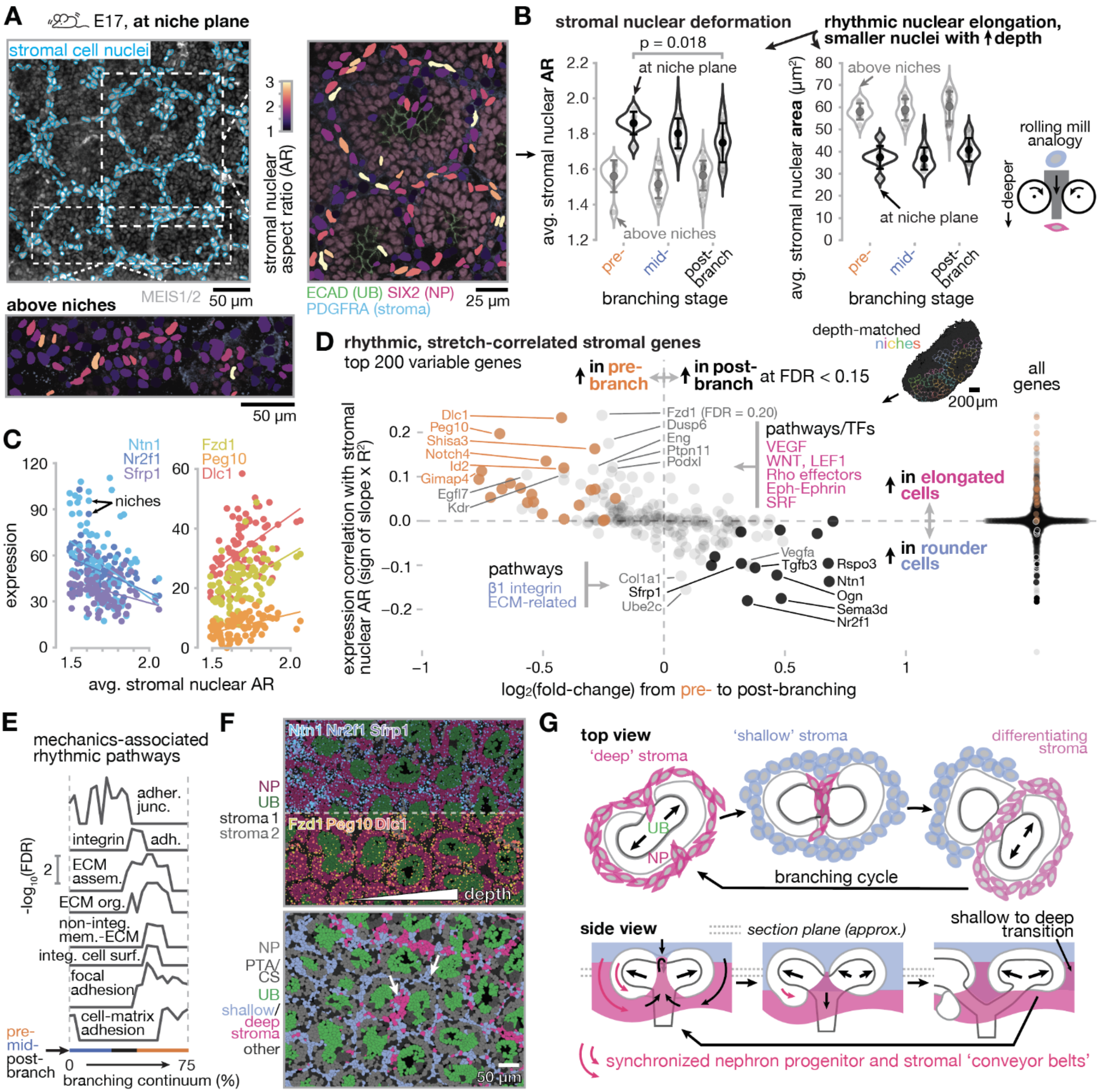


**Fig. S22: The stromal rhythm correlates with a biophysical transition during differentiation and descent of coordinated 'conveyor belts' of stroma and nephron progenitors.** (**A**) *Top*, Cellpose outlines of stromal nuclei from mouse E17 kidney confocal immunofluorescence slice, insets show stromal nuclear aspect ratio detail relative to other niche compartments at their midplane and at a higher *z* plane on top of niches. (**B**) Plots of average stromal nuclear aspect ratio and area vs. niche branching stage (mean ± s.d.; $n$ = 40 niches across 5 fields of view across 2 kidneys). (**C**) Plots of nuclear aspect ratio-correlated transcripts. (**D**) Plot of transcript correlation with nuclear aspect ratio vs. degree of rhythmic variation over the branching lifecycle. Transcripts in the top left quadrant are enriched in pre-branching niches and stromal cells with higher aspect ratio. Notable ToppGene pathways and transcription factor binding sites associated with AR-correlated transcripts are listed. (**E**) Line plots of GO/KEGG/reactome analyses performed using a sliding window over rhythmic genes fitting a sine wave model at $p < 0.2$ and ordered by peak position along the branching continuum axis (adher. junc., adherens junction; integrin adh., cell adhesion mediated by integrin; ECM assem., extracellular matrix assembly; ECM org., extracellular matrix organization; non.-integ. mem.-ECM, non-integrin membrane-ECM interactions; integ. cell surf., integrin cell surface interactions). (**F**) *Top*, Xenium view of notable hits from (**D**) overlaid on niche compartments colored by k-means cluster; point number and size indicate relative transcript density. *Bottom*, Stromal cells in the same field, colored by shallow/deep state scored by gene set expression. (**G**) Schematic model of rhythmic changes in stromal cell biophysical and differentiation state local to the niche over branching lifecycle.
224
225

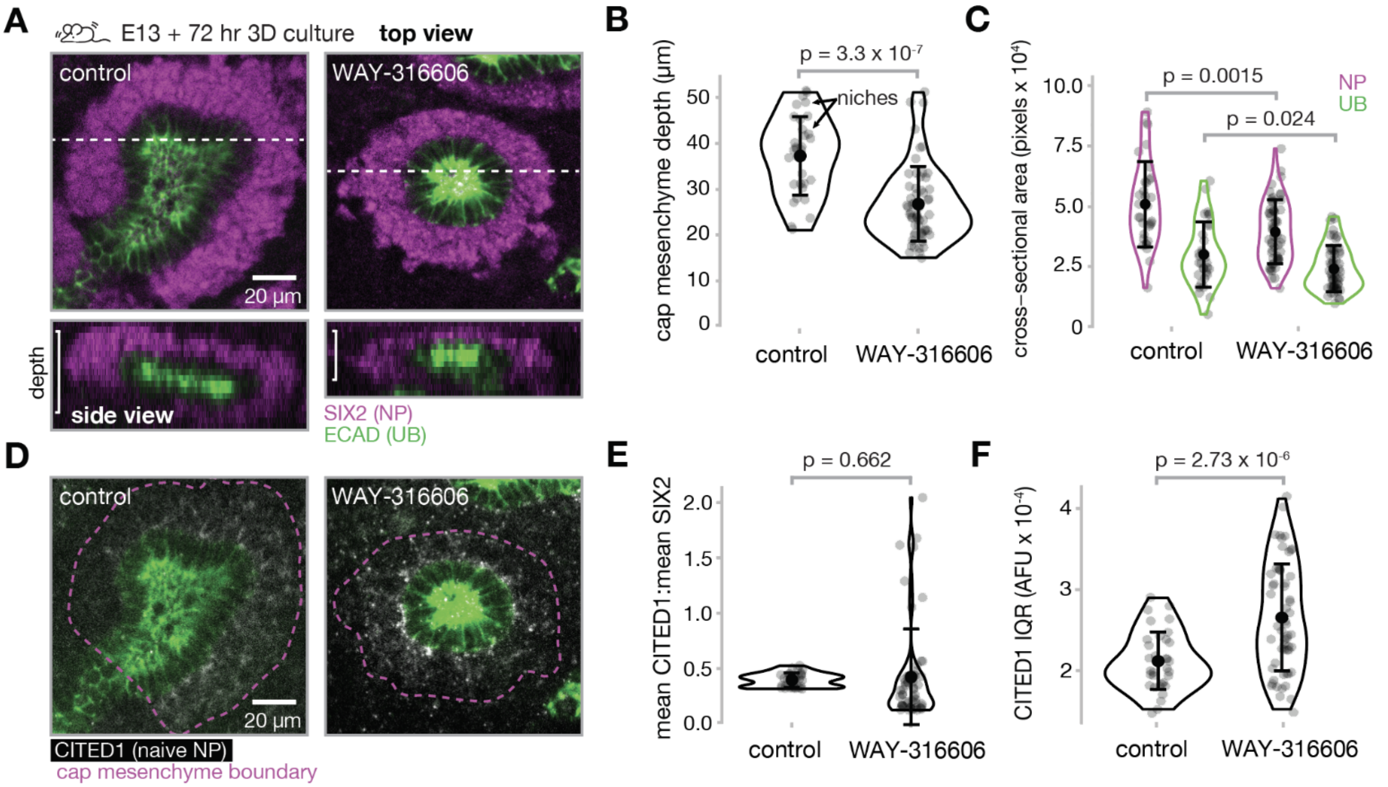


**Fig. S23: Stroma-derived SFRP1 is necessary for nephron progenitor maintenance.** (**A**) Representative confocal fluorescence micrographs of cap mesenchyme (nephron progenitor compartment) spatial extent after 72 hr embryonic kidney explant culture in presence and absence of the SFRP1 inhibitor WAY-316606 (*n* = 33 niches across 2 kidneys for control and 57 niches across 3 kidneys for WAY-316606). (**B**) Plot of cap mesencyme *z* depth vs. treatment group. (**C**) Plots of per-niche nephron progenitor and ureteric bud tip cell cross-sectional areas in *xy* vs. treatment group. (**D**) Representative micrographs of CITED1 naive nephron progenitor marker distribution in the cap mesenchyme vs. treatment group. Note CITED1 retention near the ureteric bud interface with a decreasing radial gradient outward in the cap mesenchyme. (**E**) Ratio of mean cap mesenchyme CITED1:SIX2 fluorescence intensity vs. treatment group. SIX2 is a marker of all nephron progenitors. (**F**) Interquartile range (IQR) in CITED1 fluorescence intensity among pixels within each niche, capturing the CITED1 gradient that appears in WAY-316606-treated niches. Plot error bars represent mean ± SD.

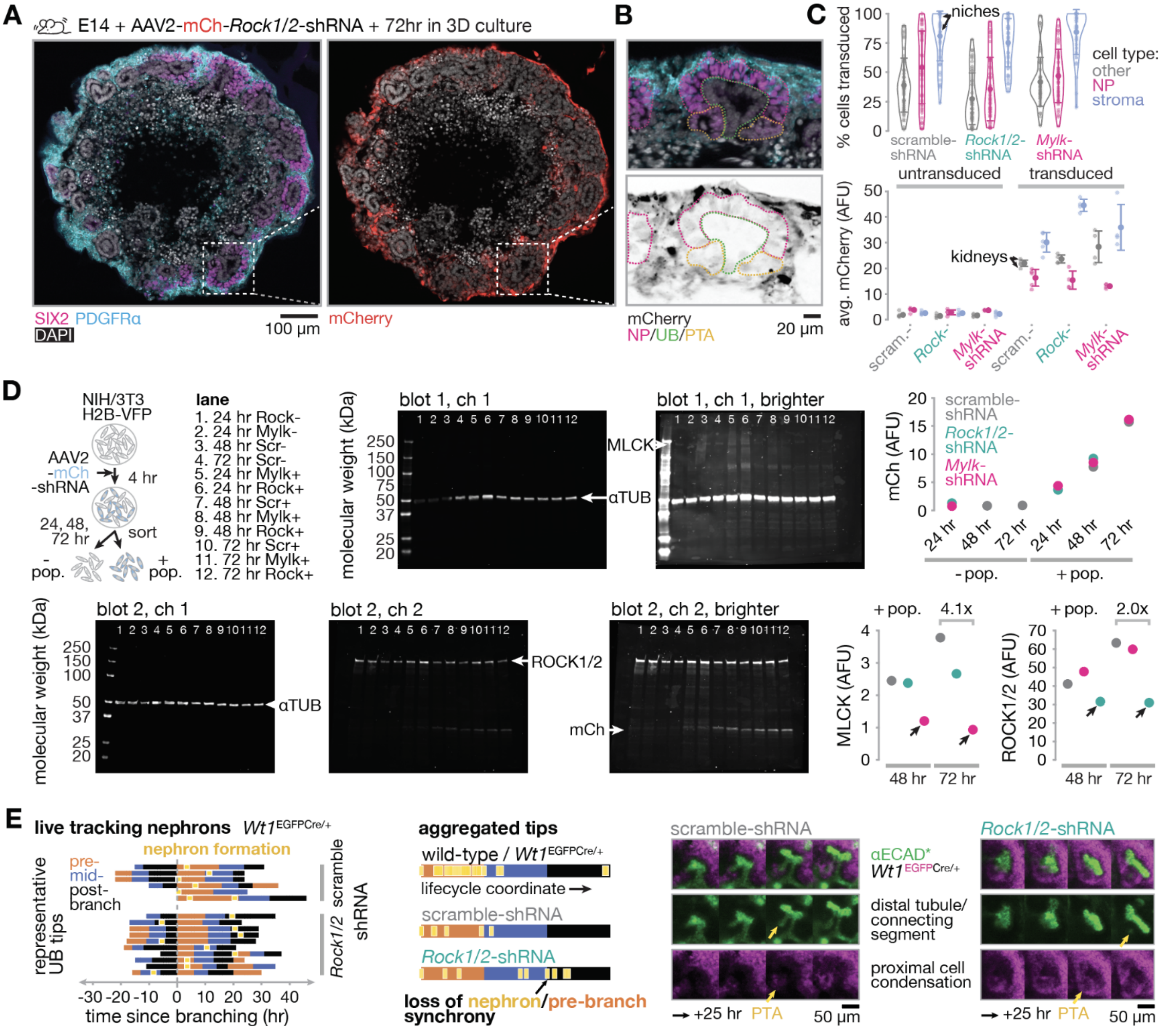


**Fig. S24: Validation data for mouse embryonic kidney transduction with AAV2 and additional nephron formation tracking data.** (**A**) Fixed slice immunofluorescence images for nephrogenic zone markers and mCherry transduction marker. (**B**) Inset views detailing enriched expression in the cortical nephrogenic zone stroma relative to nephron progenitor (NP), ureteric bud (UB), and early nephron (pretubular aggregate, PTA) compartments. (**C**) Quantification of cell transduction efficiency and expression level by cell segmentation ($n$ = 47, 47, and 44 niches across 5, 4, 4 kidneys from scramble-shRNA, Rock1/2-shRNA, and Mylk-shRNA treated kidneys). (**D**) Validation of AAV2 knockdown efficacy by western blot. *Left*, cartoon and sample key for NIH/3T3 infection and sorting for positive ('+ pop.') and negative ('- pop.') fractions at 24, 48, and 72 hr timepoints. *Middle*, Full western blots with lane numbers corresponding to the sample key. *Right*, Densitometry plots of mCherry (mCh) transduction marker (*top*) and MLCK and ROCK1/2 protein (*bottom*) expression. Arrows indicate conditions in which the knockdown hairpin matched the plotted target. Fold-changes are listed relative to scramble-shRNA controls. (**E**) *Left*, Ticker-tape plots of nephron formation events in *Wt1*$^{EGFPCre/+}$ kidneys relative to ureteric bud tip branching stage after aligning each tip to a common time since branching initiation; $n$ = 7 and 10 tips across 2 kidneys each for scramble-shRNA and *Rock1/2*-shRNA conditions respectively. *Middle*, Aggregated ticker-tapes in which nephron formation events are plotted at the fraction of the corresponding branching stage that they formed in. Stages are idealized as equal thirds of the branching life-cycle. Data from uninfected control kidneys are reproduced for completeness. *Right*, timelapse frames capturing representative nephron formation dynamics in scramble-shRNA and *Rock1/2*-shRNA treated kidneys. Yellow arrows indicate proper distal (ECAD+) and proximal (EGFP+) cell recruitment into PTAs in each case.

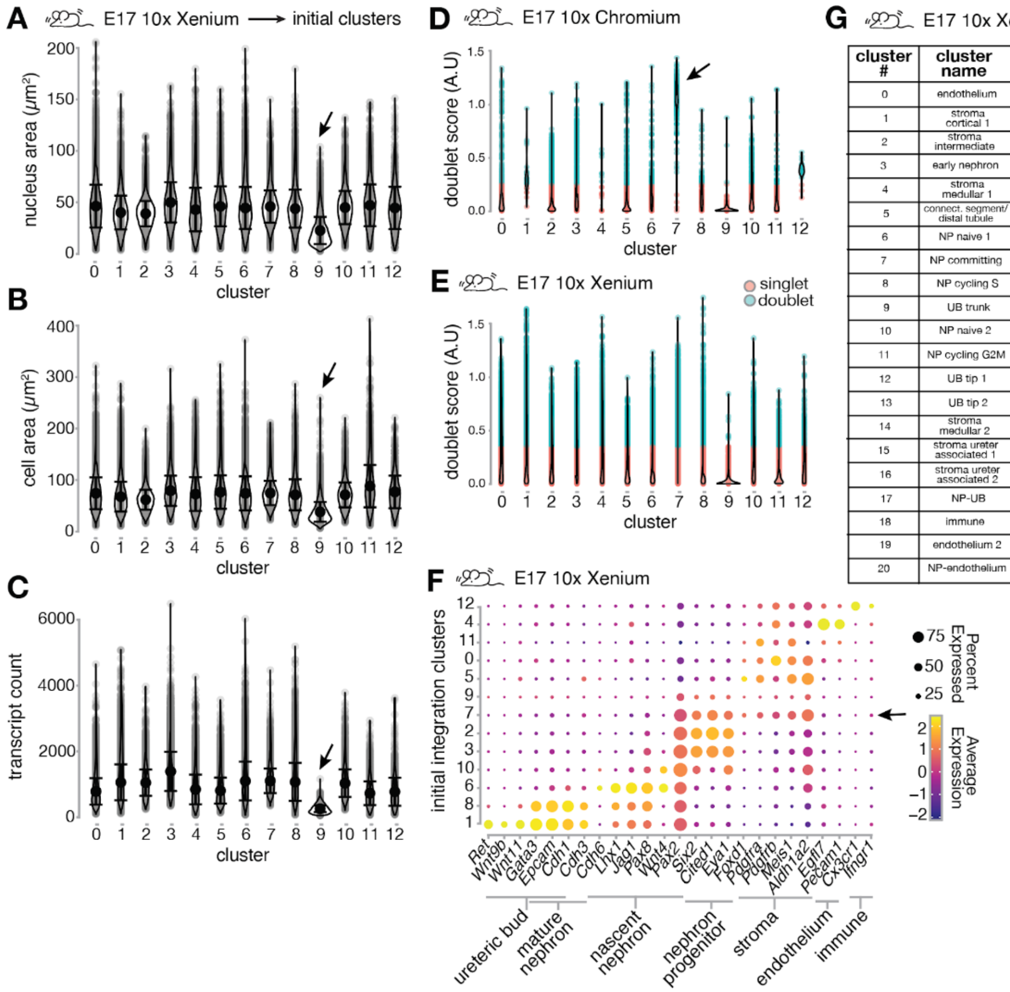


**G** E17 10x Xenium ⟶ refined clusters

| cluster # | cluster name | top 20 differentially expressed genes |
|---|---|---|
| 0 | endothelium | Egfl7, Podxl, Cdh5, Pecam1, Plxnd1, Gimap4, Sox18, Igfbp3, Tie1, Kdr, Eng, Ets1, Aplnr, Arhgef15, Dysf, Ednrb, She, Mcam, Epas1, Plk2 |
| 1 | stroma cortical 1 | Ntn1, Sfrp1, Col1a1, Arap1, Hic1, Dact1, Cldn1, Col1a2, Plagl1, Aox3, Slc8a1, Meis1, Cdkn1c, Aldh1a2, Smoc2, Dhrs3, Pdgfra, Postn, Bnc2, Ogn |
| 2 | stroma intermediate | Hic1, Meis1, Emilin1, Dact1, Col1a1, Gdnf, Col1a2, Fhl2, Aldh1a2, Sfrp1, Ntn1, Anxa2, Arap1, Pdgfra, Fn1, Plagl1, Pdgfrb, Slc8a1, Nfia, Smtn |
| 3 | early nephron | Clu, Lhx1, Jag1, Wnt4, Pax8, Ptpn13, Osr2, Ncs1, Dach1, Gja1, Ccnd1, Svopl, Arl4c, Clic4, Rai14, Dll1, Scube3, Hey1, Celsr2, Uncx |
| 4 | stroma medullar 1 | Col1a2, Pdgfrb |
| 5 | connect. segment/ distal tubule | Cdh1, Epcam, Ly6a, Scube3, Tfap2a, Dact2, Emx2, Gata3, Ptpn13, Pou3f3, Col18a1, Ldhb, Arl4c, Tfap2b, Cldn7, Dach1, Pax8, Slit1, Emx1, Adgrg1 |
| 6 | NP naive 1 | Cited1, Traf1, Robo2, Btbd11, Six2, Spock2, Vcan, Dst, Uncx, Eya1, Sema4g, Meox1, Rspo1, Dock5, Hmcn1, Gas1, Amph, Itga8, Col2a1, Unc5c |
| 7 | NP committing | C1qtnf12, Bmper, Col2a1, Dlc1, Wt1, Wnt4, Spock2, Scube3, Tenm4, Slit1, Myc, Pdgfc, Id2, Pax2, Eya1, Tnfrsf19, Ncam1, Lhfpl2, Rdh10, Nr2f1 |
| 8 | NP cycling S | Cited1, Spock2, Six2, Robo2, Btbd11, Uncx, Eya1, Vcan, Rrm2, Shroom3, Dst, Gas1, Col2a1, Wt1, Itga8, Pdgfc, Traf1, Uhrf1, Amph, Sall1 |
| 9 | UB trunk | Gata3, Tbx3, Epcam, Calb1, Cdh1, Lama5, Efnb2, Wwc1, Tfap2a, Wnt9b, Ldhb, Dact2, Fgfr2, Adgrg1, Cldn7, Emx2, Col18a1, Glis3, Gfra1, Sox9 |
| 10 | NP naive 2 | Cited1, Spock2, Btbd11, Traf1, Robo2, Vcan, Six2, Gas1, Capn6, Col2a1, Ttc28, Uncx |
| 11 | NP cycling G2M | Cited1, Six2, Uncx, Spock2, Btbd11, Robo2, Eya1, Rspo1, Traf1, Vcan, Meox1, Shroom3, Cenpa, Dst, Gas1, Amph, Wt1, Col2a1, Cdc20, Sulf2 |
| 12 | UB tip 1 | Wnt11, Ret, Sox9, Calb1, Epcam, Gata3, Vldlr, Cdh1, Tfap2a, Sema6a, Etv5, Gfra1, Dact2, Irs2, Hoxb9, Ppfibp1, Pou3f3, Ldhb, Spry1, Cldn7 |
| 13 | UB tip 2 | Gata3, Calb1, Epcam, Sox9, Cdh1, Gfra1, Tbx3, Ldhb, Tfap2a, Ret, Wnt11 |
| 14 | stroma medullar 2 | Pdgfrb, Mcam, Gucy1a1, Abcc9, Heyl, Plxnd1, Afap1l2, Cspg4, Epas1, Tbx2, Col1a2, Atp1b2, Zeb2, Cd248, Kcnj8, Plekhg2, Fhl2, Peg10, Col4a1, Vim |
| 15 | stroma ureter associated 1 | Dcn, Postn, Lum, Col1a1, Col6a2, Col5a1, Col6a3, Col14a1, Col6a1, Prrx1, Ogn, Cd248, Col1a2, Sfrp1, Igf1, Sfrp4, Agtr2, Flnc, Lox, Tbx18 |
| 16 | stroma ureter associated 2 | Myh11, Actg2, Dcn, Tagln, Actc1, Lum, Tbx18, Gata2, Myocd, Cnn1, Col6a3, Col6a2, Col1a1, Tgfb1i1, Nkd1, Sfrp2, Ckb, Gata5, Jph2, Col6a1 |
| 17 | NP-UB | Wnt11, Gfra1, Cited1, Sox9, Calb1, Ret, Capn6, Epcam, Hoxb9, Etv5, Btbd11, Tfap2a, Gata3, Traf1, Hoxb3, Hoxb8, Vldlr, Ldhb, Cdh1, Pax2 |
| 18 | immune | Cx3cr1, Maf, Csf1r, Mrc1, Sirpa, F13a1, Spi1, Adgre1, C3ar1, Lcp2, Aif1, Dab2, Arrb2, Slc40a1, Zeb2 |
| 19 | endothelium 2 | Dcn, Cdh5, Pecam1, Eng, Aplnr, Epas1, Lum, Esam, Clec14a, Cav1, Tbx18, Rgcc, Entpd1, Egfl7, Sox18, Hecw2, Igf1, Prrx1, Postn, Kdr |
| 20 | NP-endothelium | Traf1, Cited1, Btbd11, Pecam1, Gimap4, Spock2, Podxl, Tril, Cdh5, Gas1, Igfbp3, Vcan, Six2, Col2a1, Hmcn1 |

**Fig. S25: mouse kidney single cell RNA-seq cluster quality control and identification.** (**A-F**) Initial clustering assignments after integration of mouse E17 Xenium with the mouse E17 nephrogenic zone cell 10x Chromium dataset. (**A**) nucleus area, (**B**) cell area, and (**C**) transcript count for all mapped cells per cluster. Arrows highlighting cluster 9 as an outlier based on cell size and transcript count. (**D,E**) Doublet score evaluated by scDblFinder's computeDoubletDensity function for all mapped cells per cluster. Cells are colored by their doublet estimation determined by scDblFinder's doubletThresholding function. (**D**) Doublet estimation from 10x Chromium nephrogenic zone datasets. Arrow showing cluster 7 as a likely doublet. (**E**) Doublet estimation from 10x Xenium datasets. (**F**) Marker gene dot plot with size representing percentage of expressing cells in each cluster and color representing average scaled expression within each cluster. Marker genes for ureteric bud (*Ret*, *Wnt9b*, *Wnt11*, *Gata3*, *Epcam*, *Cdh1*, *Cdh3*), mature nephron (*Gata3*, *Epcam*, *Cdh1*, *Cdh3*), nascent nephron (*Cdh6*, *Wnt4*, *Lhx1*, *Jag1*, *Pax8*, *Pax2*), nephron progenitor (*Pax2*, *Six2*, *Cited1*, *Eya1*), stroma (*Foxd1*, *Pdgfra*, *Pdgfrb*, *Meis1*, *Aldh1a2*), endothelium (*Egfl7*, *Pecam1*), and immune (*Cx3cr1*, *Ifngr1*). Arrow highlighting cluster with expression consistent with possible contribution from both nephron progenitor and stromal cells. (**G**) Table showing top differentially expressed genes for clusters from the refined cluster assignments. Column 1 shows cluster number, column 2 shows annotated cluster identity, and column 3 are top DEGs.

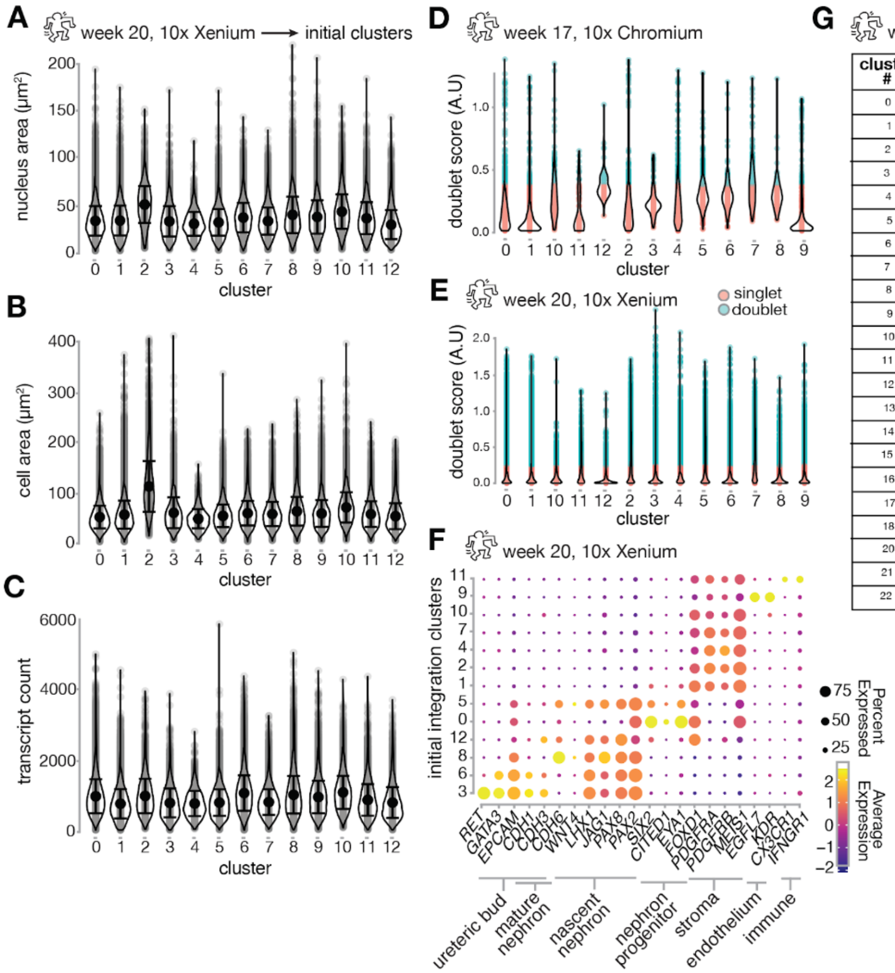


**G** week 20, 10x Xenium → refined clusters

| cluster # | cluster name | top 20 differentially expressed genes |
|---|---|---|
| 0 | stroma cortical 1 | TGFBI, CYP1B1, NTN1, SULF2, IGFBP5, NDNF, DNAH11, COL3A1, MFAP4, SFRP1, ITGA4, OLFML3, SMOC2, POSTN, CCBE1, DHRS3, ITM2A, PDGFRA, FAP, ADAMTS5 |
| 1 | stroma medullar | CXCL12, PDGFRA, POSTN, ITGA4, CYP1B1, OLFML3, TPBG, TGFBI, DHRS3, SNAI2, PDGFRB, GATA6, THBS1, FST, CCN4, VCAM1, PBX1, MFAP4, GDNF, ADAMTS5 |
| 2 | NP naive 1 | PCDH15, BMPER, SIX2, EYA1, MEOX1, TMEM100, ITGA8, PDGFC, NNAT, COL2A1, ECEL1, OSR1, WASF3, CHRNA1, FAT3, CDH24, TNFRSF19, CRABP2, FOXC1, STXBP5 |
| 3 | NP naive 2 | TMEM100, PCDH15, EYA1, MEOX1, ITGA8, SIX2, BMPER, NNAT, WASF3, ECEL1, PDGFC, COL2A1, UNC5B, CRABP2, OSR1, HAS2, ROBO2, SALL1, CDH24, TNFRSF19 |
| 4 | stroma cortical 2 | TCF21, CCN4, SNAI2, THBS1, EDNRA, S1PR3, PDGFRB, COL3A1, PDGFRA, OLFML3, TNC, TGFBI, PALLD, COL5A2, FBLN5, PLEKHH2, PCDH18, COL5A1, ABCC9, ADGRA2 |
| 5 | connect. segment/ distal tubule | HNF1B, MAL, EMX2, LHX1, COBLL1, PAX8, PKHD1, TMEM72, KCNJ15, TFAP2A, POU3F3, FMN1, EPCAM, MST1, SLIT2, COL4A4, GATA3, PAX2, COL4A3, ITGA2 |
| 6 | early nephron | PAX8, LYPD1, JAG1, LHX1, SFRP2, PLOD2, CCND1, CDH6, SORL1, DNM1, HES4, HEY1, HIST1H1A, HAS2, TOP2A, NOTCH1, UHRF1, PAX2, ID1, CENPU |
| 7 | stroma intermediate | CENPK, ZWINT, AURKB, CENPU, TOP2A, TYMS, BRIP1, EZH2, NUSAP1, CDK1, LMNB1, PDGFRA, ARHGAP11B, THBS1, H2AFX, SMC4, FOXM1, TPX2, MELK, FANCI |
| 8 | UB tip 1 | RET, ALDH1A1, TFAP2A, EMX2, LHX1, MYO6, AKR1B1, SCIN, FREM2, KCNC3, GATA3, SOX9, PKHD1, PLEKHA1, PAX8, GRHL2, EPCAM, HELLS, NPNT, COBLL1 |
| 9 | NP primed | HAS2, EYA1, SIX2, ITGA8, TMEM100, NNAT, PHF19, COL2A1, LYPD1, LTBP1, CRABP2, PCDH15, PDGFC, WASF3, UHRF1, TYMS, TOP2A, ECEL1, CDH24, CENPU |
| 10 | proximal tubule | CDH6, HNF1B, DPP4, TSPAN12, FGFR3, PDZK1, JAG1, GLYATL1, LRP2, EMX2, EPHA7, PAX8, TRPM3, FLRT3, ID1, LIPG, LRATD2, ACAA2, GGT1, FGFR4 |
| 11 | endothelial | TIE1, CD34, PLVAP, PECAM1, SHANK3, APLNR, FLT4, TEK, PRCP, KDR, EGFL7, NOTCH4 CDH5, HSPG2, ROBO4, STAB1, PODXL, SH2D3C, NOS3, TCIM |
| 12 | UB tip 2 | ALDH1A1, RET, EMX2, MYO6, LHX1, GATA3, PKHD1, TFAP2A, SCIN, EPCAM, KCNC3, PAX8, AKR1B1, COBLL1, PAX2, MAL, EPOR, SOX9, HNF1B, TRIB2 |
| 13 | NP committing | LYPD1, COL2A1, LTBP1, EYA1, HAS2, ITGA8, BMPER, DAPL1, SIX2, NNAT, MSI1, PEG10, PAX2, DNM1, ITPR1, CDH6, CDH24, TMEM100, PDGFC, PAX8 |
| 14 | UB trunk | GATA3, PKHD1, KCNC3, COBLL1, ALDH1A1, EMX2, MAL, SCIN, LHX1, MYO6, PAX8, EPOR, HNF1B, RET, NPNT, GPRC5B, TFAP2A, EPCAM, CDH16, EHF |
| 15 | UB tip cycling | RET, ALDH1A1, TFAP2A, EMX2, MYO6, SCIN, LHX1, AKR1B1, GATA3, FREM2, HELLS, EPCAM, COBLL1, PAX8, GRHL2, PKHD1, PLEKHA1, SOX9, TOP2A, ST14 |
| 16 | stroma | DCN, SULF1, TNXB, SFRP2, ADAMTS2, LOX, EMP1, SEMA3C, LTBP2, ITM2A, TWIST1, ELN, ADAMTSL3, CCDC80, SMOC2, SULF2, CCN1, IGF1, SFRP4, COL5A1 |
| 17 | immune | STAB1, CSF1R, CD14, SLCO2B1, TGFBI, ITGB2, CYBB, MRC1, CD4, SLC40A1, MAF, CIITA, SAMHD1, ARRB2, GAS6, CX3CR1, MSR1 |
| 18 | glomeruli | NPHS1, NPHS2, MAFB, PODXL, OLFM3, BCAM, PLOD2, CCN2, PAX8, WT1, SFXN2, FYN, BMP7, VEGFA, SULF1, LOXL1, CLDN1, KCNC3, NES, ITGA3 |
| 20 | NP naive 3 | ALDH1A1, TMEM100, NNAT, EYA1, ITGA8, SIX2, WASF3, RET, MEOX1, HAS2, ECEL1, PCDH15, CRABP2, AKR1B1, PDGFC, TNFRSF19, ETV5, COL2A1, PAX2, BMPER |
| 21 | UB tip 3 | TNF, ICAM1, CD40, CFB, RELB, PLAU, MACC1, SOCS3, NFKB2, TNFRSF11B, CDH1, ICOSLG, NFE2L3, RET, LAD1, ST14, CXCL16, TRIM47, BCL3, MYO6 |
| 22 | proximal tubule 2 | CDH6, DPP4, TRPM3, TSPAN12, HES4, FGFR4, LRP2, PDZK1, OSR2, LAMP5, GLYATL1, HNF4A, HNF4G, TRPC6, LIPC, EPHA7, HMMR, CDCA3, DLL1, FGFR3 |

**Fig. S26: human kidney single cell RNA-seq cluster quality control and identification.** (**A-F**) Initial clustering assignments after integration of human week 20 Xenium with the human week 17 nephrogenic zone cell 10x Chromium dataset from Lindstrom *et. al* 2018. (**A**) nucleus area, (**B**) cell area, and (**C**) transcript count for all mapped cells per cluster. (**D,E**) Doublet score evaluated by scDblFinder's computeDoubletDensity function for all mapped cells per cluster. Cells are colored by their doublet estimation determined by scDblFinder's doubletThresholding function. (**D**) Doublet estimation from 10x Chromium nephrogenic zone datasets. (**E**) Doublet estimation from 10x Xenium datasets. (**F**) Marker gene dot plot with size representing percentage of expressing cells in each cluster and color representing average scaled expression within each cluster. Marker genes for ureteric bud (*RET*, *GATA3*, *EPCAM*, *CDH1*, *CDH3*), mature nephron (*GATA3*, *CDH1*, *CDH3*, *PAX2*, *PAX8*), nascent nephron (*CDH6*, *WNT4*, *LHX1*, *JAG1*, *PAX8*, *PAX2*), nephron progenitor (*PAX2*, *SIX2*, *CITED1*, *EYA1*), stroma (*FOXD1*, *PDGFRA*, *PDGFRB*, *MEIS1*), endothelium (*EGFL7*, *KDR*), and immune (*CX3CR1*, *IFNGR1*). (**G**) Table showing top differentially expressed genes for cluster from refined cluster assignments. Column 1 shows cluster number, column 2 shows annotated cluster identity, and column 3 are top DEGs.

#### Supplementary Tables

**Table S1: Xenium experiment statistics.**

| sample, slide # as in GEO record: | ms E15, 1 | ms E15, 2 | ms E17, 1 | ms E17, 2 | ms E18, 1 | ms E18, 2 | hu wk 17, 1 | hu wk 17, 2 | hu wk 20, 1 | hu wk 20, 2 | |
|---|---|---|---|---|---|---|---|---|---|---|---|
| slide ID | 0040294 | 0040559 | 0040294 | 0040559 | 0040294 | 0040559 | 0049058 | 0049060 | 0049058 | 0049060 | |
| slide # as in analysis_summary.html | 2 | 1 | 2 | 1 | 2 | 1 | 2 | 1 | 2 | 1 | **row sum** |
| # kidneys | 6 | 9 | 10 | 5 | 4 | 4 | 1 | | 1 | | 40 |
| # tissue slices | 12 | 18 | 15 | 5 | 7 | 12 | 21 | 8 | 3 | 8 | 109 |
| # tissue slices with seg. niches | 5 | 4 | 4 | 5 | 2 | 3 | - | 3 | 1 | - | 27 |
| # niches segmented | 102 | 67 | 303 | 194 | 171 | 294 | - | 57 | 152 | - | 1340 |
| # depth-filtered niches | 39 | 46 | 204 | 81 | 106 | 91 | - | 57 | 120 | - | 744 |
| # cells detected | 172,225 | 244,107 | 329,957 | 111,807 | 304,788 | 328,589 | 338,916 | 288,214 | 295,662 | 433,710 | 2,847,975 |
| # transcripts detected* | 239,811,289 | 320,147,754 | 345,981,919 | 110,489,754 | 352,403,784 | 357,055,486 | 195,177,950 | 302,095,575 | 290,687,652 | 372,131,550 | 2,885,982,713 |
| % high quality transcripts | 83.4 | 85.6 | 86.8 | 87 | 84.3 | 83.9 | 86.1 | 81.7 | 81.8 | 83.8 | **row average** |
| median genes per cell | 762 | 714 | 616 | 590 | 675 | 647 | 341 | 637 | 611 | 534 | 613 |
| median transcripts per cell | 1,223 | 1,132 | 904 | 854 | 1,022 | 971 | 423 | 912 | 869 | 732 | 904 |
| est. false positive transcripts/cell** | 1.9 | 2.0 | 1.9 | 3.2 | 1.5 | 2.0 | 1.8 | 3.5 | 2.2 | 1.9 | 2.2 |

* high quality
** including genomic counts

**Table S2: Geometric features used to generate branching continuum.**

| feature index (ureteric bud) | feature index (nephron progenitor) | feature name |
|---|---|---|
| 0 | 55 | Area |
| 1 | 56 | Perimeter |
| 2 | 57 | Compactness |
| 3 | 58 | Convex Hull Perimeter |
| 4 | 59 | Convex Hull Area |
| 5 | 60 | Convexity |
| 6 | 61 | Concavity |
| 7 | 62 | Solidity |
| 8 | 63 | Minimum bounding rectangle area |
| 9 | 64 | Minimum bounding rectangle perimeter |
| 10 | 65 | Rectangularity |
| 11 | 66 | Mean ROI edge length |
| 12 | 67 | STD ROI edge length |
| 13 | 68 | Skewness ROI edge length |
| 14 | 69 | Kurtosis ROI edge length |
| 15 | 70 | Mean ROI edge angles |
| 16 | 71 | STD ROI edge angles |
| 17 | 72 | Skewness ROI edge angles |
| 18 | 73 | Kurtosis ROI edge angles |
| 19 | 74 | Mean ROI curvature |
| 20 | 75 | STD ROI curvature |
| 21 | 76 | Skewness ROI curvature |
| 22 | 77 | Kurtosis ROI curvature |
| 23 | 78 | Best fit ellipse center (X) |
| 24 | 79 | Best fit ellipse center (Y) |
| 25 | 80 | Best fit ellipse major axis length |
| 26 | 81 | Best fit ellipse minor axis length |
| 27 | 82 | Best fit ellipse eccentric angle |
| 28 | 83 | Best fit ellipse eccentricity |
| 29-38 | 84-93 | $1^{st}$-$10^{th}$ Zernike moments |
| 39-48 | 94-103 | $1^{st}$-$10^{th}$ Fourier coefficients |
| 49-54 | 104-109 | $1^{st}$-$6^{th}$ Hu moments |

**Table S3: Curated gene list for radial average *z*-score plots.**

| tissue | cell type | cell state | curated genes |
|---|---|---|---|
| **hu week 20** | nephron progenitors | naive | *TCF4, MEOX1, NBL1, ABTB2, ROBO2, TRIL, CHRNA1, PCDH18, TCF7L2, NPTX2, PCDH15, ELAVL4, CCDC80, TTC28, FOXD1, TMEM100* |
| | | differentiation | *ID3, ITPR1, NOTCH2, GXYLT2, PAX2, ID1, RXRA, DNM1, LYPD1, PAX8* |
| | | proliferation | *HIST1H1A, TYMS, CENPU, TOP2A, HIST1H4C, H2AFX, FEN1, MYBL2, PCLAF, ZWINT, MCM7, MCM6, HELLS, MCM4, PCNA, MCM2, CHEK1, CENPK* |
| | stroma | shallow | *SMOC2, EBF2, SFRP1, CDCA7L, MPPED2, PDE5A, NTN1, TGFBI, DNAH11, NDNF, SULF2, FOXD1, CYP1B1, TRIL* |
| | | deep | *IRS4, DKK1, GATA6, THBS1, CXCL12, ID2, ID3, MAN1A1, TNC, GBP1, EPHA7* |
| **ms E17** | nephron progenitors | naive | *Traf1, Cited1, Tril, Gas1, Mest, Meox1, Tpcn1, Ttc28, Ppp1r16b, Sost, Btbd11* |
| | | differentiation | *Dlc1, Ptpn13, H19, Bcam, Nr2f1, Gja1, Slit1, Id2, Scube3* |
| | | proliferation | *Pttg1, Kpna2, Racgap1, Fam83d, Ccnb1, Cdc20, Cenpa, Knstrn, Bub1b, Hist1h4h, Foxm1, Rrm2, Hist2h2bb, Hist1h2ap, Mki67, Top2a* |
| | stroma | shallow | *Ntn1, Arap1, Nlgn2, Adgrl2, Cldn1, Igfbp5, Plppr3* |
| | | deep | *Shisa3, Pcna, Pdgfrb, Rrm2, Bmper* |

**Table S4: Primer sequences used for organoid qPCR analysis.**

| gene | forward primer | reverse primer |
|---|---|---|
| **CRABP2** | CCTAGGAGTCTACGGGGACC | ATCACATTCACCCCCAGCAC |
| **CTGF** | CGCACAAGGGCCTATTCTGT | GAGCACCATCTTTGGCGGT |
| **CYP26A1** | GCTGCGATCAAGCTCTGGG | GAACTTCCTCCGCTGCAGTA |
| **CYR61** | GATTCGATGCCTCCGAGGTG | TCCATTCCAAAAACAGGGAGC |
| **GAPDH** | AGGTCGGAGTCAACGGATTT | TGGAATTTGCCATGGGTGGA |
| **HPRT** | TGCTTTCCTTGGTCAGGCAG | TTCAAATCCAACAAAGTCTGGC |
| **RARB** | ATCGGCACACTGCTCAATCAAT | GGTTTGTACACTCGAGGGGG |

#### Supplementary Movies

**Movie S1: Nascent nephron annotation in 3D-cultured wild-type mouse embryonic kidney explants via EpCAM staining.** Representative confocal fluorescence average projection timelapse of a FITC-labeled anti-EpCAM*-stained wild-type CD1 mouse embryonic kidney in 3D hydrogel culture (see **Methods**). *Inset:* Detail of nascent nephron formation local to a branching ureteric bud tip. White arrows indicate position and timing of nephron detection based on manual annotation.

**Movie S2: Nascent nephron annotation in 3D-cultured** ***Wt1***$^{\mathbf{EGFPCre/+}}$ **mouse embryonic kidney.** Representative confocal fluorescence timelapse of an AF660-labeled anti-ECAD-stained *Wt1*$^{EGFPCre/+}$ mouse embryonic kidney, following a single plane in 3D hydrogel culture (ECAD, green; EGFP *Wt1* reporter, magenta; see **Methods**). Frames were registered in *z* among timepoints in the original stack to follow an example nephron condensation event. *Inset:* Detail of nascent nephron formation local to a branching ureteric bud tip. White arrows indicate position and timing of nephron detection based on manual annotation.

**Movie S3: Schematic of nephrogenic niche branching life-cycle with cell state and mechanical rhythms.** (**A**) Schematic of the mouse ureteric bud branching life-cycle, cap mesenchyme dynamics, nephron formation, and distal nephron anastomosis with the ureteric bud tip, following the reference frame of a representative nephrogenic niche. (**B**) Model for spatial distribution of stromal and nephron progenitor cell transcriptional states. (**C**) Model for spatial distribution of mechanical stresses inferred from this work and ref. [38]. Stresses local to the ureteric bud branch point are speculative.

**Movie S4: Live explant timelapse for validation of branching continuum score and stromal dynamics.** Confocal fluorescence timelapse of an E11.5 Foxd1GC/+; R26mTmG/+ mouse kidney explant imaged between 24 and 72 hr of transwell culture. Orange cells are derived from Foxd1+ stroma; blue cells primarily belong to ureteric bud, cap mesenchyme (nephron progenitor), and early nephron compartments.

**Movie S5: Branching and nephrogenesis dynamics in AAV2-transduced mouse embryonic kidneys.** Representative anti-EpCAM*-labeled E13.5 CD1 wild-type mouse embryonic kidneys infected with AAV2-mCh-shScramble (top row), AAV2-mCh-shRock1/2 (middle row), or AAV2-mCh-shMylk (bottom row). AAV2 targets cortical nephrogenic zone stromal cells with the indicated shRNA. For each row, the views are: *Left*, Anti-EpCAM* and mCh composite confocal timelapses at a single plane lying at a depth of ~25% of the kidney diameter. *Middle*, The same timelapses isolating the mCh channel (showing cortical to medullar stromal infiltration). *Right*, Average *z* projection timelapse across the full imaged depth of the anti-EpCAM* channel. Anti-EpCAM* stains the ureteric bud and nascent pretubular aggregates/distal tubules/connecting segments. White arrows indicate connecting site/tubule formation superficial to the normal location.
